# Estimating single cell clonal dynamics in human blood using coalescent theory

**DOI:** 10.1101/2023.02.24.529817

**Authors:** Brian Johnson, Yubo Shuai, Jason Schweinsberg, Kit Curtius

## Abstract

While evolutionary approaches to medicine show promise, measuring evolution itself is difficult due to experimental constraints and the dynamic nature of body systems. In cancer evolution, continuous observation of clonal architecture is impossible, and longitudinal samples from multiple timepoints are rare. Increasingly available DNA sequencing datasets at single cell resolution enable the reconstruction of past evolution using mutational history, allowing for a better understanding of dynamics prior to detectable disease. We derive methods based on coalescent theory for estimating the net growth rate of clones from either reconstructed phylogenies or the number of shared mutations. Using single-cell datasets from blood, we apply and validate our analytical methods for estimating the net growth rate of hematopoietic clones, eliminating the need for complex simulations. We show that our estimates may have broad applications to improve mechanistic understanding and prognostic ability. Compared to clones with a single or unknown driver mutation, clones with multiple drivers have significantly increased growth rates (median 0.94 vs. 0.25 per year; p = 1.6 × 10^-6^). Further, stratifying patients with a myeloproliferative neoplasm (MPN) by the growth rate of their fittest clone shows that higher growth rates are associated with shorter time from clone initiation to MPN diagnosis (median 13.9 vs. 26.4 months; p = 0.0026).

## Introduction

Clonal expansions of cells that acquire certain mutations post-conception are a direct result of somatic evolution and are prevalent across the human body^1–7^. By estimating the timing of clone initiation and subsequent growth rates of clones, we can characterize evolutionary mechanisms that underlie aging^8^ and malignant progression^9–12^. In blood, for example, this evolutionary process is known as *clonal hematopoiesis* and has been associated with many aging-related disorders such as anemia^13^, impaired immunity^14,15^, and cardiovascular disease^16,17^, as well as progression to hematopoietic cancers^5,9,18^. Previous analyses found that somatic mutations conferring higher fitness, measured by clonal growth rate, lead to a higher risk of malignant transformation^19–21^. However, validated methods for measuring these important evolutionary parameters, which can vary from patient to patient, remain limited. Fast, accurate estimates of the underlying clonal dynamics using genomic data could serve to improve prognostic ability and ultimately lead to better patient outcomes.

Recent whole genome single-cell sequencing experiments in blood^19,22–24^ allow for phylogenetic reconstruction of the ancestral relationships between cells. Information on the growth dynamics of individual clones is contained in the phylogeny of sampled cells from a population^25^. Classical phylodynamics approaches to estimate population size trajectories depend on Kingman’s coalescent and its subsequent generalization to variable population size^26–28^. The coalescent, with the assumption of an underlying model such as logistic growth, provides the basis for clone growth rate estimation using the *phylodyn* R package^19,23,29^, as well as a method called Phylofit^22,24^. Alternatively, the package BEAST 2^30^ enables phylodynamic inference either by using the coalescent method or by modeling the population as a birth-death process, which allows the population size to vary stochastically without relying on coalescent approximations^31,32^. Due to the lack of an analytical solution for confidence intervals, these previous approaches estimate the growth rate using Markov chain Monte Carlo (MCMC), Integrated nested Laplace approximations (INLA), or Approximate Bayesian computation (ABC). Here we introduce new methods for estimating the net growth rate of a continuous time, supercritical birth-death process. The birth-death process is consistent with a cellular model of symmetric division (birth) and death or differentiation, and our methods remain valid for other models of clonal expansion that begin with an exponential growth phase (e.g., logistic, Gompertzian, etc.) in the context of hematopoietic data. Importantly, our methods require few assumptions and do not depend on computationally expensive simulation, allowing for near instantaneous estimation of the growth rate and its confidence intervals.

Our methods build on the mathematical work of Harris et al.^33^ and Lambert^34^, who recently discovered a relatively simple way to describe the *exact* genealogy of a sample of size *n* at time *T* from a birthdeath process. Using Lambert’s construction, we derive an approximation to the genealogy when *T* and *n* are large and use this approximation to obtain a maximum likelihood estimate of the net growth rate of a clone. We prove a limit theorem which gives the asymptotic distribution for the total lengths of the internal and external branches in the phylogenetic tree. The asymptotic distribution of the total internal branch length leads to a second method for estimating the net growth rate. This also allows us to estimate the net growth rate directly from the number of internal or shared mutations, those which are inherited by more than one of the sampled cells. Additionally, we provide an estimate for clone age which is applicable when the growth rate is known and the mutation rate is unknown.

Recent single-cell sequencing datasets ^19,22–24^ have generated novel insights about the nature of clones in the blood, identifying high risk mutations and revealing that clonal expansions with known drivers are present decades before symptoms appear^22^. Applying our methods to these datasets generates additional insights on the early growth rates of clones, which appear to be clinically relevant. We validate our estimates with longitudinal data and show that our methods contribute to a better understanding of the overall trajectory of the population size of a clone, refining previous estimates and further advancing our understanding of hematopoiesis, aging and cancer initiation.

## Results

We derived new mathematical estimates for evolutionary parameters (e.g., growth rate of a clone) when analyzing single cell-derived DNA sequencing data from a *sample* of the clone. A sample is a random subset of the total cells in the clone, as is commonly available in a realistic single-cell dataset. In the blood datasets analyzed below, samples of unique clones range from 4-109 cells^19,22–24^, while total clone size can hypothetically be as large as the total number of hematopoietic stem cells (HSCs) in the human body (estimates range from 25,000-300,000^20,35,36^). Analysis of coalescence times then requires explicit consideration of the size of the sample, and new theoretical results were needed to obtain analytical growth rate estimates in this setting. In this section, we provide our estimates for clonal growth parameters under a wide range of applicable modeling assumptions, then apply them to simulated and real data. We also compare our results to those produced using Phylofit, a recent coalescent-based MCMC approach^22^, and a birth-death MCMC approach introduced by Stadler^31^.

### Mathematical models for estimating clonal growth

First, we describe the biological rationale for inferring the growth rate from a genealogical tree. All cells sampled from the same clone progeny will have a common ancestor dating back to the clone’s origin, i.e., when the first cell acquired the identifying mutation leading to clonal expansion. Any two sampled cells may have a more recent common ancestor, and the most recent time at which the two cells have a common ancestor is called the *coalescence time* for these cells. In a sample of *n* cells, there will be *n* – 1 distinct coalescence times. For larger populations, it is less likely that any two sampled cells will have a recent common ancestor. Therefore, a faster growing (larger) population should have older common ancestors and a slower growing (smaller) population should have more recent common ancestors. Because the probability of a shared ancestor is dependent on the total clone size, the distribution of coalescence times provides information on the clone size trajectory, and here we use it to infer the early growth rate of the clone, *r*.

To connect growth rates to a genealogical tree, we consider the following birth-death process. We assume each cell divides symmetrically at rate *λ* and dies or differentiates at rate *μ*, acquiring mutations through time at rate *ν*. We wish to estimate *r* = *λ* – *μ*, the net growth rate (see Figure 1A). The data consists of a sample of *n* cells from the clone (of total size = *N* cells) at a clone age *T*. We assume that *r* is positive and constant during the expansion phase of the clone. We also assume the sample size *n* is much smaller than the total clone size *N*, which is usually valid for single cell-derived datasets described herein that typically have most coalescence events occurring shortly after clone initiation (i.e., star-shaped genealogies) ^19,22–24^. Although our mathematical results are proved for this simple birth-death process, they can be applied to a larger class of models that describe the expansion of clones in blood with an early exponential growth phase (e.g., logistic growth^19,22^, purely exponential growth^20^, Wright-Fisher with selection^23^, etc.). Our results will be valid when observed coalescence times are not impacted by changing growth rates that may occur after the initial expansion phase (see Figure 1B), and when the time at sampling, *T*, does not bias the coalescence times.

**Figure 1:**
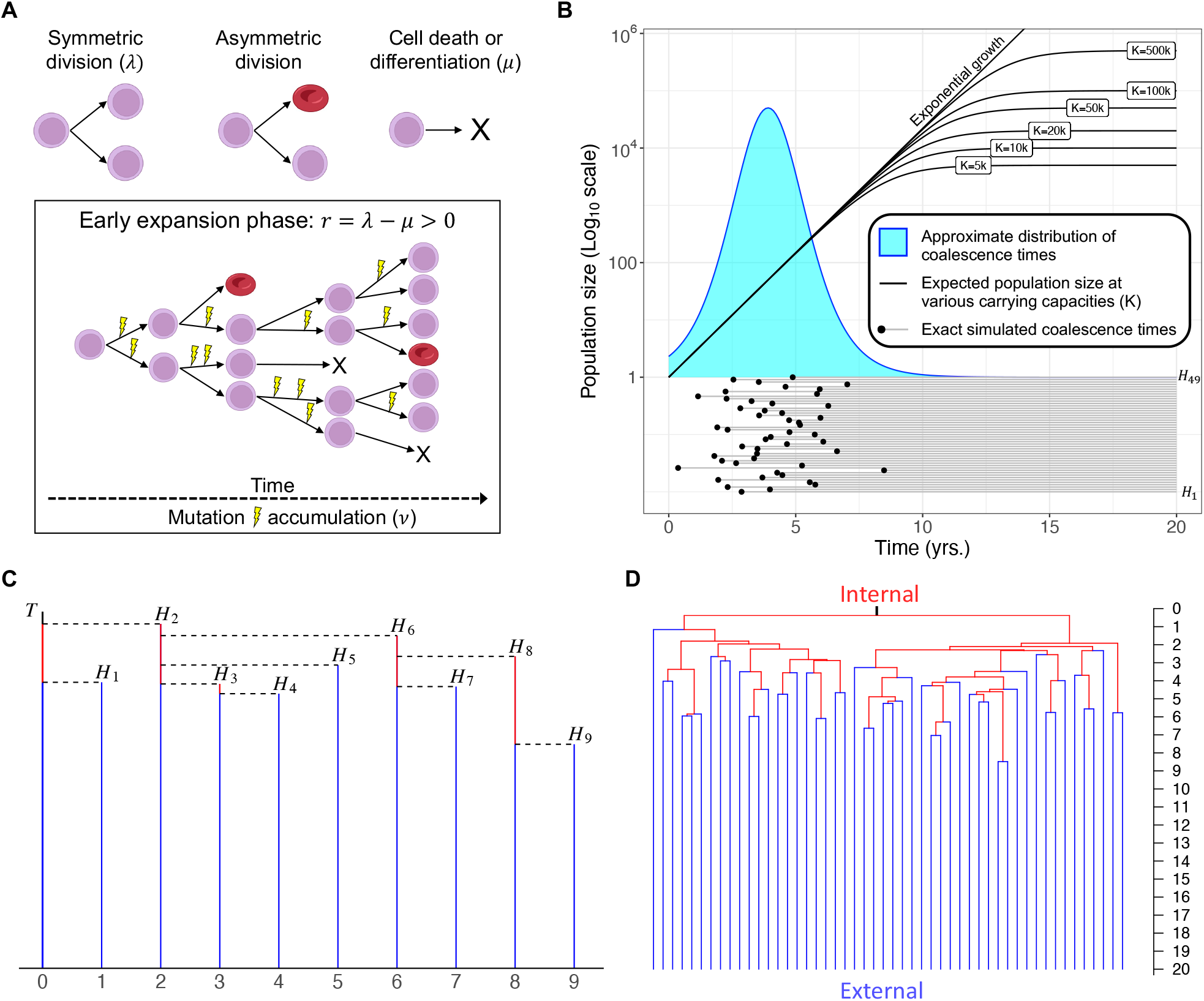
Model schematic and coalescent results. **A:** Stem cells undergo symmetric division at rate *λ*, increasing the population of stem cells by 1. Asymmetric division does not affect the population size or phylogeny except to introduce mutations. Cell death (or differentiation) occurs at rate *μ*, which removes the cell’s inherited history from the phylogeny and decreases the population size by 1. Our methods seek only to estimate the growth rate during the expansion phase of a clone, when the rate of symmetric division is greater than the rate of cell death (*r* = *λ* – *μ* > 0) and both rates are assumed to be constant during this phase. Mutations, which can occur at or between divisions, are assumed to accumulate linearly with time at rate *ν*. **B:** The approximate distribution of coalescence times for *n* = 50 cells is plotted above one example of *n*–1 = 49 coalescence times drawn from the exact distribution of coalescence times for a birth-death process. The expected population size assuming logistic growth with different carrying capacities shows that most coalescence events occur at smaller population sizes, when the growth trajectory is still approximately exponential. Other parameters: *r* = 1, (*λ* =1.5, *μ* = 0.5), *T* = 20. Note that a sampling time *T* < 10 years would artificially affect the distribution of coalescence times, introducing bias. **C:** Method overview: Reconstruction of a genealogical tree using the coalescent point process (CPP) can be done by first adding a vertical line of length T, and then adding successive vertical lines representing the coalescence times (*H_i_*). The coalescence times are drawn i.i.d. from the distribution defined in **Simulating the exact genealogy** and are then connected via horizontal lines to form the ultrametric tree. **D:** Tree reconstructed by randomly merging lineages with coalescence times from (B), which is statistically equivalent to using the coalescent point process. Red edges are internal, representing shared mutational history, and blue edges are external.

### Approximating genealogy using a coalescent point process

A recent elegant method by Lambert for computing the exact genealogy of a sample of size *n* at time *T* from a birth-death process is described in Simulating the exact genealogy^34^. Because we are mostly interested in the case when the clone age at sampling *T* and the sample size *n* are large, we can obtain a useful approximation by letting *T* →∞ and then *n* →∞ in Lambert’s construction. This leads to the following simpler method for approximating the coalescence times *H*_1_,…, *H*_n-1_, which provides the foundation for our estimates of the net growth rate:

1. Let *W* have an exponential distribution with mean 1.
2. Let *U*_1_, *U*_2_, …, *U*_*n*-1_ be i.i.d. random variables having the logistic distribution, which is a symmetric distribution on the real line with density given by

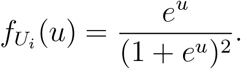
3. Let

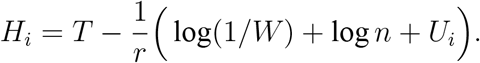

For mathematical details of the derivation, see Supplementary section 1.1. Note that we model the coalescence times as independent and identically distributed (i.i.d.) random variables having a logistic distribution, plus a random shift which accounts for the randomness in the initial growth of the branching process. Ignatieva, Hein, and Jenkins^37^ showed using a different method involving a random time change that the coalescence times can be approximated well by i.i.d. logistic random variables.

Once the *H_i_* have been determined, we can construct the genealogical tree by randomly merging two lineages at each coalescence time, or by using the coalescent point process as shown in Figure 1C. To understand the formula for *H_i_*, note that in a supercritical branching process (*r* > 0), the individuals sampled at time *T* are likely to have been descended from different ancestors fairly close to time zero, when the clone began expanding. That is, the genealogical tree will be nearly star-shaped, with most coalescence occurring near time 0. Note that the expected population size of the clone at time *t* is *e^rt^*. Considering the case when *λ* = *r* and *μ* = 0, the size of the population after a large time *t* can therefore be approximated by *We^rt^*, where *W* has an exponential distribution with mean 1. This expression equals *n* when 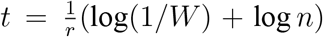. We expect most lineages to coalesce when the size of the population is comparable to *n*, which is why the coalescence times *H_i_* are close to 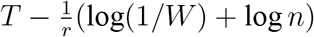.

### Estimating growth rate of a clone

If we can reconstruct the full genealogical tree from data, then we have estimates for the *n* – 1 coalescence times *H*_1_, …, *H*_*n*-1_. From the discussion above, we can write

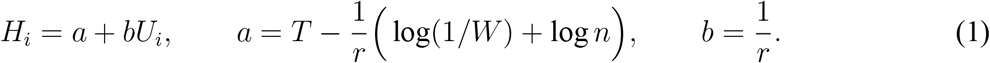

where the random variables *U*_1_, …, *U_n_* are i.i.d. and have a logistic distribution. Note that we can write *b* = 1/*r* instead of *b* = −1/*r* because the logistic distribution is symmetric. We can therefore estimate the growth rate *r* by estimating the parameter *b*. We introduce here three methods. Alongside the results below, we also created an *r* package *cloneRate* for implementing growth rate estimation on novel user input data (see Data and code availability).

### Growth rate estimation using maximum likelihood

Maximum likelihood can be used to estimate *b* from *H*_1_, …, *H*_*n*-1_. Because the maximum likelihood estimate does not have a closed form expression, it must be found using numerical methods. We computed the maximum likelihood estimate 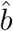 in R using the Nelder-Mead method^38^. From Equation 1, we can estimate *r* by

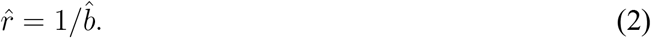

Let 0 < *α* < 1. A 100(1 – *α*)% confidence interval (CI) for *r* is

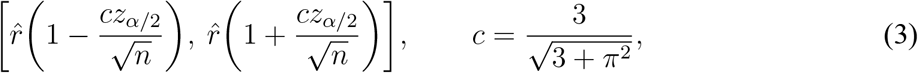

where *z*_*α*/2_ is the number such that if *Z* has a standard normal distribution, then *P*(*Z* > *z*_*α*/2_) = *α*/2. Note that *c* ≈ .836, which we use to compare to the confidence intervals of the following estimate based on internal branch lengths. See Supplementary section 1.2 for CI derivations.

### Growth rate estimation using internal branch lengths

If we are able to reconstruct the full tree, then we know the internal branch lengths 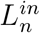 (e.g., sum of the lengths of red branches in Figure 1D). By Theorem 1 in Internal and external branch lengths in Methods, the distribution of 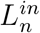 is approximately normal with mean *n*/*r* and variance *n*/r^2^. Therefore, we can estimate the growth rate by

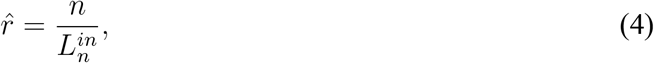

and we obtain an asymptotically valid 100(1 – *α*)% confidence interval for *r* by

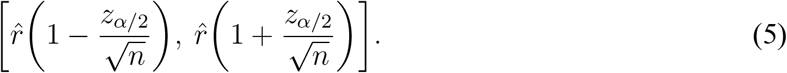

This estimate based on the internal branch lengths can be compared directly to the maximum likelihood estimate, as both methods take a time-based ultrametric tree as input. If the coalescence times are accurate, then considering only the internal branch lengths discards relevant information and one would expect the maximum likelihood estimate to perform better. The confidence bounds of the internal lengths estimate reflect this, as the confidence bounds of the internal lengths method in Equation 5 are identical in form to the confidence bounds for maximum likelihood in Equation 3, except that the internal lengths effectively has *c* =1. Because *c* =1 > 0.836, the internal lengths method has wider confidence intervals.

When reconstructing the tree from mutations, there will be some randomness inherent in the number of observed mutations. Because the previous methods use the time-based tree as input, neither accounts for this uncertainty. The following section uses the ideas presented here to estimate the net growth rate directly from the observed mutations, providing confidence bounds which account for the randomness of mutation accumulation.

### Growth rate estimation using shared mutations rather than full tree

If we can estimate the mutation rate *ν* during the expansion phase of the clone, then we can also estimate the growth rate directly from the number of shared mutations, defined as those mutations present in more than 1 but not all of the *n* sampled cells. The key idea is that there will be more shared mutations when the growth rate is smaller and fewer when the growth rate is larger. As shown in Shared and private mutations in Methods, the distribution of the number of shared mutations *M^in^* is approximately normal with mean *nν*/*r* and variance *σ*^2^ = *n*(*v*/*r* + *ν*^2^/*r*^2^). Therefore, if the mutation rate *ν* is known, we can estimate the growth rate by

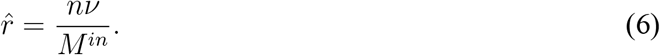

An asymptotically valid 100(1 – *α*)% confidence interval for *r* is given by

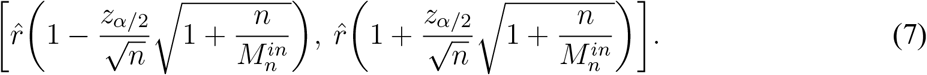

Accounting for Poissonian fluctuations in the observed number of shared mutations leads to confidence bounds slightly wider than those from the internal lengths method (Equation 5).

### Estimation performance on simulated data

To verify the performance of our methods, we generated trees using the exact genealogy reconstruction discussed in full detail in Simulating the exact genealogy in Methods. Recent work by Lambert^34^ allows for instantaneous generation of the exact genealogy of a sample from a supercritical birthdeath process, removing the need for complex simulations (such as those employed by the stochastic simulator MASTER^39^ in BEAST 2^30^) for many population genetics and coalescent applications. We briefly describe this process here and note that the tree generator is available in *cloneRate*. For a sample of size *n* at time *T* from a clone expanding with birth rate *λ* and death rate *μ*, *n* – 1 exact coalescence times are drawn using the process described in Simulating the exact genealogy. An example of a set of 49 coalescence times for a sample of 50 cells is shown in Figure 1B. Given the coalescence times *H_i_*,…, *H*_*n*-1_, the coalescent point process can be used to quickly reconstruct the genealogy. To reconstruct the tree from the coalescence times, we begin by drawing a vertical line of height *T*. We then draw vertical lines of heights *H*_1_,…, *H*_*n*-1_ and, at the top of each vertical line, draw a horizontal line to the left, stopping when it hits a vertical branch. The resulting tree is ultrametric, meaning that the root to tip distance is the same for all tips. Figure 1C shows a schematic example of generated a tree with 10 tips from 9 coalescence times, and Figure 1D shows the tree constructed using the coalescence times in Figure 1B.

Then, applying our methods to these reconstructed trees gives a distribution of estimates which allows for benchmarking since we know the ‘true’ growth rate in the simulated data. We compared the performance of our methods to the Markov chain Monte Carlo (MCMC)-based approach, Phylofit, introduced by Williams et al.^22^. We did not compare to the performance of their Approximate Bayesian computation (ABC)-based estimates, but note that the authors show a strong correlation between estimates from Phylofit and the ABC-based method (correlation coefficient *r* = 0.96)^22^. We also compared our methods to another MCMC approach based on the birth-death model using the likelihood given in Equation (5) by Stadler^31^. Whereas Phylofit is based on Kingman’s coalescent assuming logistic population growth, the method based on Stadler’s work models the population as a birth-death process, and assumes each individual is sampled with some fixed probability *ρ*. Our methods also use a birth-death process but instead assume a fixed sample size *n*, allowing us to obtain analytical approximations when the sample size is much smaller than the population size at the time of sampling.

### Performance across varying sample size *n* and growth rate *r*

We found similar performance using our methods compared to the MCMC methods. As shown in Figure 2A-B, both MCMC methods appear to perform slightly better for small sample size *n*, while our maximum likelihood method outperforms Phylofit for *n* ≥ 100. Of our two analytical methods, maximum likelihood has the lower root mean square error for larger values of *n* and converges to the birth-death MCMC as *n* becomes large (see Figure 2B). When the sample size *n* is too low, our approximation of the distribution of coalescence times, which is valid as *n* →∞, no longer accurately describes the population. Intuitively, a smaller sample provides less information available to make an accurate estimate of the growth rate. As such, performance deteriorates and the confidence intervals of our estimates expand with decreasing *n* (see Equations 3, 5, and 7). We use a cutoff sample size of *n* = 10 for each clone when applying to real data below, but note that this cutoff depends on the desired accuracy of the estimate.

**Figure 2:**
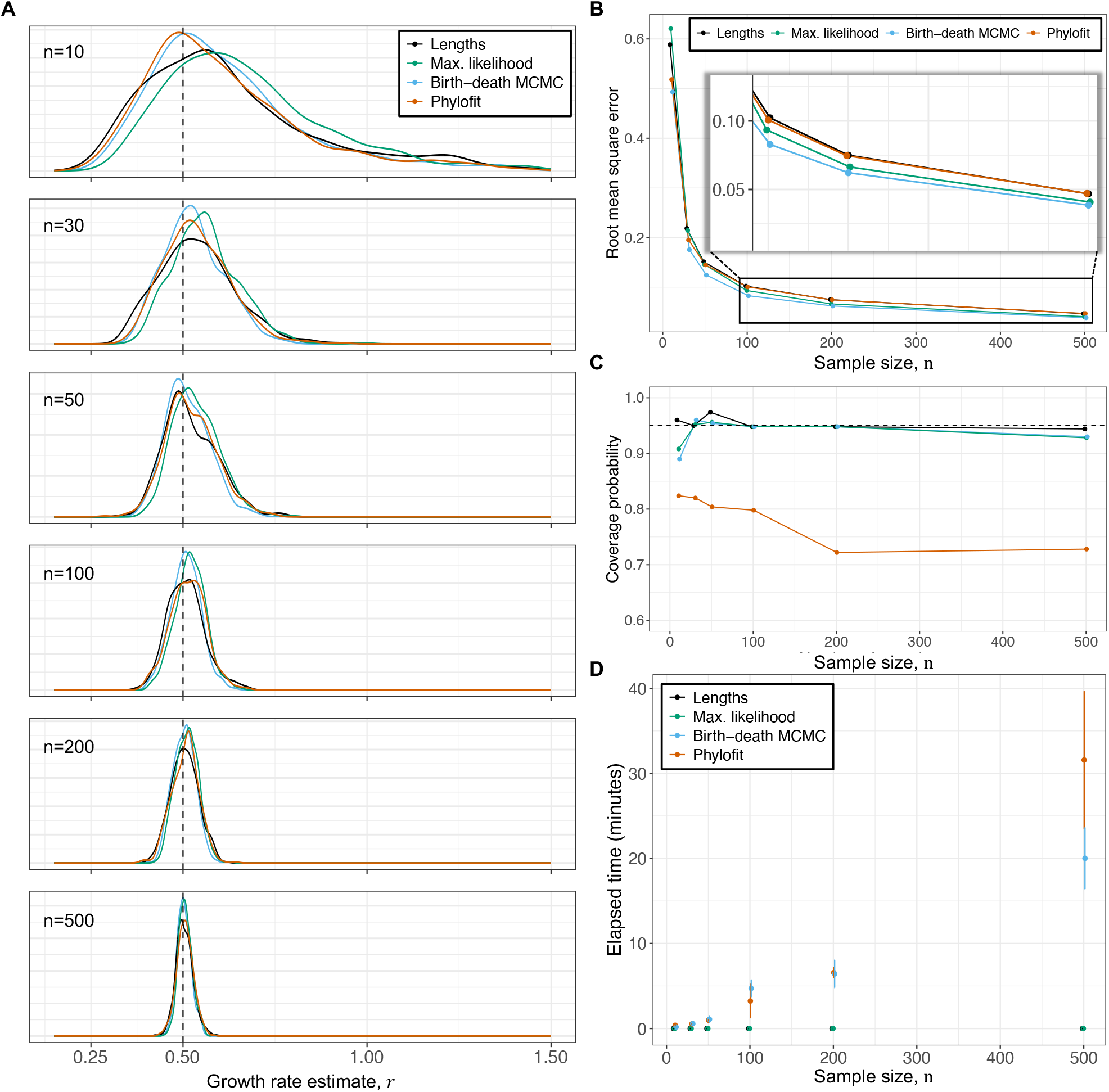
Performance and number of samples, *n*. **A:** Distribution of estimates from our methods using Equation 2 (green) and Equation 4 (black), Phylofit (orange), and the birth-death MCMC (blue) on 500 simulated ultrametric trees for each *n* value, where *n* is the number of sampled cells. Simulated trees were generated assuming a continuous time birthdeath branching process with *r* = 0.5 and *T* = 40. Birth rate *λ* was sampled from auniform distribution on [0.5, 1.5] and death rate *μ* = *λ* – *r*. **B:** Root mean square error for each method from simulated data shown in (A) illustrates improved performance with number of samples. MCMC methods are most accurate for small *n*, while the birth-death MCMC and maximum likelihood perform best for large *n*. **C:** Coverage of 95% confidence intervals methods based on simulations in (A). **D:** Runtime (mean +/- st. dev.) of various methods of estimating net growth rate shows that while the MCMC-based methods scale with the number of samples, *n*, our methods run effectively instantaneously for any tree size.

The confidence intervals for our methods are approximately accurate, as shown in Figure 2C, though we note that the maximum likelihood confidence intervals may be slightly too narrow for small *n* because the variance estimate is based on the asymptotic Cramer-Rao bound^40^. Alternatively, the 95% highest posterior densities (HPD) for Phylofit are consistently too narrow, where less than 80% of coverage is typically observed for growth rate estimates (Figure 2C). A similar observation about overconfidence of coalescent-based estimates has been made previously^32,41^, which is likely due to the fact that the birth-death process explicitly models stochastic population changes while the coalescent-based approach assumes that the change in population size is deterministic. Finally, we show that the MCMC methods’ runtimes scale with the number of samples, while our analytical methods are essentially instantaneous regardless of the size of the tree (Figure 2D).

Similarly, we quantified the performance of our methods across *r* values. As shown in Figure 3A-B, the four methods perform comparably well, with the birth-death MCMC having the lowest root mean square error. Again, Figure 3C shows that our confidence intervals are very accurate, especially when *r* ≥ 0.5, while coverage by Phylofit is below 80% when applied to the birth-death trees. In Figure 3, the smallest growth rate *r* = 0.25 with *T* = 40 shows concerning performance, which motivated further investigation into the impact of small growth rates.

**Figure 3:**
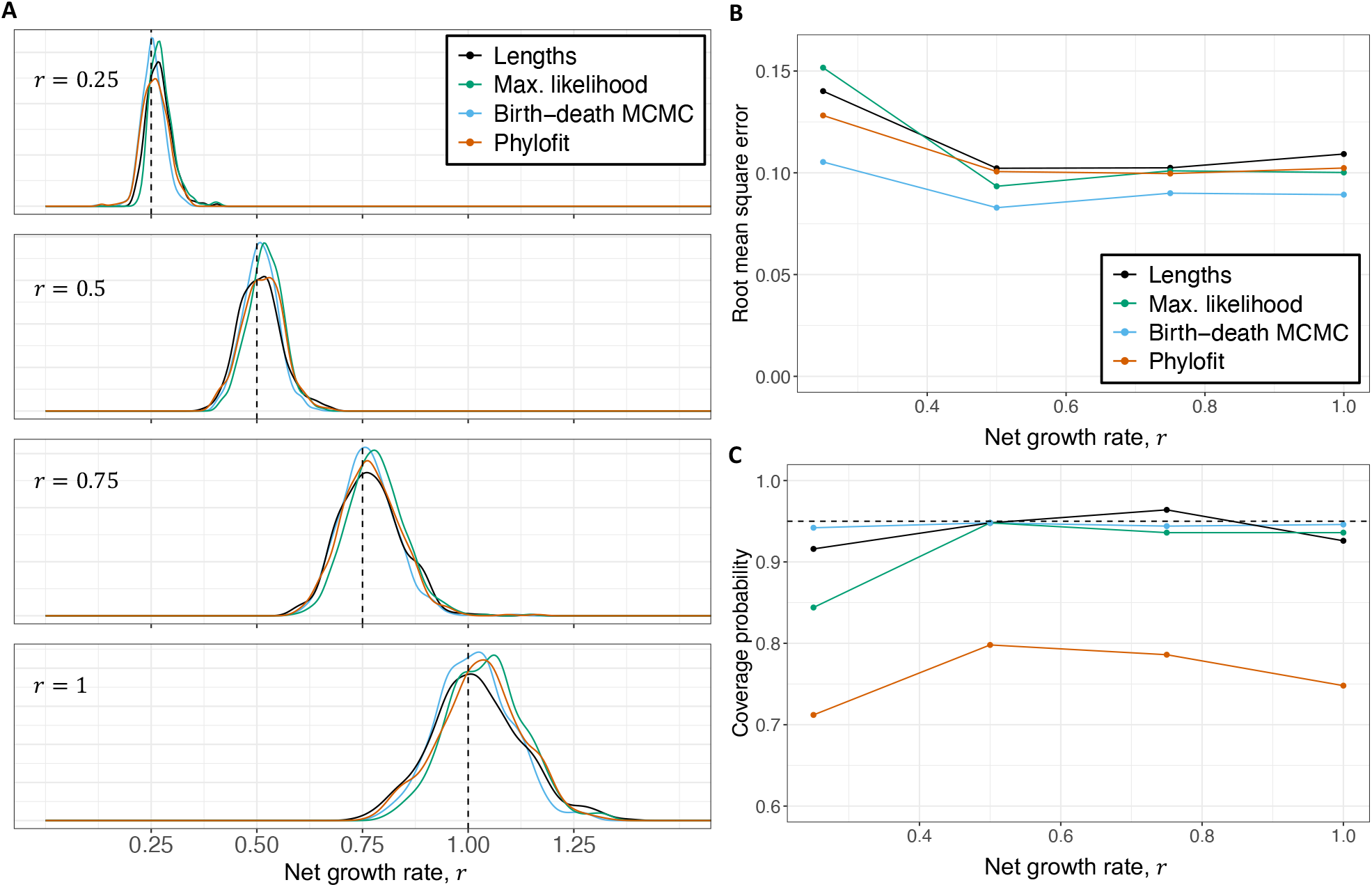
Performance across different growth rates, *r*. **A:** Distribution of estimates from our methods, Phylofit, and the birth-death MCMC on 500 simulated ultrametric trees for each *r* value, where *r* is the growth rate. Simulated trees were generated assuming a continuous time birth-death branching process with *n* = 100 and *T* = 40. Birth rate *λ* was sampled from a uniform distribution on [*r*, 1 + *r*] and death rate *μ* = *λ* – *r*. **B:** Root mean square error, normalized to account for the different true growth rates, for each method calculated for simulated data shown in (A) illustrates a decrease in accuracy for small growth rates. Performance of MCMC methods are less affected by small growth rates. **C:** Accuracy of 95% confidence intervals of the methods using data from simulations in (A).

### Small growth rate diagnostic for method utilization

We investigated the performance failure at small growth rates and determined why this happens. We then derived a diagnostic to determine when the growth rate is large enough for our analytical methods to be applicable (Figure 4A). As shown in Figure 4B, our methods perform worse than the MCMC methods in the problematic small *r* regime. To understand why our analytical methods can fail for small *r*, we note that when *r* > 0, so that the branching process is supercritical, the *n* sampled cells should all have distinct ancestors that were alive a short time after the initiation of the clone. Consequently, the genealogical tree will be nearly star-shaped, with long external branches and short internal branches near the root (see example in Figure 1D). On the other hand, when *r* = 0, so that the branching process is critical, the population size is nearly stable over time. Then the genealogy of the *n* sampled cells would more closely resemble the classical Kingman’s coalescent, in which lineages merge at a constant rate and most coalescence events occur near the time of sampling, leading to long internal branches and short external branches. When *r* is small but positive, the genealogical tree will be star-shaped if the sampling time *T* is sufficiently large, but the internal branch lengths will still be longer than when *r* is larger. Consequently, if *T* is not sufficiently large, then the constraint that the coalescence times must be less than *T* will affect the distributions of both the coalescence times and the internal lengths, and the approximation that we derived will not be accurate.

**Figure 4:**
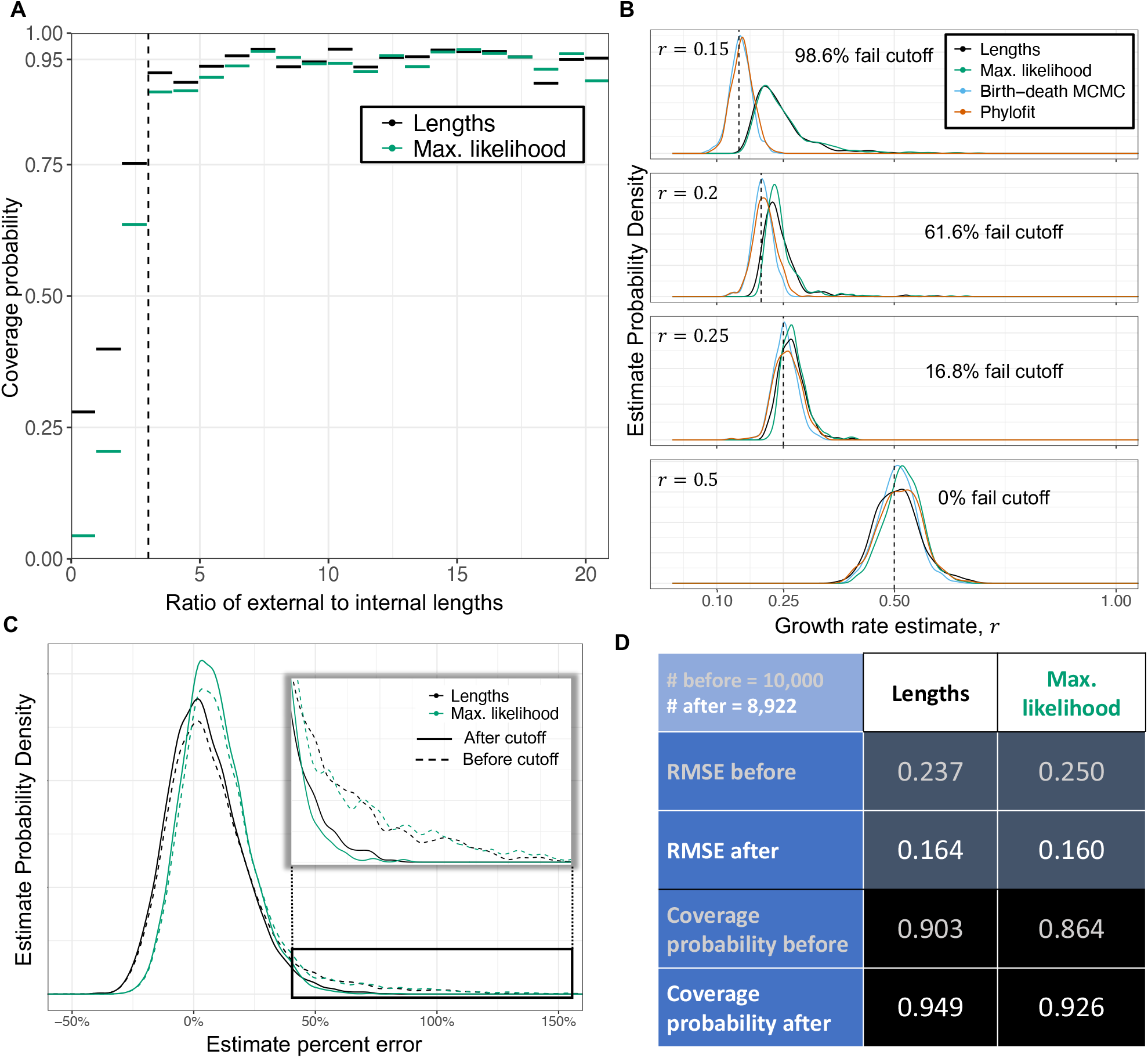
Small growth rate *r* diagnostic. **A:** Trees with *n* = 50 tips and *T* = 40 were created from 10,000 randomly sampled *r* values from a uniform distribution on (0.1, 1) and then binned by the ratio of external to internal lengths in increments of one. For corresponding estimates of *r*, accuracy of the 95% confidence intervals (CI) shows that the ratio of external to internal lengths can be used as a diagnostic to determine whether our method can accurately estimate growth rate. We use a ratio of 3 as a cutoff value as simulations with a ratio greater than 3 are captured by the 95% confidence intervals approximately 95% of the time. Minimum number of simulations captured in a single bin is 61. **B:** Distribution of estimates from our methods, Phylofit, and the birth-death MCMC on 500 simulated ultrametric trees from each *r* value, where *r* is the growth rate. Simulated trees generated assuming a continuous time birth-death branching process with *n* = 100 and *T* = 40. For each tree, we calculate the ratio of external lengths to internal lengths. For each growth rate, we show the percentage of trees which have a ratio of external to internal lengths below the diagnostic value of 3. **C:** Relative fractional error distribution for four methods from the same simulations shown in (A) before (dashed lines, iterations = 10,000) and after (solid lines, iterations = 8,922) the cutoff of 3 was applied. Inset shows significant reduction in overestimates due to the diagnostic cutoff. **D:** Normalized root mean square error (RMSE) and coverage of 95% confidence intervals for four methods using same simulations shown in (A) and (C) before and after the diagnostic cutoff was applied. The diagnostic provides a significant reduction in error and improvement in accuracy of confidence intervals.

We found that as long as the total length of the external branches is much larger than the total length of the internal branches, this problem does not arise, and our methods give accurate results. This leads naturally to a diagnostic which determines when our methods are applicable. As shown in Figure 4A, when the ratio of external to internal lengths is greater than or equal to 3, our methods and confidence intervals are accurate. Figure 4B shows that most of the simulated trees with problematic small growth rates fail this diagnostic cutoff. Applying this cutoff to a simulated dataset with growth rates between 0.1 and 1 reduces overestimates and greatly improves performance (Figure 4C-D). Notably, small growth rate clones at a relatively young clone age are unlikely to be observed in enough sampled cells in real data to make an accurate estimate; a requirement of *n* ≥ 10 cells is unlikely to be satisfied in these clones. In fact, none of the 42 clones which we analyze from the blood datasets below has an external to internal length ratio less than 4, and only two clones have a ratio less than 5.

### Application to human blood datasets

We applied our methods to single-cell derived sequencing data from human blood (Table 1). The methods for generating the data are fairly similar across the studies: single hematopoietic progenitor cells were clonally expanded and each single-cell derived colony was sequenced to a mean depth of roughly 15x, with slight differences depending on the study^19,22–24^. Time-based ultrametric trees generated in these studies are used as input for our methods, Phylofit, and the birth-death MCMC. Manual annotation is required to identify clonal expansions, associate clones with specific drivers, and to remove nested subclonal expansions from the clone of interest. We generally designated the clones as annotated in the studies which produced the data^19,22–24^ and provide details in Supplementary section 3.3.

**Table 1:** Whole genome sequencing datasets of single-cell derived colonies. Number of clones indicates the number of clonal expansions with *n* ≥ 10 cells sampled. As some clones profiled by Williams et al.^22^ had *n* ≥ 10 cells sampled at multiple timepoints from the same clone, we also specify the number of unique clones. See Supplementary section 3.3 for details on annotating clones.

| Number of Individuals | Number of Clones | Data source | Diagnosis | Ref. |
| --- | --- | --- | --- | --- |
| 11 | 18 (15 unique) | Adult peripheral blood (PB) and/or bone marrow (BM) | Myeloproliferative Neoplasm (MPN) | 22 |
| 2 | 2 | Adult BM | MPN | 23 |
| 3 | 15 | Adult PB and/or BM | Normal | 24 |
| 3 | 7 | Adult PB | Normal | 19 |
| **19** | **42 (39 unique)** | **Total** |  |  |

First, we check our assumption of neutrality within expanding clones (i.e., all cells within the clone grow at approximately the same rate). Previous authors have studied the expected site frequency spectrum for a sample from a birth-death process. Letting 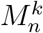 denote the number of mutations inherited by *k* of the *n* sampled individuals, Durrett^42^ showed that as *T* →∞, for *k* ≥ 2 we have

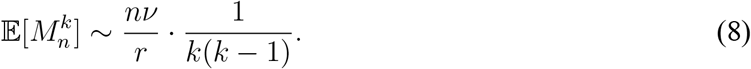

Gunnarsson, Leder, and Foo^43^ calculated the exact expectation in the case when the entire clone is sampled (see also^44,45^ for similar calculations). Therefore, we expect the site frequency spectrum to follow the curve 1/*k*(*k* – 1), where *k* equals the number of cells. In Figure 5A, we show the averaged site frequency spectrum across all clones, with any nested subclones removed, along with the 95% confidence interval of the mean. The agreement between the observed mean and the expectation indicates neutrality within clones, consistent with previous conclusions in blood^19,22–24^. For more detailed data on the site frequency spectrum for each clone, see Supp. Table 5.

**Figure 5:**
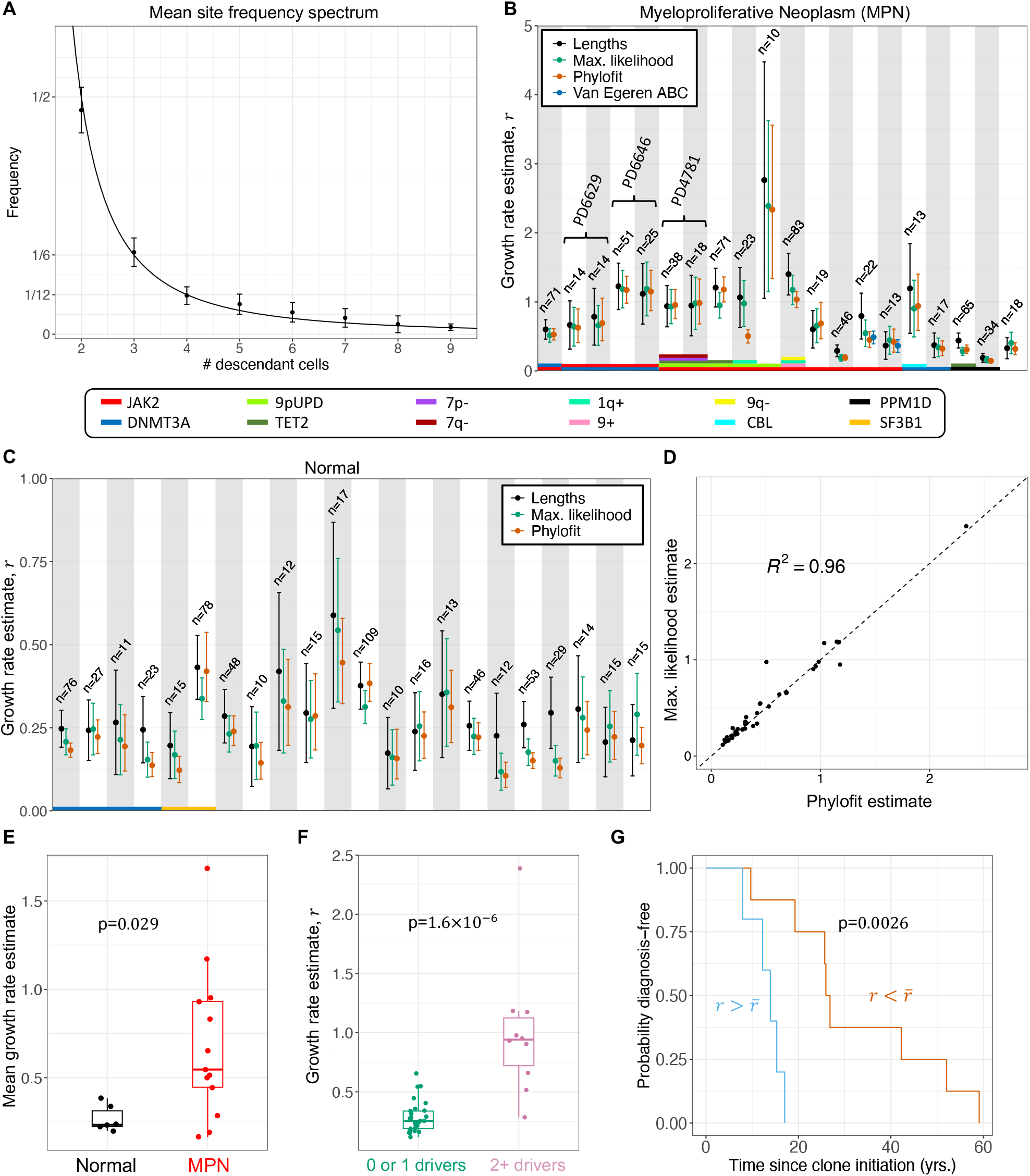
Applying estimates to blood data. **A:** Averaged site frequency spectrum across 42 clones shows agreement with the 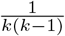 neutral expectation (solid line). Error bars show 95% confidence interval of the mean. **B-C:** Our estimates and Phylofit for clones with *n* ≥ 10 tips from individuals with (B) and without (C) Myeloproliferative neoplasms (MPN) shows good agreement across methods. Brackets in (B) group estimates from the same clones in the same patient estimated from two distinct samples taken years apart, showing consistency of estimates. Note that we also include estimates from Van Egeren et al.^23^ in dark blue for the two clones from their dataset. **D:** Correlation between our maximum likelihood estimate and estimates from Phylofit for all clones from (B) and (C). **E:** Mean maximum likelihood net growth rate estimate for clones from patients with (red) and without (black) a diagnosis of MPN shows that more aggressive expansions are associated with MPN. **F:** Maximum likelihood net growth rate estimate for clones with single or unknown drivers (green) and multiple drivers (magenta) show that fitness predicted by our methods is consistent with effects of known drivers. Non-parametric Mann-Whitney test used for p-value calculation in (E, F). **G:** In the single most aggressive clone from each patient diagnosed with MPN, stratification by mean net growth rate 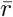 shows significant differences in Kaplan-Meier survival curves from clone initiation to MPN diagnosis (log-rank test p=0.0026) though sample set was small (13 patients). At time of sampling, mean age of high growth rate group was 60.3 years, median was 50.4 years. Mean age of low growth rate group was 60.9 years, median was 63 years.

In applying the methods to real data, we found agreement across our two analytical methods and agreement with the estimates from Phylofit (see Figure 5B-D). Close agreement was also found between our maximum likelihood method and the birth-death MCMC approach (*R*^2^ = 0.995, see Supp. Fig. 1). Estimates using the internal lengths method were slightly higher than maximum likelihood in some clones, and we expect that this is due to non-random merging of lineages as a result of slight fitness differences within the clone (see Supp. Fig. 2 for details). As discussed in Comparing analytical estimates to those using Phylofit in Methods, we only include the estimates from Phylofit without including the sampled clonal fraction as a target because clones have been shown to behave unpredictably at high clonal fractions, decelerating more than would be expected by a logistic growth traj ectory ^19,24^. Also, sampled clonal fraction and/or Variant Allele Frequency (VAF) may be a poor estimate of mutant allele burden in progenitors and HSCs (see Supplementary section 3.2), possibly due to lineage bias in mutated cells, such as the erythroid lineage bias observed in *JAK2* mutants^23^.

The most fit clones (those with fastest growth rates) were observed in patients with myeloproliferative neoplasms (MPN). As shown in Figure 5E, we found significantly increased estimates of mean detected clone fitness in individuals diagnosed with MPN as opposed to healthy adults (p=0.029). Additionally, Figure 5F shows that multiple-driver clones have significantly increased rates of expansion as compared to clones with just one or zero known driver mutations (p=1.6 × 10^-6^). This suggests increasing fitness effects from the accumulation of additional mutations. Higher growth rates may also be associated with shorter time from clone initiation to cancer diagnosis (log-rank p=0.0026), as shown in Kaplan-Meier curves in Figure 5G. Here the clone initiation time is estimated to occur 1/*r* years before the first coalescence (i.e., first surviving symmetric division). Together, these findings indicate that mechanistic rates for clonal dynamics such as the early growth rate may provide clinically important information in our understanding of hematopoietic stem and progenitor cell evolution and transformation to malignancy.

### Longitudinal validation of clone growth estimates

We leveraged available longitudinal data to validate our growth rate estimates. Again, for single cell-derived data, we used a lower bound of *n* = 10 cells per clone to include in our analysis. For longitudinal bulk data, we restrict analysis to expanding clones with a minimum of 4 timepoints available of the same bulk cell type. Further, because coalescent estimates are relevant for the early growth rate, we require that longitudinal data have at least two samples with a variant allele frequency between 0 and 0.25. The longitudinal data consists of peripheral (whole) blood samples^19^ and peripheral blood granulocyte samples^22^. It has been suggested that clonal fraction may differ across different blood cell types (granulocyte vs. whole vs. mononuclear)^23^. Data from Williams et al.^22^ is consistent with this finding, as there is significantly different clonal fraction across sampled cell type in 3 out of 4 patients where multiple cell types were sampled within a month of each other (see Supp. Fig. 5). Therefore, we require that a consistent type be used within each longitudinal growth rate estimate. For more details on the criteria for analysis, see Methods section Longitudinal clone inclusion criteria.

We analyzed 4 clones from Williams et al.^22^ and 56 clones from Fabre et al.^19^ that had appropriate longitudinal data. Of these 60 clones, 3 have sufficient matched data from single cell-derived WGS samples (1 clone from Williams et al.^22^ and 2 clones from Fabre et al.^19^) to allow for orthogonal estimates from the same clone. Results for these 3 clones are shown in Figure 6A-C, along with a logistic growth model fit. While we show the corresponding single cell colony clonal fraction (orange, divided by 2 to scale to the VAF of a diploid mutant), we do not use this data point in the fitting as the different cell type may affect the clonal fraction, as noted above. The logistic growth model is used primarily to identify the growth rate, *r*, and is chosen because it has been shown to face fewer parameter identifiability issues than other sigmoid growth models, such as Gompertz or Richards’, when applied to similar data^46^. For details on the longitudinal modeling, see Methods section Logistic modeling of longitudinal data.

**Figure 6:**
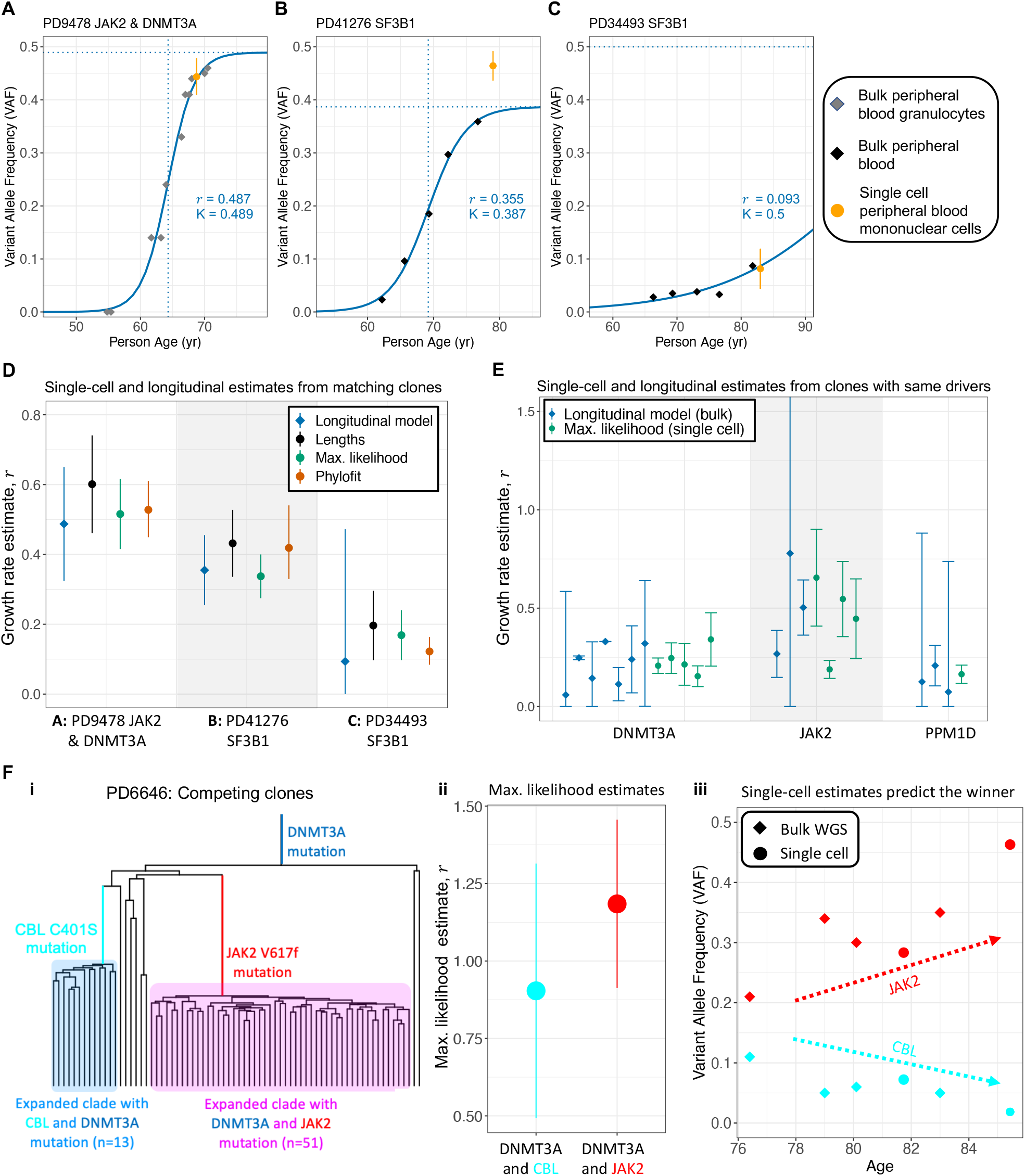
Longitudinal validation. **A-C:** Logistic fit to longitudinal data for three clones which have both single cell and longitudinal data. Only longitudinal bulk WGS data was used for fitting. Single cell colony clonal fraction (divided by 2) and 95% confidence intervals are shown in orange. Source for (A) is Williams et al.^22^, and source for (B) and (C) is Fabre et al.^19^. **D:** Longitudinal and single-cell estimates for each of the clones in (A-C) show agreement across data types. **E:** Longitudinal and single-cell estimates for different clones sharing the same driver. **F:** Clonal competition between a *DNMT3A+CBL* clone and a *DNMT3A+JAK2* clone shown in the reconstructed phylogeny (i). Maximum likelihood singlecell estimate from each clone (ii) shows that the *DNMT3A* + *JAK2* clone likely has higher fitness. Longitudinal data shows that the *DNMT3A+JAK2* clone increase in VAF over time while the *DNMT3A+CBL* clone decreases, confirming that the *DNMT3A+JAK2* clone has higher fitness, as predicted by our maximum likelihood estimate. All error bars represent 95% confidence intervals.

In comparing the growth rates from the longitudinal fits to our methods and Phylofit (Figure 6D), we found general agreement in the estimates, though we note the wide confidence intervals especially from the logistic model fit. Additional longitudinal data from Fabre et al.^19^, although not from clones with matched single cell data, was also used to compare to our coalescent estimates. First, we identify longitudinal clones with drivers also present in the single cell data. Then, we fit the logistic growth model to these longitudinal clones. After filtering (see Longitudinal clone inclusion criteria) and excluding the 3 clones shown in Figure 6A-C, there were 13 clones with longitudinal data and a driver gene also found in single cell clones. This data comes from clones with a mutation in one of the following genes: *DNMT3A, JAK2*, or *PPM1D*. The estimated growth rates are shown in Figure 6E. Similar growth rates in the same driver genes shows general consistency across all methods, though the small amount of data and wide confidence intervals limit the conclusions that can be drawn.

Finally, we considered competing clones within the same patient. If our estimates are relevant for clonal fitness, we would expect that clones with higher estimated growth rates should out-compete clones with lower estimated growth rates. The only example of competing clones with sufficient single cell data comes from patient PD6646 from Williams et al.^22^ (Figure 6F). A *CBL* and a *JAK2* mutation gave rise to two independent clones, both of which already have a *DNMT3A* mutation (Figure 6F i). By our maximum likelihood estimate, the *DNMT3A* + *JAK2* clone is slightly more fit than the *DNMT3A* + *CBL* clone (Figure 6F ii). Both Phylofit and our internal lengths method also estimate a higher growth rate for the *JAK2* clone. While this patient was undergoing treatment in this time period and the trajectory does not appear logistic, the *JAK2* clone increases in variant allele frequency while the *CBL* clone decreases, consistent with our estimate suggesting that the *JAK2* clone is more fit. There is an important caveat in this example because the specific interactions between clone/mutation and treatment may be responsible for the increasing/decreasing VAF, which is not accounted for by our estimate of fitness that characterizes early growth before treatment would have begun.

## Discussion

We developed new methods using coalescent theory to estimate rates of clonal expansion (and clone age) at greatly reduced computational expense. Leveraging previous work^34^, we validated our methods using efficient computational realizations of phylogenies resulting from birth-death branching processes. We then applied our methods to single cell resolution data from blood, showing that our growth rate estimates are both meaningful and consistent in biological and clinical contexts. We found general consistency of estimates with a previously published MCMC-based approach, Phylofit^22^ (R^2^ = 0.94-0.96), and a birth-death MCMC approach^31^ (R^2^ = 0.965-0.995). Where possible, we validated our estimates using single cell data from multiple timepoints, and also show that our estimates are consistent with and generally more precise than orthogonal estimates of net growth rate derived from longitudinal bulk data. Because they are based on analytical results, our methods for estimating growth rates from phylogenetic reconstruction are simple and run quickly without sacrificing accuracy. For future datasets with a higher number of sampled cells *n* and larger numbers of patients and clones, near instantaneous runtime at any tree size may be a critical feature separating our methods from MCMC or ABC-based alternatives. We provide a simple and easy to use R package, *cloneRate*, which will allow other researchers to estimate growth rates with their own input data.

For testing model performance on simulated data, we use results of Harris et al. ^33^ and Lambert^34^ to reconstruct the exact genealogy of a sample of size *n* from a birth-death process at time *T*, conditional on the population size being at least *n* at time *T*. This method avoids the need to simulate the entire large clonal population starting from a single cell as is commonly performed in other methods. From a mathematical point of view, the idea of using the coalescent point process to obtain results about statistics such as the site frequency spectrum and the allele frequency spectrum goes back to Lambert^47^ and was later developed further^48–50^, and then was recently applied to cancer modeling by Dinh et. al.^51^. Here we combine these ideas with the results from Lambert^34^ to obtain asymptotic results for quantities that can be derived from a large sample from a birth-death process. By taking advantage of the independence that is inherent in the coalescent point process, we are able to apply the *m*-dependent Central Limit Theorem to show that the total internal branch length, which can be used to estimate the growth rate of the process, has an asymptotic normal distribution. This observation allows us to obtain an asymptotically valid confidence interval for the growth rate; to our knowledge, ours is the first method for confidence interval construction that does not rely on Bayesian inference techniques. Finally, this is a unifying method for growth rate estimation that is applicable to many biologically relevant models assumed in previous works for clonal dynamics in blood^19,20,22–24^.

Acknowledging that this is both a limitation and a strength, our methods estimate only the growth rate in the early expansion phase, when growth is approximately exponential. Growth rates following the initial expansion phase may change over time in unpredictable ways and this, in most cases, should not affect our results. As such, our methods work well for clones with star-shaped trees that are common in blood and other somatic cell datasets. In the case of critical branching behaviour (i.e., trees with relatively longer internal branches), our simulations indicate that the MCMC approaches should be applied. In focusing only on the early growth rate, our methods do not rely on assumptions of the overall growth trajectory. Additionally, we have shown that early growth rates are relevant to the greater context of clonal and malignant hematopoiesis. For example, we found that higher growth rates are associated with shorter time from clone initiation to MPN diagnosis. The association between MPN diagnosis and growth rate suggests a possible avenue for early detection by predicting which patients are more likely to remain asymptomatic and which are more likely to undergo malignant transformation. Understanding the role of evolutionary dynamics to predict risk of progression in clonal hematopoiesis and provide prognostic information in hematological malignancies has been noted as a top clinical priority^52,53^.

Further, multi-driver clones show significantly increased rates of expansion, suggesting possible cumulative and/or synergistic effects of driver mutations. We found wide heterogeneity of fitness effects for *JAK2* clones, consistent with previous findings^19^, and relatively low fitness effects with smaller variation for *DNMT3A* clones. In the context of clonal hematopoiesis, single hit drivers with lower growth rates may increase risk for MPN by increasing the reservoir of cells at risk of additional stochastic mutations, thus initiating multi-hit driver clones with potentially additive fitness effects. There are other possible benefits to knowing the early rate of expansion. For example, early expansion rates are affected by fewer outside pressures such as treatment^54^ and may be more consistent across patients.

Our findings also provide guidance to experimental researchers designing single-cell DNA sequencing experiments that aim to determine clone fitness. The minimum number of sampled cells required for reliable estimates of growth rates falls roughly between 10 and 30, depending on desired accuracy (see Figure 2). Bulk whole genome sequencing performed prior to single-cell experiments could provide variant allele frequency information that can be used to estimate the cell fraction of clones of interest. Then, the total number of cells sequenced can be decided in a way that ensures enough sampled cells from clones of interest are included, while reducing overall costs.

One limitation is that current methods rely on the manual annotation of clones from a phylogenetic tree. While this is currently a fairly easy task given the relatively small size of single cell DNA sequencing datasets, it may become more challenging for expected increases in throughput^55^. An automated way to detect clonal expansions and distinguish normal cell turnover from expansions may be required to effectively scale the application of our methods. Such an automated algorithm would likely have to leverage not just the distribution of coalescence times, but also measures of tree balance.

Phylodynamics for human somatic data and cancer is a rapidly growing area and new tools are needed for useful applications^25,56,57^. With our methods, phylogenetic reconstruction can become an even more powerful tool to infer the past evolutionary dynamics of a population of cells. It has been hypothesized that individuals at high risk of developing myeloid malignancies can be identified before presenting with any symptoms^21^. Knowing which drivers are associated with more aggressive expansions will provide clinicians with better tools to direct treatment and/or prevention strategies. Additionally, clonal expansions without known drivers can provide mechanistic and biological insight. While blood is currently the most convenient medium for creation of single cell-derived DNA sequencing data and validation of these methods, age-related clonal expansions are also a feature of somatic evolution in tissues with spatial organization. Selection of the same drivers are found at similar burdens in solid tissues across patients, and thus accurate phylogenetic reconstruction in solid tissues may allow our method to be applicable in a variety of disease types. More comprehensive methods and datasets leading to the construction of more accurate phylogenetic trees^58,59^, when combined with the methods presented here, will enable researchers and clinicians to quickly draw conclusions about net growth rate from mutational data.

## Methods

### Simulating the exact genealogy

We present here Lambert’s construction of the exact genealogy of a sample of size *n* at time *T* from a birth-death process^34^. The idea is to describe the genealogical tree of *n* individuals from *n* – 1 random variables *H*_1_,…, *H*_*n*-1_ which represent coalescence times. To reconstruct the tree from the coalescence times, we begin by drawing a vertical line of height *T*. We then draw vertical lines of heights *H*_1_,…, *H*_*n*-1_ and, at the top of each vertical line, draw a horizontal line to the left, stopping when it hits a vertical branch. The resulting tree is ultrametric, meaning that the root to tip distance is the same for all tips. See figure 1C-D. This construction is known as a coalescent point process and goes back to the work of Popovic^60^ and of Aldous and Popovic^61^ in the setting of critical branching processes.

By building on earlier work of Stadler^31^ and Lambert and Stadler^62^, who considered the case in which each individual in the population is sampled with some fixed probability *y*, Lambert^34^ showed that we obtain exact the genealogy of a sample of size *n* from a birth-death process at time *T*, conditional on the population size at time *T* being at least *n*, if we choose *H*_1_,…, *H*_*n*-1_ by the following two-step procedure:

1. Choose a random variable *Y* with probability density function on (0,1) given by

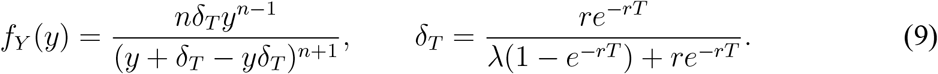
2. Conditional on *Y* = *y*, let the random variables *H*_1_,…, *H*_*n*-1_ be i.i.d. with probability density function on (0, *T*) given by

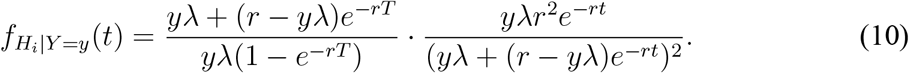

Note that the formula for the density of *Y* comes from equation (12) in Lambert^34^, and that *δ_T_* here is 1 – *a* in Lambert^34^. The density for *H_i_* comes from equation (7) in Lambert^34^. One can check that the resulting joint density for *H*_1_,…, *H*_*n*-1_ matches the joint density for the ordered coalescence times given in Proposition 19 of Harris et al.^33^.

While Lambert’s construction is only exact when the birth and death rates are constant over time, leading to a population which grows exponentially at a constant rate, Cheek^63^ has shown that under certain conditions, the construction remains approximately valid even when the growth rate of the population slows over time, provided that the population is still growing superlinearly at the time *T* when the sample is taken. For example, this method should give a good approximation in certain models of logistic population growth, provided that the sample is taken before the population reaches a fraction *x* of its carrying capacity, where 0 < *x* < 1^63^. Consequently, we believe that our methods, while derived in the case of constant birth and death rates, may be more broadly applicable.

### Internal and external branch lengths

Here, we state our main limit theorem, which describes the lengths of the internal and external branches for the genealogical tree of a birth-death process when *T* and *n* are large. The asymptotic distribution of the internal branch lengths can be used to estimate the net growth rate of a clone, while the ratio of external to internal branch lengths could provide an estimate of the clone age when the growth rate is known, as detailed further in Estimating the clone age.

We call a branch of the genealogical tree internal if it is ancestral to between 2 and *n* – 1 of the *n* leaves (red edges in Figure 1C-D) and external if it is ancestral to only one of the *n* leaves (blue edges in Figure 1C-D). Note that a mutation along an internal branch will be inherited by more than one of the sampled individuals, while a mutation along an external branch will be unique to one of the sampled individuals. Therefore, one can estimate the internal and external branch lengths from the number of shared and private mutations respectively.

The site frequency spectrum and the allele frequency spectrum have previously been studied for populations whose genealogy is described by a coalescent point process^47–51^. Because the internal and external branch lengths are closely related to the site frequency spectrum, our methods for using the coalescent point process to understand the internal and external branch lengths are similar to the methods used in these previous works. However, these earlier results are applicable when we are interested in the site frequency spectrum of the entire population, or when each individual is sampled independently with some probability *p*. Our result pertains to the case of a sample of fixed size *n* from a much larger population, leading to a star-shaped genealogical tree with long external branches in which most of the coalescence occurs near the root of the tree.

The asymptotic distribution for the internal and external branch lengths was obtained for the classical Kingman’s coalescent^64^ and for for coalescents with multiple mergers^65,66^. Recently, an asymptotic result for external branch lengths in Yule trees was proved^67^. However, as far as we know, such results have not previously been established for a sample of size *n* from a birth-death tree.

To state our theorem, we need to consider a sequence of birth-death processes indexed by the sample size *n*, and the time at which the sample is taken, which we will now denote by *T_n_*, must tend to infinity with *n*. We will also write *λ_n_*, *μ_n_*, *r_n_*, and *ν_n_* for the birth, death, growth, and mutation rates respectively to emphasize that we allow them to depend on *n*. We will let 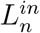 and 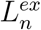 denote the total length of all internal and external branches respectively in the genealogical tree.

#### Theorem 1.

*Suppose*

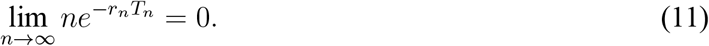

*Then, using 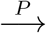 to denote convergence in probability as n →∞, we have*

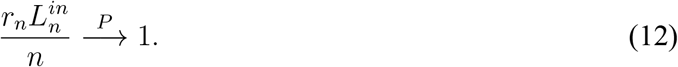

*Furthermore, suppose instead we have*

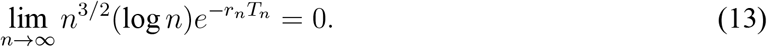

*Let Z have a standard normal distribution, and let W have an exponential distribution with mean 1, independent of Z. Then*

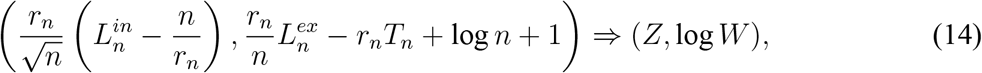

*where ⟹ denotes convergence in distribution as n →∞*.

Recall that the expected population size at time *T_n_* is *e^r_n_T_n_^*, so the condition (11) means that the sample size *n* must be much smaller than the population size. Under this condition, the total internal branch length 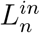 is close to *n*/*r_n_* with high probability, which means the growth rate estimate (4) should be accurate. Under the stronger condition (13), the distribution of the total internal branch length 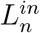 is approximately normal with mean *n*/*r_n_* and standard deviation 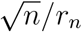, which we denote by

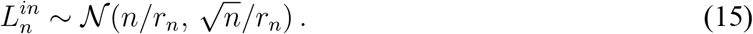

This means that the confidence interval in (5) should be accurate.

Because 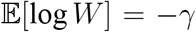, where *γ* ≈ .577 is Euler’s constant, Theorem 1 also suggests that for the total external branch length,

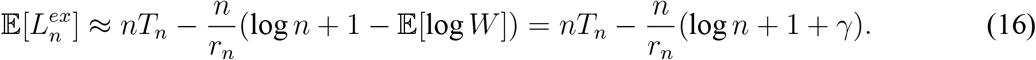

### Shared and private mutations

Let 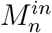 denote the number of mutations that appear on two or more of the sampled individuals, and let 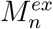 denote the number of mutations that appear on only one of the sampled individuals. Because we are assuming that mutations occur along each lineage at rate *ν_n_*, the conditional distribution of 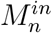 given 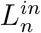 is Poisson with mean 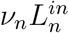, and likewise for 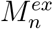. In particular, we have

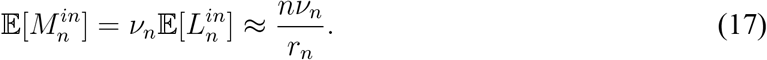

and, using the conditional variance formula,

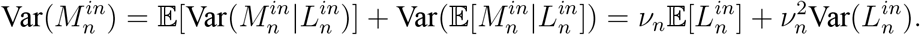

Note that the approximation for 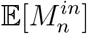 is consistent with (8) because 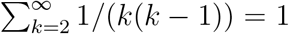. The following corollary to Theorem 1 shows that 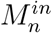 has an asymptotically normal distribution.

#### Corollary 2.

*Suppose that (13) holds and that*

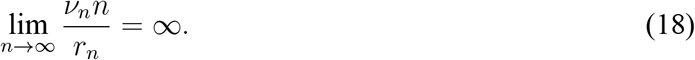

*Let*

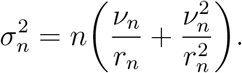

*Let Z have a standard normal distribution. Then*

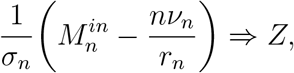

*where ⟹ denotes convergence in distribution as n →∞*.

Also, using (16) we have the approximation

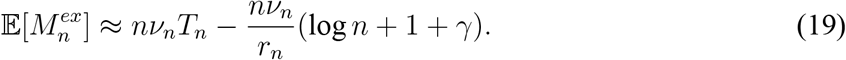

A similar formula was derived by Durrett^42^. For the private mutations 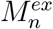, depending on the mutation rate, the dominant source of fluctuations could either be the Gaussian fluctuations from the mutations process or the non-Gaussian fluctuations from the random variable *W*.

### Comparing analytical estimates to those using Phylofit

As described by Williams et al., Phylofit is “an efficient MCMC approach that models selection/ growth by directly fitting the three parameter deterministic phase population trajectory using the joint probability density of coalescence times given the population size trajectory”^22^. The three parameters mentioned are the growth rate, the total number of hematopoietic stem cells, and the midpoint time of a deterministic logistic growth population trajectory. The likelihood function for the coalescence times is based on Equation 1 in Lan et. al^68^. In this sense, it is similar to our approach, leveraging the information provided by coalescence times, but is based on the Kingman coalescent with deterministic growth rather than a stochastic growth process. There are two ways to run Phylofit, only one of which is directly comparable to our methods.

Phylofit optionally incorporates Aberrant Cell Fraction (ACF) into its calculation of the likelihood. Aberrant Cell Fraction is simply the number of sampled cells within a clone divided by the total number of sampled cells. We refer to it alternatively as “sampled clonal fraction” in Application to human blood datasets, but use “Aberrant Cell Fraction” here for consistency with the terminology in Williams et al. ^22^. In this sense, it is analogous to the variant allele frequency that would be observed in bulk whole genome sequencing. When incorporating ACF, Phylofit assumes that the population trajectory of the clone is logistic, with a carrying capacity for the clone equal to the total number of hematopoietic stem cells. While coalescence events provide information from the early history of the clone, when the total clone size is on the order of the sample size *n*, the ACF provides information on the clone size at the time of sampling. Therefore, while the assumptions for our methods simply require an early expansion phase with constant birth and death rates, Phylofit with ACF assumes a logistic population size trajectory with a carrying capacity indicating that a clone becomes completely dominant in the blood (ACF=1). This is a much stronger assumption than is necessary for our methods or when using Phylofit without incorporating ACF. If the logistic population size trajectory is not the actual population size trajectory of a clone, the estimated growth rate from Phylofit with ACF will be affected. Moreover, the population size of a clone at the time of sampling is likely affected by treatments if applied, other competing expansions, and/or other reasons that carrying capacities would be lower than the total HSC pool. In fact, deceleration of clonal expansion rates diverging from the logistic trajectory is observed in the recent work published by Mitchell et al.^24^ and Fabre et al.^19^. Therefore, we compare our estimates to Phylofit without ACF, as this method, like our methods, estimates the growth rate during the expansion phase of the clone. However, published results in Williams et al. ^22^ consider ACF for growth rate inference, so our results using Phylofit differ from those which they present. For more details, see Supplementary section 3.2.

### Estimating the clone age

While we did not find a dataset appropriate for applying our method of estimating the clone age (i.e., time from clone initiation to time of sampling), we show here how it can be done when the growth rate of a clone is known and the mutation rate is unknown. If the mutation rate is known, then estimating the clone age is straightforward because we can simply divide the average number of mutations on the sampled individuals by the mutation rate. We therefore focus on how to estimate the clone age when the growth rate is known but the mutation rate is unknown. Note first that it is not possible to make such an estimate by using only the shared mutations. To see this, consider the figure below in which the dots represent mutations. Because the genealogical tree is nearly star-shaped, we will see the same shared mutations regardless of whether we take the sample at time *T*/2 or time *T*. The only difference is that if the sample is taken at time *T*, we will see more private mutations. We therefore estimate the tumor age by comparing the number of shared mutations to the number of private mutations. From (17) and (19), a natural estimate of the age of the tumor is

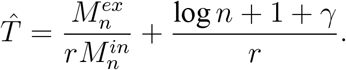

The result below establishes some asymptotic properties of this estimate, and shows how to obtain a confidence interval for *T*. Condition (20) is needed to ensure that the Gaussian fluctuations from the lengths of the internal branches and the mutations process are the dominant source of fluctuations, rather than the non-Gaussian fluctuations from the random variable *W* that measures the initial growth of the branching process.

#### Corollary 3.

*Suppose the following conditions hold:*

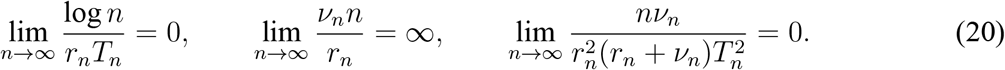

*Define*

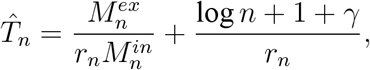

*as introduced above. Let Z have a standard normal distribution. Then*

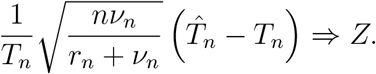

From this result, one can show that for 0 < *α* < 1, an asymptotically valid 100(1 – *α*)% confidence interval for *T_n_* can be obtained by

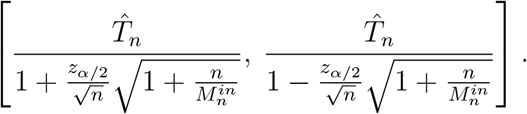

### Longitudinal modeling

#### Longitudinal clone inclusion criteria

As noted in Longitudinal validation of clone growth estimates, we only use data for longitudinal validation that has multiple timepoints sampled from a single cell type, as sampled cell type may affect the estimated clonal fraction. For example, data from bone marrow granulocytes may show a different clonal fraction than data from peripheral blood mononuclear cells, even if those samples are taken at the same time. This presents obvious difficulties for longitudinal growth modeling, so we exclude such data. This is discussed further in Supplementary section 3.2 and shown in Supp. Fig. 5.

In longitudinal data from individuals without hematological malignancies^19^, sequential data typically shows a lower but increasing VAF, making it ideal for our estimates of early growth rate. However, in longitudinal data from individuals with MPN^22^, increased clonal competition and treatment gives rise to more complicated dynamics. Many clones which at first appear advantageous are outcompeted by clones with higher fitness or knocked down by treatment, leading to clones occasionally decreasing in Variant Allele Frequency (VAF) over the sampled time period. To avoid these external effects, which are not representative of the growth rate during the early expansion phase, we did not consider longitudinal data from clones decreasing in size. Any data points featuring more than a 20% decrease in VAF from a previous timepoint were removed. Removal due to a 20% drop was only applied if the previous VAF was ≥ 0.05, to avoid removing data due to small fluctuations at low VAF. After removing any decreasing data, we required 4 data points in total and at least 2 with a VAF > 0 and ≤ 0.25. Finally, because we are looking for expanding clones, we remove any clones which do not increase in VAF by at least 0.05.

This filtering leaves us with 60 clones, with 56 from Fabre et al.^19^ and 4 from Williams et al.^22^. Of these, one clone from Williams et al.^22^ and two clones from Fabre et al.^19^ have sufficient matched single cell data to make growth rate estimates from both data modalities. We note that the only MPN clone with matched single-cell and longitudinal data (PD9478: *JAK2* + *DNMT3A*) is also from a patient that is essentially untreated, with the only intervention being venesection (bleeding). Further, the *JAK2* + *DNMT3A* clone in this patient appears to be the only large expansion, and there are 11 longitudinal samples from the same cell type, making this clone an ideal candidate for validation.

From the remaining 57 clones without matched single cell data, we identified 17 clones with mutations in the same driver gene as a single cell clone. We then excluded the longitudinal data from two MPN clones from Williams et al.^22^, as treatment likely affected the VAF in both cases. Further, we removed one *JAK2* clone where there was a competing clone at high VAF, as this was likely to affect the growth rate. After fitting the logistic growth rate model to each of the remaining clones, as detailed below in Logistic modeling of longitudinal data, we removed one DNMT3A clone where the fit failed to converge. After all filtering steps, there were 13 longitudinal clones matching to 10 single cell clones. *DNMT3A* mutant clones were the most abundant, with 7 *DNMT3A* clones having longitudinal data and 5 having single cell data. There were 3 longitudinal *JAK2* clones and 4 single cell *JAK2* clones. There were 3 longitudinal *PPM1D* clones and 1 single cell *PPM1D* clone.

#### Logistic modeling of longitudinal data

We used the *nls()* function from the *stats* package in R, with the port algorithm, to perform fitting to the following logistic growth equation, which models the VAF over time:

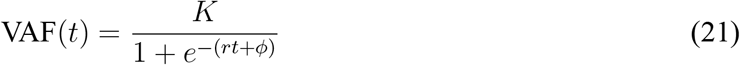

Bounds for *K*, representing the carrying capacity, are [0, 0.5], as all mutations are in diploid regions. Bounds for *r* are [0, 5] per year, and bounds for *φ* are [−500, 0]. 95% confidence intervals are found by assuming normality of the parameter estimate, using *r* ± 1.96 * stdError to calculate the bounds. Midpoint time of the logistic curve is given by *t_m_* = −*φ*/*r*.

Importantly, we do not assume a carrying capacity *K* equal to a variant allele frequency of 0.5, instead allowing the carrying capacity to be fit simultaneously with the growth rate and midpoint time. This is a distinction from the inherent assumptions in the Phylofit with ACF approach, which is discussed in Comparing analytical estimates to those using Phylofit and further in Supplementary section 3.2. Our decision to fit *K* rather than fixing it at 0.5 is motivated by data^19^ showing growth rates slowing more than would be expected by a logistic fit with *K* = 0.5, even in the absence of treatment. In fitting to longitudinal data from 16 clones (3 from Figure 6A-C and 13 from Figure 6E), nine clones have a fit VAF carrying capacity below 0.4, consistent with the claim that clones do not always saturate at an allele frequency of 0.5. A possible explanation for this lower carrying capacity is lineage bias which may lead to clonal dominance in only a subset of the blood progenitors. For example, *JAK2* mutants were found at higher clonal fractions in megakaryocyte and erythroid progenitors^23^. Variant allele frequency in whole blood may saturate below 0.5, even when a clone has become completely dominant within a specific type of progenitor (i.e. Megakaryocyte Erythroid Progenitors (MEP) in *JAK2* mutants). In fact, single cell clonal fraction data from the 13 MPN patients we have analyzed shows no example of a somatic clone that is completely dominant, despite matched stromal ^23^ or buccal^22^ normal cell samples distinguishing between somatic and germline variants.

It should be noted that a logistic growth model with any carrying capacity may not be the most appropriate model for clones in the blood, especially in the presence of other clones and/or treatment^54^. Our coalescent methods avoid this dependence on a particular model by assuming only that exponential growth occurs immediately following the initiation of a clone, while its population size is still on the order of the sample size, *n*. However, longitudinal validation requires the choice of a particular growth model in order to estimate a growth rate. Based on previous work exploring practical parameter identifiability in sigmoid growth models^46^, the logistic model outperformed Gompertz and Richards’ models, which is why we use it for longitudinal validation.

**Figure 7:**
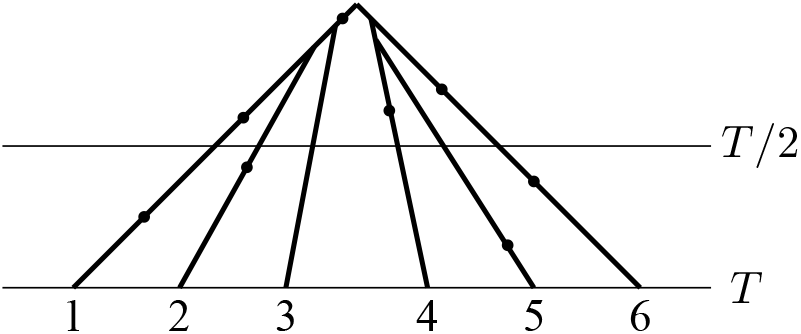
Six lineages sampled at times *T*/2 and *T*

## Data and code availability

We have created an open source *r* package called *cloneRate* to perform growth rate estimation using ultrametric or mutation-based phylogenetic trees as input data (https://github.com/bdj34/cloneRate/). *cloneRate* also includes methods for rapid generation of exact sampled trees from supercritical birth-death processes based on the work of Lambert^34^. Users can apply our methods to input data and also recreate our results presented herein using published data^19,22–24^. The de-identified ultrametric trees derived from mutational data from these papers are included in the package for convenience, please cite the appropriate references if this data is used. For method comparison, we also include the birth-death MCMC targeting the likelihood introduced in Equation (5) of Stadler^31^ in *cloneRate*. All modeling results for simulated and real data are provided as supplementary tables.

## Notes

### Competing Interest Statement

The authors have declared no competing interest.

### Summary of Updates

This updated manuscript contains additional results using a birth-death Markov chain Monte Carlo (MCMC) inference technique for comparison.

