## Supplementary material for "Estimating single cell clonal dynamics in human blood using coalescent theory"

### 1 Supplementary Information - Proofs

#### 1.1 Approximating the coalescent point process

Recall that we are considering a continuous-time birth-death process in which each individual gives birth at rate $\lambda$ and dies at rate $\mu$, and we let $r = \lambda - \mu$ be the growth rate of the process. We assume $r > 0$. According to Theorem 3 and Corollary 4 of[34], we can construct the genealogy of a sample of size $n$ from this process at time $T$, conditioned on the population having size at least $n$ at time $T$, by first defining the coalescence times $H_1, \dots, H_{n-1}$ and then using the coalescent point process to obtain the genealogical tree, as described in Figure 1. Recall that we can obtain the random variables $H_1, \dots, H_{n-1}$ in the following way, where we record the dependence on $n$ and $T$ in the notation:

1. Choose a random variable $Y_{n,T}$ with probability density function on $(0, 1)$ given by

$$f_{Y_{n,T}}(y) = \frac{n\delta_T y^{n-1}}{(y + \delta_T - y\delta_T)^{n+1}}, \qquad \delta_T = \frac{re^{-rT}}{\lambda(1 - e^{-rT}) + re^{-rT}}.$$

2. Conditional on $Y_{n,T} = y$, let the random variables $H_{i,n,T}$ for $1 \leq i \leq n-1$ be i.i.d. with probability density function on $(0, T)$ given by

$$f_{H_{i,n,T}|Y_{n,T}=y}(t) = \frac{y\lambda + (r - y\lambda)e^{-rT}}{y\lambda(1 - e^{-rT})} \cdot \frac{y\lambda r^2 e^{-rt}}{(y\lambda + (r - y\lambda)e^{-rt})^2}.$$

Our goal is to take a limit as $T \to \infty$ and then as $n \to \infty$ to obtain the approximation to the coalescent point process that was described in the section Approximating genealogy using a coalescent point process. We first make a change of variables. We can write $Y_{n,T}$ as $\delta_T \cdot Q_{n,T}$, where $Q_{n,T}$ has density

$$f_{Q_{n,T}}(q) = \frac{nq^{n-1}}{(1 + q - q\delta_T)^{n+1}}, \quad q \in (0, 1/\delta_T).$$

Let

$$G_{i,n,T} = T - H_{i,n,T}.$$

Note that $G_{i,n,T}$ represents the amount of time, after time zero, that the coalescence event occurred, whereas $H_{i,n,T}$ represents the amount of time, before the sampling time, that the coalescence event occurred. It will sometimes be more convenient to work with the times $G_{i,n,T}$. Replacing $y$ by $\delta_T \cdot q$, the density for $G_{i,n,T}$ given $Q_{n,T} = q$ is, for $0 < t < T$,

$$f_{G_{i,n,T}|Q_{n,T}=q}(t) = \frac{q\delta_T\lambda + (r - q\delta_T\lambda)\, e^{-rT}}{q\delta_T\lambda\,(1 - e^{-rT})} \cdot \frac{q\delta_T\lambda r^2 e^{-r(T-t)}}{\left(q\delta_T\lambda + (r - q\delta_T\lambda)\, e^{-r(T-t)}\right)^2}.$$

By writing $U_{i,n,T} = rG_{i,n,T} - \log q$, we have, for $-\log q < u < rT - \log q$,

$$f_{U_{i,n,T}|Q_{n,T}=q}(u) = \frac{q\delta_T\lambda + (r - q\delta_T\lambda)\,e^{-rT}}{q\delta_T\lambda\,(1 - e^{-rT})} \cdot \frac{q\delta_T\lambda r^2 e^{-rT} q e^u}{(q\delta_T\lambda + (r - q\delta_T\lambda)\,e^{-rT} q e^u)^2} \cdot \frac{1}{r}$$
$$= \frac{q\delta_T\lambda + (r - q\delta_T\lambda)\,e^{-rT}}{q\delta_T\lambda(1 - e^{-rT})} \cdot \frac{\delta_T\lambda r e^{-rT} e^u}{(\delta_T\lambda + (r - q\delta_T\lambda)\,e^{-rT} e^u)^2}.$$

Thus, to obtain the coalescence times $H_{i,n,T}$, we can:

1. Choose $Q_{n,T}$ from the density

$$f_{Q_{n,T}}(q) = \frac{nq^{n-1}}{(1 + q - q\delta_T)^{n+1}}, \quad q \in (0, 1/\delta_T)\,.$$

2. Given $Q_{n,T} = q$, sample $\{U_{i,n,T}\}_{i=1}^{n-1}$ i.i.d. from the density

$$f_{U_{i,n,T}|Q_{n,T}=q}(u) = \frac{q\delta_T\lambda + (r - q\delta_T\lambda)\,e^{-rT}}{q\delta_T\lambda\,(1 - e^{-rT})} \cdot \frac{\delta_T\lambda r e^{-rT} e^u}{(\delta_T\lambda + (r - q\delta_T\lambda)\,e^{-rT} e^u)^2},$$
$$u \in (-\log q, rT - \log q).$$

3. Let $H_{i,n,T} = T - \frac{1}{r}\,(\log Q_{n,T} + U_{i,n,T})$.

We now take a limit as $T \to \infty$. We note that the $T \to \infty$ limit was previously considered, in a more general setting allowing for non-binary branching, in[63,69]. For fixed $\lambda$, we have $\delta_T \to 0$ as $rT \to \infty$. Therefore, the density $f_{Q_{n,T}}(q)$ converges pointwise to the density

$$f_{Q_{n,\infty}}(q) = \frac{nq^{n-1}}{(1 + q)^{n+1}}, \quad q \in (0, \infty).$$

By the definition of $\delta_T$, for fixed $\lambda$, we have $\delta_T\lambda \sim re^{-rT}$ as $rT$ goes to $\infty$, where $\sim$ means that the ratio of the two sides tends to 1. Therefore, the conditional density $f_{U_{i,n,T}|Q_{n,T}=q}$ converges pointwise to

$$f_{U_{i,n,\infty}|Q_{n,\infty}=q}(u) = \frac{q+1}{q} \cdot \frac{e^u}{(1 + e^u)^2}, \quad u \in (-\log q, \infty).$$

Therefore, to obtain approximate coalescence times $H_{i,n,\infty}$ for $1 \le i \le n-1$, we can

1. Choose $Q_{n,\infty}$ from the density

$$f_{Q_{n,\infty}}(q) = \frac{nq^{n-1}}{(1 + q)^{n+1}} dq, \quad q \in (0, \infty),$$

2. Given $Q_{n,\infty} = q$, sample $\{U_{i,n,\infty}\}_{i=1}^{n}$ i.i.d. from the density

$$f_{U_{i,n,\infty}|Q_{n,\infty}=q}(u) = \frac{q+1}{q} \cdot \frac{e^u}{(1 + e^u)^2}, \quad u \in (-\log q, \infty),$$

3. Let $H_{i,n,\infty} = T - \frac{1}{r}\left(\log Q_{n,\infty} + U_{i,n,\infty}\right)$.

Next, we take a limit as $n \to \infty$. Let $\widetilde{Q}_{n,\infty} = Q_{n,\infty}/n$, which has density

$$f_{\widetilde{Q}_{n,\infty}}(q) = \frac{q^{n-1}}{\left(\frac{1}{n} + q\right)^{n+1}}, \quad q \in (0, \infty).$$

As $n$ goes to infinity, $f_{\widetilde{Q}_{n,\infty}}(q)$ converges pointwise to

$$\lim_{n\to\infty} \frac{q^{n-1}}{\left(\frac{1}{n} + q\right)^{n+1}} = \frac{e^{-1/q}}{q^2}, \quad q \in (0, \infty),$$

which is the density function for $1/W$ where $W$ has an exponential distribution with rate 1. Conditional on $\widetilde{Q}_{n,\infty} = q$, the density of $U_{i,n,\infty}$ for $1 \le i \le n-1$ is

$$f_{U_{i,n,\infty}|\widetilde{Q}_{n,\infty}=q}(u) = \frac{nq+1}{nq} \cdot \frac{e^u}{(1+e^u)^2}, \quad u \in (-\log q - \log n, \infty)\,.$$

For each $q > 0$, as $n$ goes to infinity, $f_{U_{i,n,\infty}|\widetilde{Q}_{n,\infty}=q}(u)$ converges pointwise to

$$\lim_{n\to\infty} f_{U_{i,\infty,\infty}|\widetilde{Q}_{n,\infty}=q}(u) = \frac{e^u}{(1+e^u)^2}, \quad u \in (-\infty, \infty),$$

which is the density function for the standard logistic distribution. Therefore, when both $T$ and $n$ are large, the coalescence times $H_{i,n,T}$ can be well approximated in the following way, which was described the section Approximating genealogy using a coalescent point process. Note that here, and frequently in the rest of the document, we write $U_i$ in place of $U_{i,\infty,\infty}$ to lighten notation.

1. Let $W$ have an exponential distribution with mean one.

2. Let $\{U_i\}_{i=1}^{n-1}$ be i.i.d. random variables with density

$$f_{U_i}(u) = \frac{e^u}{(1+e^u)^2}, \quad u \in (-\infty, \infty)$$

3. Let

$$H_i = T - \frac{1}{r}\Big(\log(1/W) + \log n + U_i\Big).$$

#### 1.2 Error bounds

In this section, we will show that the random variables $H_i$ give good approximations to the random variables $H_{i,n,T}$. Recall that $G_{i,n,T} = T - H_{i,n,T}$, and likewise define $G_{i,n,\infty} = T - H_{i,n,\infty}$ and $\widetilde{G}_{i,n,\infty} = T - H_i$. It then suffices to compare the random variables $\widetilde{G}_{i,n,\infty}$ and $G_{i,n,T}$. We will always assume that $n < e^{rT}$, which holds for sufficiently large $n$ when (13) is satisfied.

To show that the random variables $\widetilde{G}_{i,n,\infty}$ give a good approximation to the random variables $G_{i,n,T}$, we will use the technique of coupling. That is, we will construct these random variables on the same probability space in such a way that the absolute value of their difference is small with high probability. More precisely, suppose a random variable $X$ has distribution $\mu_X$, and a random variable $Y$ has distribution $\mu_Y$. When we say that the random variables $X$ and $Y$ can be coupled so that a certain condition holds, we mean that on some probability space, we can define random variables $X'$ and $Y'$ such that $X'$ has distribution $\mu_X$, $Y'$ has distribution $\mu_Y$, and the random variables $(X', Y')$ satisfy the given condition.

We will often use what is known as the maximal coupling (see section 4.4 of chapter 1 of[70]). If $X$ and $Y$ are random variables with probability density functions $f$ and $g$ respectively, then the random variables $X$ and $Y$ can be coupled so that $P(X = Y) = \int_{-\infty}^{\infty} f(x) \wedge g(x)\, dx$ and $P(X \neq Y) = \frac{1}{2}\int_{-\infty}^{\infty} |f(x) - g(x)|\, dx$.

Throughout this section, $C$ will be some constant varying from line to line, independent of the parameters $n, T, \lambda$ and $\mu$. By $f(x) = g(x)(1 + O(h(x)))$, we mean $|f(x) - g(x)| \leq Cg(x)h(x)$.

**Lemma 4.** *Suppose $n \leq h(n) \leq e^{rT-1}$. Then we can couple $Q_{n,T}$ and $Q_{n,\infty}$ so that*

$$\mathbb{P}(Q_{n,T} \neq Q_{n,\infty}) \leq Cne^{-rT}, \qquad \mathbb{P}(Q_{n,T} = Q_{n,\infty} > h(n)) \leq \frac{n}{h(n)},$$

*and we can also couple $U_{i,n,T}$ and $U_{i,n,\infty}$ so that*

$$\mathbb{E}\left[|U_{i,n,T} - U_{i,n,\infty}| \mathbb{1}_{\{Q_{n,T} = Q_{n,\infty} \leq h(n)\}}\right] \leq \frac{C \log\left(e^{rT}/h(n)\right)}{e^{rT}/h(n)}.$$

*Proof.* We can couple $Q_{n,T}$ and $Q_{n,\infty}$ so that

$$\mathbb{P}\left(Q_{n,T} = Q_{n,\infty}\right) = \int_0^\infty f_{Q_{n,T}}(q) \wedge f_{Q_{n,\infty}}(q)\, dq.$$

Likewise, on the event $\{Q_{n,T} = Q_{n,\infty} = q\}$, we can couple $U_{i,n,T}$ and $U_{i,n,\infty}$ so that

$$\mathbb{P}\left(U_{i,n,T} = U_{i,n,\infty} \mid Q_{n,T} = Q_{n,\infty} = q\right) = \int_0^\infty f_{U_{i,n,T}|Q_{n,T}=q}(u) \wedge f_{U_{i,n,\infty}|Q_{n,\infty}=q}(u)\, du.$$

On the event $\{Q_{n,T} \neq Q_{n,\infty}\}$, we take $U_{i,n,T}$ and $U_{i,n,\infty}$ to be arbitrary random variables with the prescribed conditional densities.

Since $\delta_T = \frac{r}{\lambda} e^{-rT}(1 + O(e^{-rT}))$, we have

$$\frac{nq^{n-1}}{(1 + q - q\delta_T)^{n+1}} = \frac{nq^{n-1}}{(1+q)^{n+1}} \left(1 + O\left(n\delta_T\right)\right), \tag{22}$$

$$\frac{q\delta_T\lambda + (r - q\delta_T\lambda)\, e^{-rT}}{q\delta_T\lambda\left(1 - e^{-rT}\right)} = \frac{q+1}{q}\left(1 + O\left(e^{-rT}\right)\right), \tag{23}$$

$$\frac{\delta_T\lambda r e^{-rT} e^u}{\left(\delta_T\lambda + (r - q\delta_T\lambda)\, e^{-rT} e^u\right)^2} = \frac{e^u}{(1+e^u)^2}\left(1 + O\left((q+1)e^{-rT}\right)\right). \tag{24}$$

Using (22), we have

$$\begin{aligned}\mathbb{P}(Q_{n,T} \neq Q_{n,\infty}) &= \int_0^\infty \frac{1}{2}\left|f_{Q_{n,\infty}}(q) - f_{Q_{n,T}}(q)\right|\,dq\\ &\leq Cn\delta_T \int_0^{1/\delta_T} \frac{nq^{n-1}}{(1+q)^{n+1}}\,dq + \int_{1/\delta_T}^\infty \frac{nq^{n-1}}{(1+q)^{n+1}}\,dq\\ &\leq Cn\delta_T + \int_{1/\delta_T}^\infty \frac{n}{q^2}\,dq\\ &\leq Cne^{-rT}.\end{aligned}$$

Also

$$\mathbb{P}(Q_{n,T} = Q_{n,\infty} > h(n)) \leq \mathbb{P}(Q_{n,\infty} > h(n)) = \int_{h(n)}^\infty \frac{nq^{n-1}}{(1+q)^{n+1}}\,dq \leq \int_{h(n)}^\infty \frac{n}{q^2}\,dq = \frac{n}{h(n)}.$$

Because $\lambda \geq r$, we have

$$\frac{1}{\delta_T} = \frac{\lambda(1-e^{-rT}) + re^{-rT}}{re^{-rT}} = \frac{\lambda}{r}(e^{rT}-1) + 1 \geq e^{rT} \geq h(n),$$

and therefore $q \leq 1/\delta_T$ whenever $q \leq h(n)$. Also, for $q \leq h(n)$, we have $e^{rT}/q = e$. It follows that on the event $Q_{n,T} = Q_{n,\infty} = q$, by (23) and (24) we have for $q \leq h(n)$,

$$\begin{aligned}&\mathbb{E}\left[|U_{i,n,T} - U_{i,n,\infty}|\,\middle|\,Q_{n,T} = Q_{n,\infty} = q\right]\\ &\quad\leq \mathbb{E}\left[(|U_{i,n,T}| + |U_{i,n,\infty}|)\,\mathbb{1}_{\{U_{i,n,T}\neq U_{i,n,\infty}\}}\,\middle|\,Q_{n,T} = Q_{n,\infty} = q\right]\\ &\quad= \int_{-\infty}^\infty |u|\left|f_{U_{i,n,T}|Q_{n,T}=q}(u) - f_{U_{i,n,\infty}|Q_{n,\infty}=q}(u)\right|\,du\\ &\quad\leq \int_{-\log q}^{rT-\log q} |u| C(q+1)e^{-rT}\cdot\frac{q+1}{q}\cdot\frac{e^u}{(1+e^u)^2}\,du + \int_{rT-\log q}^\infty |u|\cdot\frac{q+1}{q}\cdot\frac{e^u}{(1+e^u)^2}\,du\\ &\quad\leq C\left(\frac{(q+1)^2}{q}e^{-rT} + \frac{q+1}{q}\cdot\frac{\log\left(e^{rT}/q\right)}{e^{rT}/q}\right).\end{aligned} \tag{25}$$

Since the function $f(x) = (\log x)/x$ is decreasing on $(e,\infty)$, it follows that

$$\frac{\log\left(e^{rT}/q\right)}{e^{rT}/q} \leq \frac{\log\left(e^{rT}/h(n)\right)}{e^{rT}/h(n)}, \qquad \text{for } q \leq h(n).$$

Using this fact and integrating (25), we have

$$\begin{aligned}
&\mathbb{E}\left[|U_{i,n,T} - U_{i,n,\infty}|\,\mathbb{1}_{\{Q_{n,T}=Q_{n,\infty}\le h(n)\}}\right] \\
&\quad\le \int_0^{h(n)} C\left(\frac{(q+1)^2}{q}e^{-rT} + \frac{q+1}{q}\cdot\frac{\log\left(e^{rT}/q\right)}{e^{rT}/q}\right)\frac{nq^{n-1}}{(1+q)^{n+1}}\,dq \\
&\quad= C\int_0^{h(n)} (q+1)e^{-rT}\frac{nq^{n-2}}{(1+q)^n}\,dq + C\int_0^{h(n)} \frac{\log\left(e^{rT}/q\right)}{e^{rT}/q}\cdot\frac{nq^{n-2}}{(1+q)^n}\,dq \\
&\quad\le C(h(n)+1)e^{-rT}\int_0^\infty \frac{nq^{n-2}}{(1+q)^n}\,dq + \frac{C\log\left(e^{rT}/h(n)\right)}{e^{rT}/h(n)}\int_0^\infty \frac{nq^{n-2}}{(1+q)^n}\,dq \\
&\quad\le \frac{C\log\left(e^{rT}/h(n)\right)}{e^{rT}/h(n)},
\end{aligned}$$

which completes the proof. □

**Lemma 5.** *We can couple $\widetilde{Q}_{n,\infty}$ with $1/W$ so that*

$$\mathbb{P}\left(\widetilde{Q}_{n,\infty} \ne 1/W\right) \le \frac{C}{n},$$

*and we can also couple $U_{i,n,\infty}$ with $U_{i,\infty,\infty}$ so that*

$$\mathbb{E}\left[|U_{i,n,\infty} - U_{i,\infty,\infty}|\mathbb{1}_{\{\widetilde{Q}_{n,\infty}=1/W\}}\right] \le \frac{C\log n}{n}.$$

*Proof.* We can couple $\widetilde{Q}_{n,\infty}$ with $1/W$ so that

$$\mathbb{P}\left(\widetilde{Q}_{n,\infty} = 1/W\right) = \int_0^\infty f_{\widetilde{Q}_{n,\infty}}(q) \wedge f_{1/W}(q)\,dq.$$

On the event $\{\widetilde{Q}_{n,\infty} = q\}$, we can couple $U_{i,n,\infty}$ and $U_{i,\infty,\infty}$ so that

$$\mathbb{P}\left(U_{i,n,\infty} = U_{i,\infty,\infty} \mid \widetilde{Q}_{n,\infty} = q\right) = \int_0^\infty f_{U_{i,n,\infty}|\widetilde{Q}_{n,\infty}=q}(u) \wedge f_{U_{i,\infty,\infty}}(u)\,du.$$

We start with a bound on $\left|f_{\widetilde{Q}_{n,\infty}}(q) - f_{1/W}(q)\right|$. Note that

$$\frac{f_{\widetilde{Q}_{n,\infty}}(q)}{f_{1/W}(q)} = \frac{q^{n-1}}{\left(\frac{1}{n}+q\right)^{n+1}}\Bigg/\frac{e^{-1/q}}{q^2} = \exp\left(\frac{1}{q} - (n+1)\log\left(1+\frac{1}{nq}\right)\right).$$

Take $x_1 = 0$, $x_2 = 1/q - (n+1)\log(1+1/nq)$, and $F(x) = e^x$ so that $f_{1/W}(q) = F(x_1)f_{1/W}(q)$ and $f_{\widetilde{Q}_{n,\infty}}(q) = F(x_2)f_{1/W}(q)$. Using the standard inequality that $\frac{x}{x+1} < \log(1+x) < x$ for $x > 0$, we have

$$-\frac{1}{nq} < \frac{1}{q} - (n+1)\log\left(1+\frac{1}{nq}\right) < \frac{1-q}{(nq+1)q} < \frac{1}{nq^2}.$$

In particular,

$$|x_2 - x_1| \le \frac{1}{nq} \vee \frac{1}{nq^2},$$

and

$$f_{\widetilde{Q}_{n,\infty}}(q) = F(x_2) f_{1/W}(q) \le e f_{1/W}(q) \qquad \text{for } q \ge \frac{1}{\sqrt{n}}. \tag{26}$$

Applying the mean value theorem, we have

$$\begin{aligned} \left|f_{\widetilde{Q}_{n,\infty}}(q) - f_{1/W}(q)\right| &= f_{1/W}(q)|F(x_2) - F(x_1)| \\ &\le f_{1/W}(q)\,|F'(x_2) \vee F'(x_1)|\,|x_2 - x_1| \\ &= f_{1/W}(q)\,|F(x_2) \vee F(x_1)|\,|x_2 - x_1| \\ &\le \left|f_{\widetilde{Q}_{n,\infty}(q)} \vee f_{1/W}(q)\right| \left(\frac{1}{nq} \vee \frac{1}{nq^2}\right). \end{aligned} \tag{27}$$

For $g(q)$ where $g(q)$ can be $1/q$ or $1/q^2$, by (26) we have

$$\begin{aligned} &\int_0^\infty g(q)\left(f_{\widetilde{Q}_{n,\infty}}(q) \vee f_{1/W}(q)\right)\, dq \\ &\le \int_0^\infty g(q) f_{1/W}(q)\, dq + \int_0^\infty g(q) f_{\widetilde{Q}_{n,\infty}}(q)\, dq \\ &= \int_0^\infty g(q) f_{1/W}(q)\, dq + \int_0^{1/\sqrt{n}} g(q) f_{\widetilde{Q}_{n,\infty}}(q)\, dq + \int_{1/\sqrt{n}}^\infty g(q) f_{\widetilde{Q}_{n,\infty}}(q)\, dq \\ &\le (1+e)\int_0^\infty g(q) f_{1/W}(q)\, dq + \int_0^{1/\sqrt{n}} g(q) f_{\widetilde{Q}_{n,\infty}}(q)\, dq. \end{aligned} \tag{28}$$

Since $g(q)f_{1/W}(q)$ is integrable on $(0,\infty)$ the first term in (28) is bounded above by some absolute constant. For the second term, using the fact that $q^2 g(q)$ is integrable on $(0,1)$,

$$\begin{aligned} \int_0^{1/\sqrt{n}} g(q) f_{\widetilde{Q}_{n,\infty}}(q)\, dq &= \int_0^{1/\sqrt{n}} g(q) \frac{q^{n-1}}{\left(\frac{1}{n}+q\right)^{n+1}}\, dq \\ &= \int_0^{1/\sqrt{n}} q^2 g(q) \frac{q^{n-3}}{\left(\frac{1}{n}+q\right)^{n+1}}\, dq \\ &= \int_0^{1/\sqrt{n}} q^2 g(q) \frac{n^4 (nq)^{n-3}}{(1+nq)^{n+1}}\, dq \\ &\le \int_0^{1/\sqrt{n}} n^4 q^2 g(q) \left(\frac{nq}{1+nq}\right)^{n-3} dq \\ &\le \int_0^{1/\sqrt{n}} n^4 q^2 g(q) \left(1 - \frac{1}{1+\sqrt{n}}\right)^{n-3} dq \\ &\le n^4 \exp\left(-\frac{n-3}{1+\sqrt{n}}\right) \int_0^{1/\sqrt{n}} q^2 g(q)\, dq \\ &\le C. \end{aligned} \tag{29}$$

By (27), (28), and (29), we have

$$\begin{aligned}\mathbb{P}\big(\widetilde{Q}_{n,\infty} \neq 1/W\big) &= \frac{1}{2}\int_0^\infty \Big| f_{\widetilde{Q}_{n,\infty}}(q) - f_{1/W}(q)\Big|\, dq \\ &\leq \int_0^\infty \Big(f_{\widetilde{Q}_{n,\infty}}(q) \vee f_{1/W}(q)\Big)\left(\frac{1}{nq} \vee \frac{1}{nq^2}\right) dq \\ &\leq \frac{C}{n}. \end{aligned} \tag{30}$$

We also have

$$\begin{aligned}&\mathbb{E}\left[|U_{i,n,\infty} - U_{i,\infty,\infty}|\, \mathbb{1}_{\{\tilde{Q}_{n,\infty}=1/W\}}\right] \\ &\leq \int_0^\infty \int_0^\infty |u|\big|f_{U_{i,n,\infty}|\tilde{Q}_{n,\infty}=q}(u) - f_{U_{i,\infty,\infty}}(u)\big|\big(f_{1/W}(q) \wedge f_{\tilde{Q}_{n,\infty}}(q)\big)\, du\, dq \\ &\leq \int_0^\infty \left(\int_{-\infty}^{-\log q - \log n} \frac{|u|e^u}{(1+e^u)^2}\, du + \int_{-\infty}^\infty |u|\frac{1}{nq}\frac{e^u}{(1+e^u)^2}\, du\right)\frac{e^{-1/q}}{q^2}\, dq \\ &= \int_0^\infty \int_{-\infty}^{-\log q - \log n} |u|\frac{e^u}{(1+e^u)^2}\frac{e^{-1/q}}{q^2}\, du\, dq + \int_0^\infty \int_{-\infty}^\infty |u|\frac{1}{nq}\frac{e^u}{(1+e^u)^2}\frac{e^{-1/q}}{q^2}\, du\, dq \\ &\leq \int_{-\infty}^\infty \int_0^{1/(ne^u)} |u|\frac{e^u}{(1+e^u)^2}\frac{e^{-1/q}}{q^2}\, dq\, du + \frac{C}{n} \\ &= \int_{-\infty}^\infty |u|\frac{e^u}{(1+e^u)^2}e^{-ne^u}\, du + \frac{C}{n} \\ &\leq \frac{C\log n}{n}. \end{aligned} \tag{31}$$

The lemma follows from (30) and (31). ☐

For the rest of the proof, we will assume that we have a sequence of birth-death processes indexed by the sample size $n$, and we will denote the growth rate and the sampling time by $r_n$ and $T_n$. The next lemma, which is proved using the previous two lemmas, establishes that the random variables $G_{i,n,T_n}$ can be well approximated by the random variables $\widetilde{G}_{i,n,\infty}$.

**Lemma 6.** *Assume condition* (11) *holds. The random variables $G_{i,n,T_n}$ and $\widetilde{G}_{i,n,\infty}$ can be coupled so that as $n \to \infty$,*

$$\frac{r_n}{n}\sum_{i=1}^{n-1}\big|G_{i,n,T_n} - \widetilde{G}_{i,n,\infty}\big| \xrightarrow{P} 0.$$

*Assume condition* (13) *holds. The random variables $G_{i,n,T_n}$ and $\widetilde{G}_{i,n,\infty}$ can be coupled so that as $n \to \infty$,*

$$\frac{r_n}{\sqrt{n}}\sum_{i=1}^{n-1}\big|G_{i,n,T_n} - \widetilde{G}_{i,n,\infty}\big| \xrightarrow{P} 0.$$

*Proof.* Under condition (11), we take $h(n) = n^{1/2}e^{r_nT_n/2}$ in Lemma 4 so that

$$\lim_{n\to\infty}\frac{n}{h(n)} = \lim_{n\to\infty}\left(ne^{-r_nT_n}\right)^{1/2} = 0,$$

and

$$\lim_{n\to\infty}\frac{\log\left(e^{r_nT_n}/h(n)\right)}{e^{r_nT_n}/h(n)}=\lim_{n\to\infty}n^{1/2}e^{-r_nT_n/2}\log\left(n^{-1/2}e^{r_nT_n/2}\right).$$

Using condition (11) and the fact that $\lim_{x\to 0^+}x\log(1/x)=0$, the limit above is 0. Then by Lemma 4, as $n\to\infty$ we have

$$\mathbb{P}(Q_{n,T_n}=Q_{n,\infty}\leq h(n))\to 1, \tag{32}$$

and

$$\begin{aligned}&r_n\mathbb{E}\left[|G_{i,n,T_n}-G_{i,n,\infty}|\mathbb{1}_{\{Q_{n,T_n}=Q_{n,\infty}\leq h(n)\}}\right]\\&\quad=\mathbb{E}\left[|U_{i,n,T_n}-U_{i,n,\infty}|\mathbb{1}_{\{Q_{n,T_n}=Q_{n,\infty}\leq h(n)\}}\right]\leq\frac{C\log\left(e^{r_nT_n}/h(n)\right)}{e^{r_nT_n}/h(n)}\to 0.\end{aligned}$$

Summing over $1\leq i\leq n-1$ and applying Markov's inequality, we have

$$\frac{r_n}{n}\sum_{i=1}^{n-1}|G_{i,n,T_n}-G_{i,n,\infty}|\,\mathbb{1}_{\{Q_{n,T_n}=Q_{n,\infty}\leq h(n)\}}\xrightarrow{P}0. \tag{33}$$

Combining (32) and (33), we have

$$\frac{r_n}{n}\sum_{i=1}^{n-1}|G_{i,n,T_n}-G_{i,n,\infty}|\xrightarrow{P}0. \tag{34}$$

A similar argument using Lemma 5 instead implies that

$$\frac{r_n}{n}\sum_{i=1}^{n-1}|G_{i,n,\infty}-\widetilde{G}_{i,n,\infty}|\xrightarrow{P}0. \tag{35}$$

The first claim of the lemma follows from (34) and (35).

Under condition (13), we take $h(n)=n^{1/2}(\log n)^{-1/3}e^{r_nT_n/3}$ in Lemma 4 so that

$$\lim_{n\to\infty}\frac{n}{h(n)}=\lim_{n\to\infty}\left(n^{3/2}(\log n)e^{-r_nT_n}\right)^{1/3}=0,$$

and

$$\begin{aligned}&\lim_{n\to\infty}\sqrt{n}\cdot\frac{\log\left(e^{r_nT_n}/h(n)\right)}{e^{r_nT_n}/h(n)}\\&\quad=\lim_{n\to\infty}n(\log n)^{-1/3}e^{-2r_nT_n/3}\log\left(n^{-1/2}(\log n)^{1/3}e^{2r_nT_n/3}\right)\\&\quad=\lim_{n\to\infty}n(\log n)^{-1/3}e^{-2r_nT_n/3}\left\{\log\left(n^{-1}(\log n)^{1/3}e^{2r_nT_n/3}\right)+\log(n^{1/2})\right\}.\end{aligned} \tag{36}$$

Since

$$\lim_{n\to\infty}n(\log n)^{-1/3}e^{-2r_nT_n/3}=\lim_{n\to\infty}\left(n^{3/2}\left(\log n\right)e^{-r_nT_n}\right)^{2/3}\cdot(\log n)^{-1}=0,$$

and $\lim_{x\to 0^+} x\log(1/x) = 0$, we have

$$\lim_{n\to\infty} n(\log n)^{-1/3} e^{-2r_nT_n/3} \log\left(n^{-1}(\log n)^{1/3} e^{2r_nT_n/3}\right) = 0. \tag{37}$$

Also,

$$\lim_{n\to\infty} n(\log n)^{-1/3} e^{-2r_nT_n/3} \log(n^{1/2}) = \lim_{n\to\infty} \frac{1}{2}\left(n^{3/2}(\log n)e^{-r_nT_n}\right)^{2/3} = 0.$$

Combining this with (36) and (37), we deduce that

$$\lim_{n\to\infty} \sqrt{n}\cdot \frac{\log\left(e^{r_nT_n}/h(n)\right)}{e^{r_nT_n}/h(n)} = 0.$$

Then by Lemma 4, we have

$$\mathbb{P}(Q_{n,T_n} = Q_{n,\infty} \le h(n)) \to 1, \tag{38}$$

and

$$\begin{aligned} &r_n\sqrt{n}\,\mathbb{E}\left[|G_{i,n,T_n} - G_{i,n,\infty}|\mathbb{1}_{\{Q_{n,T_n}=Q_{n,\infty}\le h(n)\}}\right] \\ &\quad = \sqrt{n}\,\mathbb{E}\left[|U_{i,n,T_n} - U_{i,n,\infty}|\mathbb{1}_{\{Q_{n,T_n}=Q_{n,\infty}\le h(n)\}}\right] \le C\sqrt{n}\cdot\frac{\log\left(e^{r_nT_n}/h(n)\right)}{e^{r_nT_n}/h(n)} \to 0. \end{aligned}$$

Summing over $1\le i\le n-1$ and applying Markov's inequality, we have

$$\frac{r_n}{\sqrt{n}}\sum_{i=1}^{n-1} |G_{i,n,T_n} - G_{i,n,\infty}|\,\mathbb{1}_{\{Q_{n,T_n}=Q_{n,\infty}\le h(n)\}} \xrightarrow{P} 0. \tag{39}$$

Combining (38) and (39), we have

$$\frac{r_n}{\sqrt{n}}\sum_{i=1}^{n-1} |G_{i,n,T_n} - G_{i,n,\infty}| \xrightarrow{P} 0. \tag{40}$$

A similar argument using Lemma 5 instead implies that

$$\frac{r_n}{\sqrt{n}}\sum_{i=1}^{n-1} |G_{i,n,\infty} - \widetilde{G}_{i,n,\infty}| \xrightarrow{P} 0. \tag{41}$$

The second claim of the lemma follows from (40) and (41). □

#### 1.3 Approximating internal and external branch lengths

In this section, we show how to use the approximation to the coalescent point process to estimate the moments of the internal and external branch lengths. We first explain how to read off the internal and external branch lengths from the coalescent point process. Consider the coalescent point process for which the coalescence times are given by $H_{i,n,T_n}$. The length of the internal portion of the 0th branch, denoted by $L^{in}_{0,n,T_n}$, is

$$L^{in}_{0,n,T_n} = \max_{1 \leq i \leq n-1} H_{i,n,T_n} - H_{1,n,T_n} = G_{1,n,T_n} - \min_{1 \leq i \leq n-1} G_{i,n,T_n}.$$

Note that the top portion of the 0th branch, which is ancestral to all $n$ of the sampled individuals, is not counted towards the internal branch length. For $1 \leq i \leq n-2$, the length of the internal portion of the $i$th branch, denoted by $L^{in}_{i,n,T}$, is

$$L^{in}_{i,n,T_n} = (H_{i,n,T_n} - H_{i+1,n,T_n})^+ = (G_{i+1,n,T_n} - G_{i,n,T_n})^+.$$

The $(n-1)$st branch is entirely external. The total internal branch length is therefore

$$L^{in}_n = \sum_{i=0}^{n-2} L^{in}_{i,n,T_n}. \tag{42}$$

The length of the external portion of the 0th branch is

$$L^{ex}_{0,n,T_n} = H_{1,n,T_n} = T_n - G_{1,n,T_n}.$$

For $1 \leq i \leq n-2$, the length of the external portion of the $i$th branch is

$$L^{ex}_{i,n,T_n} = H_{i,n,T_n} \wedge H_{i+1,n,T_n} = T_n - G_{i,n,T_n} \vee G_{i+1,n,T_n}.$$

The length of the external portion of the $(n-1)$st branch is

$$L^{ex}_{n-1,n,T_n} = H_{n-1,n,T_n} = T_n - G_{n-1,n,T_n}.$$

The total external branch length is

$$L^{ex}_n = \sum_{i=0}^{n-1} L^{ex}_{i,n,T_n}. \tag{43}$$

We collect here some results about the logistic distribution that we will need later. We refer the reader to[71]. See, in particular, the formula following (1.10) in[71] for the variance, and (2.3.9) of[71] for the formula for the expected value of the maximum of $n$ i.i.d. logistic random variables.

**Lemma 7.** *Let $U_1, U_2, \ldots$ be i.i.d standard logistic random variables. Then $\mathbb{E}[U_1] = 0$ and $\mathrm{Var}(U_1) = \pi^2/3$. Also, we have*

$$\mathbb{E}\left[\max_{1 \leq i \leq n} U_i\right] = \sum_{i=1}^{n-1} \frac{1}{i}.$$

The next lemma shows that the internal and external branch lengths can be well approximated using the random variables $\widetilde{G}_{i,n,\infty}$ in place of $G_{i,n,T_n}$. For $1 \leq i \leq n-2$, we will use the notation

$$\widetilde{L}^{in}_{i,n,\infty} = (\widetilde{G}_{i+1,n,\infty} - \widetilde{G}_{i,n,\infty})^+$$

for the approximation to the length of the internal portion of the $i$th branch.

**Lemma 8.** *Assume condition* (11) *holds. Then as* $n \to \infty$*, the following hold:*

1. *For the internal branch length, we have*

$$\frac{r_n}{n}\left(\sum_{i=0}^{n-2} L^{in}_{i,n,T_n} - \sum_{i=1}^{n-2} \widetilde{L}^{in}_{i,n,\infty}\right) \xrightarrow{P} 0.$$

2. *For the external branch length, we have*

$$\frac{r_n}{n}\left(\sum_{i=0}^{n-1}(T_n - L^{ex}_{i,n,T_n}) - \sum_{i=1}^{n-2}(\widetilde{G}_{i,n,\infty} \vee \widetilde{G}_{i+1,n,\infty})\right) \xrightarrow{P} 0. \tag{44}$$

*Assume condition* (13) *holds. Then as* $n \to \infty$*, the following hold:*

1. *For the internal branch length, we have*

$$\frac{r_n}{\sqrt{n}}\left(\sum_{i=0}^{n-2} L^{in}_{i,n,T_n} - \sum_{i=1}^{n-2} \widetilde{L}^{in}_{i,n,\infty}\right) \xrightarrow{P} 0.$$

2. *For the external branch length, we have*

$$\frac{r_n}{\sqrt{n}}\left(\sum_{i=0}^{n-1}(T_n - L^{ex}_{i,n,T_n}) - \sum_{i=1}^{n-2}(\widetilde{G}_{i,n,\infty} \vee \widetilde{G}_{i+1,n,\infty})\right) \xrightarrow{P} 0. \tag{45}$$

*Proof.* The proofs under conditions (11) and (13) are essentially the same. We present the proof under condition (13) here. For the internal branch length, we have

$$\sum_{i=0}^{n-2} L^{in}_{i,n,T_n} - \sum_{i=1}^{n-2} \widetilde{L}^{in}_{i,n,\infty} = L^{in}_{0,n,T_n} + \sum_{i=1}^{n-2}\left(L^{in}_{i,n,T_n} - \widetilde{L}^{in}_{i,n,\infty}\right). \tag{46}$$

For the second term in (46), using the triangle inequality, we have

$$|L^{in}_{i,n,T_n} - \widetilde{L}^{in}_{i,n,\infty}| \le |G_{i,n,T_n} - \widetilde{G}_{i,n,\infty}| + |G_{i+1,n,T_n} - \widetilde{G}_{i+1,n,\infty}|.$$

Summing this for $1 \le i \le n-2$ and using Lemma 6, we have

$$\frac{r_n}{\sqrt{n}}\sum_{i=1}^{n-2}\left(L^{in}_{i,n,T_n} - \widetilde{L}^{in}_{i,n,\infty}\right) \xrightarrow{P} 0. \tag{47}$$

For the first term in (46), we have

$$L^{in}_{0,n,T_n} = G_{1,n,T_n} - \min_{1\le i\le n-1} G_{i,n,T_n}$$
$$\le \widetilde{G}_{1,n,\infty} - \min_{1\le i\le n-1} \widetilde{G}_{i,n,\infty} + |G_{1,n,T_n} - \widetilde{G}_{1,n,\infty}| + \sum_{i=1}^{n-1} |G_{i,n,T_n} - \widetilde{G}_{i,n,\infty}|. \tag{48}$$

By Lemma 6, we have

$$\frac{r_n}{\sqrt{n}} \left( |G_{1,n,T_n} - \widetilde{G}_{1,n,\infty}| + \sum_{i=1}^{n-1} |G_{i,n,T_n} - \widetilde{G}_{i,n,\infty}| \right) \xrightarrow{P} 0. \tag{49}$$

Finally, since $U_i$ has a symmetric distribution, by Lemma 7, we have

$$\mathbb{E}\left[\widetilde{G}_{1,n,\infty} - \min_{1\le i\le n-1} \widetilde{G}_{i,n,\infty}\right] = \frac{1}{r_n}\mathbb{E}\left[U_1 - \min_{1\le i\le n-1} U_i\right] = \frac{1}{r_n}\mathbb{E}\left[\max_{1\le i\le n-1} U_i\right] = \frac{1}{r_n}\sum_{i=1}^{n-2}\frac{1}{i} \le \frac{C\log n}{r_n}.$$

In particular, using Markov's Inequality, we have

$$\frac{r_n}{\sqrt{n}} \left( \widetilde{G}_{1,n,\infty} - \min_{1\le i\le n-1} \widetilde{G}_{i,n,\infty} \right) \xrightarrow{P} 0. \tag{50}$$

The claim for the internal branch length follows from (46), (47), (48), (49), and (50).

For the external branch length, we have

$$\sum_{i=0}^{n-1}(T_n - L^{ex}_{i,n,T_n}) - \sum_{i=1}^{n-2}\left(\widetilde{G}_{i,n,\infty} \vee \widetilde{G}_{i+1,n,\infty}\right)$$
$$= G_{1,n,T_n} + \sum_{i=1}^{n-2}\left((G_{i,n,T_n} \vee G_{i+1,n,T_n}) - (\widetilde{G}_{i,n,\infty} \vee \widetilde{G}_{i+1,n,\infty})\right) + G_{n-1,n,T_n}. \tag{51}$$

Similar to the argument for the internal branch length, we have

$$\frac{r_n}{\sqrt{n}} \sum_{i=1}^{n-2}\left((G_{i,n,T_n} \vee G_{i+1,n,T_n}) - (\widetilde{G}_{i,n,\infty} \vee \widetilde{G}_{i+1,n,\infty})\right) \xrightarrow{P} 0. \tag{52}$$

Also, we have

$$\frac{r_n}{\sqrt{n}} \cdot G_{1,n,T_n} \le \frac{r_n}{\sqrt{n}}\left(\widetilde{G}_{1,n,\infty} + |G_{1,n,T_n} - \widetilde{G}_{1,n,\infty}|\right)$$
$$\le \frac{1}{\sqrt{n}}\left(|\log(1/W)| + |U_1| + \log n + |G_{1,n,T_n} - \widetilde{G}_{1,n,\infty}|\right) \xrightarrow{P} 0, \tag{53}$$

and similarly for $G_{n-1,n,T_n}$. The claim for the external branch length follows from (51), (52), and (53). □

**Lemma 9.** *We have the following moment estimates for the internal branch length:*

$$\mathbb{E}\left[\sum_{i=1}^{n-2} \widetilde{L}^{in}_{i,n,\infty}\right] = \frac{1}{r_n}(n-2),$$
$$\mathrm{Var}\left(\sum_{i=1}^{n-2} \widetilde{L}^{in}_{i,n,\infty}\right) = \frac{1}{r_n^2}\left(n+\frac{\pi^2}{3}-4\right).$$

*Proof.* Since $U_i$ has a symmetric distribution, we have

$$\mathbb{E}[U_i \wedge U_{i+1}] = \mathbb{E}[(-U_i) \wedge (-U_{i+1})] = -\mathbb{E}[U_i \vee U_{i+1}].$$

By Lemma 7, we have

$$\begin{aligned}\mathbb{E}\left[\left(\widetilde{G}_{i+1,n,\infty} - \widetilde{G}_{i,n,\infty}\right)^+\right] &= \frac{1}{r_n}\mathbb{E}\left[(U_{i+1}-U_i)^+\right] \\ &= \frac{1}{2r_n}\mathbb{E}[U_{i+1} \vee U_i - U_{i+1} \wedge U_i] = \frac{1}{r_n}\mathbb{E}[U_{i+1} \vee U_i] = \frac{1}{r_n}. \qquad (54)\end{aligned}$$

Therefore,

$$\mathbb{E}\left[\sum_{i=1}^{n-2} \widetilde{L}^{in}_{i,n,\infty}\right] = \frac{1}{r_n}(n-2).$$

For the variance computation, we have

$$\begin{aligned}\mathbb{E}\left[\left(\left(\widetilde{G}_{i+1,n,\infty} - \widetilde{G}_{i,n,\infty}\right)^+\right)^2\right] &= \frac{1}{r_n^2}\mathbb{E}\left[\left((U_{i+1}-U_i)^+\right)^2\right] \\ = \frac{1}{2r_n^2}\mathbb{E}\left[(U_{i+1}-U_i)^2\right] &= \frac{1}{2r_n^2}\mathrm{Var}(U_{i+1}-U_i) = \frac{1}{r_n^2}\mathrm{Var}(U_i) = \frac{\pi^2}{3r_n^2}\end{aligned}$$

and, using Mathematica to evaluate the triple integral,

$$\begin{aligned}&\mathbb{E}\left[\left(\widetilde{G}_{i+1,n,\infty} - \widetilde{G}_{i,n,\infty}\right)^+\left(\widetilde{G}_{i+2,n,\infty} - \widetilde{G}_{i+1,n,\infty}\right)^+\right] \\ &= \frac{1}{r_n^2}\mathbb{E}\left[(U_{i+1}-U_i)^+\,(U_{i+2}-U_{i+1})^+\right] \\ &= \frac{1}{r_n^2}\int_{-\infty}^{\infty}\int_{u_i}^{\infty}\int_{u_{i+1}}^{\infty}(u_{i+1}-u_i)(u_{i+2}-u_{i+1})\frac{e^{u_i}}{(1+e^{u_i})^2}\frac{e^{u_{i+1}}}{(1+e^{u_{i+1}})^2}\frac{e^{u_{i+2}}}{(1+e^{u_{i+2}})^2}\,du_{i+2}\,du_{i+1}\,du_i \\ &= \left(2-\frac{\pi^2}{6}\right)\frac{1}{r_n^2}.\end{aligned}$$

For $i, j$ with $|i-j| \geq 2$, the random variables $(\widetilde{G}_{i+1,n,\infty} - \widetilde{G}_{i,n,\infty})^+$ and $(\widetilde{G}_{j+1,n,\infty} - \widetilde{G}_{j,n,\infty})^+$ are independent. Therefore, the variance of the internal branch length is given by:

$$\begin{aligned}\mathrm{Var}\left(\sum_{i=1}^{n-2}\widetilde{L}^{in}_{i,n,\infty}\right) &= \sum_{i=1}^{n-2}\sum_{j=1}^{n-2}\mathrm{Cov}\left(\widetilde{L}^{in}_{i,n,\infty},\widetilde{L}^{in}_{j,n,\infty}\right)\\ &= (n-2)\mathrm{Var}\left(\widetilde{L}^{in}_{1,n,\infty}\right)+2(n-3)\mathrm{Cov}\left(\widetilde{L}^{in}_{1,n,\infty},\widetilde{L}^{in}_{2,n,\infty}\right)\\ &= (n-2)\left(\frac{\pi^2}{3r_n^2}-\frac{1}{r_n^2}\right)+2(n-3)\left(\left(2-\frac{\pi^2}{6}\right)\frac{1}{r_n^2}-\frac{1}{r_n^2}\right)\\ &= \frac{1}{r_n^2}\left(n+\frac{\pi^2}{3}-4\right),\end{aligned}$$

which completes the proof. □

#### 1.4 Proof of Theorem 1

*Proof of Theorem 1.* We first suppose that (11) holds. It follows from Lemma 8 that to prove (12), it suffices to show that

$$\frac{r_n}{n}\sum_{i=1}^{n-2}\widetilde{L}^{in}_{i,n,\infty}\xrightarrow{P}1. \tag{55}$$

By Lemma 9, we have

$$\mathbb{E}\left[\left(\frac{r_n}{n}\sum_{i=1}^{n-2}\widetilde{L}^{in}_{i,n,\infty}-1\right)^2\right]=\mathrm{Var}\left(\frac{r_n}{n}\sum_{i=1}^{n-2}\widetilde{L}^{in}_{i,n,\infty}\right)+\left(\frac{n-2}{n}-1\right)^2=\frac{1}{n^2}\left(n+\frac{\pi^2}{3}-4\right)+\frac{4}{n^2}.$$

The result (55) now follows from Chebyshev's inequality.

Next, we suppose that (13) holds, and we need to show (14). Taking the negative of the second coordinate in (14) and using (42) and (43), we see that (14) is equivalent to the convergence

$$\left(\frac{r_n}{\sqrt{n}}\left(\sum_{i=0}^{n-2}L^{in}_{i,n,T_n}-\frac{n}{r_n}\right),\frac{r_n}{n}\sum_{i=0}^{n-1}\left(T_n-L^{ex}_{i,n,T_n}\right)-\log n-1\right)\Rightarrow(Z,\log(1/W)).$$

By Lemma 8, it suffices to show that

$$\left(\frac{r_n}{\sqrt{n}}\left(\sum_{i=1}^{n-2}\widetilde{L}^{in}_{i,n,\infty}-\frac{n}{r_n}\right),\frac{r_n}{n}\sum_{i=1}^{n-2}\left(\widetilde{G}_{i,n,\infty}\vee\widetilde{G}_{i+1,n,\infty}\right)-\log n-1\right)\Rightarrow(Z,\log(1/W)). \tag{56}$$

Using (54), for $1\le i\le n-2$, we have

$$r_n\left(\widetilde{L}^{in}_{i,n,\infty}-\mathbb{E}\left[\widetilde{L}^{in}_{i,n,\infty}\right]\right)=r_n\left(\left(\widetilde{G}_{i+1,n,\infty}-\widetilde{G}_{i,n,\infty}\right)^+-\frac{1}{r_n}\right)=(U_{i+1}-U_i)^+-1.$$

Recall that a sequence of random variables $(X_i)_{i=1}^{\infty}$ is said to be $m$-dependent if $(X_i)_{i\le k}$ and $(X_i)_{i>k+m}$ are independent for all $k$. Note that the sequence whose $i$th term is $(U_{i+1}-U_i)^+-1$ is 1-dependent. We can therefore apply the Central Limit Theorem for $m$-dependent sequences to this sequence. By

Theorem 3 of[72], it suffices to check the following conditions:

$$\liminf_{n\to\infty}\frac{1}{n}\mathrm{Var}\left(r_n\sum_{i=1}^{n-2}\widetilde{L}^{in}_{i,n,\infty}\right)>0,$$

$$\lim_{n\to\infty}\frac{1}{n}\sum_{i=1}^{n}\mathbb{E}\left[\left|r_n\widetilde{L}^{in}_{i,n,\infty}-\mathbb{E}\left[r_n\widetilde{L}^{in}_{i,n,\infty}\right]\right|^2\mathbb{1}_{\left\{\left|r_n\widetilde{L}^{in}_{i,n,\infty}-\mathbb{E}\left[r_n\widetilde{L}^{in}_{i,n,\infty}\right]\right|>\epsilon\sqrt{n}\right\}}\right]=0,\quad\forall\epsilon>0.$$

For the first condition, it follows from Lemma 9 that

$$\liminf_{n\to\infty}\frac{1}{n}\mathrm{Var}\left(r_n\sum_{i=1}^{n-2}\widetilde{L}^{in}_{i,n,\infty}\right)=1.$$

For the second condition, we have

$$\begin{aligned}&\frac{1}{n}\sum_{i=1}^{n}\mathbb{E}\left[\left|r_n\widetilde{L}^{in}_{i,n,\infty}-\mathbb{E}\left[r_n\widetilde{L}^{in}_{i,n,\infty}\right]\right|^2\mathbb{1}_{\left\{\left|r_n\widetilde{L}^{in}_{i,n,\infty}-\mathbb{E}\left[r_n\widetilde{L}^{in}_{i,n,\infty}\right]\right|>\epsilon\sqrt{n}\right\}}\right]\\&\quad=\mathbb{E}\left[\left|r_n\widetilde{L}^{in}_{1,n,\infty}-\mathbb{E}\left[r_n\widetilde{L}^{in}_{1,n,\infty}\right]\right|^2\mathbb{1}_{\left\{\left|r_n\widetilde{L}^{in}_{1,n,\infty}-\mathbb{E}\left[r_n\widetilde{L}^{in}_{1,n,\infty}\right]\right|>\epsilon\sqrt{n}\right\}}\right]\\&\quad=\mathbb{E}\left[|(U_{1,\infty,\infty}-U_{2,\infty,\infty})^+-1|^2\mathbb{1}_{\left\{|(U_{1,\infty,\infty}-U_{2,\infty,\infty})^+-1|>\epsilon\sqrt{n}\right\}}\right],\end{aligned}$$

which goes to $0$ as $n$ goes to infinity by the Dominated Convergence Theorem. Therefore, by the $m$-dependent Central Limit Theorem and Lemma 9,

$$\frac{r_n}{\sqrt{n}}\left(\sum_{i=1}^{n-2}\widetilde{L}^{in}_{i,n,\infty}-\frac{n}{r_n}\right)\Rightarrow Z,\tag{57}$$

where $Z$ has a standard normal distribution.

We now consider the sum

$$\sum_{i=1}^{n-2}\left(\widetilde{G}_{i,n,\infty}\vee\widetilde{G}_{i+1,n,\infty}\right)=\frac{n-2}{r_n}\log(1/W)+\frac{n-2}{r_n}\log n+\frac{1}{r_n}\sum_{i=1}^{n-2}(U_i\vee U_{i+1}).$$

By Lemma 7, we have $\mathbb{E}(U_1\vee U_2)=1$. Applying the Strong Law of Large Numbers separately to $(U_1\vee U_2)+(U_3\vee U_4)+\dots$ and $(U_2\vee U_3)+(U_4\vee U_5)+\dots$ and then recombining, we have

$$\frac{r_n}{n}\sum_{i=2}^{n-2}\left(\widetilde{G}_{i,n,\infty}\vee\widetilde{G}_{i+1,n,\infty}\right)-\log n-1\to\log(1/W)\quad\text{a.s.}\tag{58}$$

Combining this result with (57) gives (56), which completes the proof. □

**Remark 10.** Note that the convergence of the second coordinate holds even if we only have (44) rather than (45) in Lemma 8. Therefore, even under the weaker condition (11), we have the convergence

$$\frac{r_n}{n} L_n^{ex} - r_n T_n + \log n + 1 \Rightarrow \log W.$$

##### 1.5 Proof of Corollaries 2 and 3

To prove Corollaries 2 and 3, we need to consider the mutations. Let $M_n^{in}$ be the number of mutations inherited by between 2 and $n-1$ individuals, and let $M_n^{ex}$ be the number of mutations inherited by and exactly one individual. Recall that conditional on $(L_n^{in}, L_n^{ex})$, $M_n^{in}$ and $M_n^{ex}$ are independent Poisson random variables with means $\nu_n L_n^{in}$ and $\nu_n L_n^{ex}$ respectively. Therefore, there are two sources of fluctuations for $M_n^{in}$ and $M_n^{ex}$. There are Poissonian fluctuations coming from the mutations process, and fluctuations resulting from the randomness in the branch lengths. We decompose and rescale the fluctuations as follows. Let

$$A_n = \frac{M_n^{in} - \nu_n L_n^{in}}{\sqrt{\nu_n n / r_n}}, \qquad B_n = \frac{M_n^{ex} - \nu_n L_n^{ex}}{\sqrt{\nu_n n T_n}},$$
$$C_n = \frac{r_n}{\sqrt{n}} \left( L_n^{in} - \frac{n}{r_n} \right), \qquad D_n = \frac{r_n}{n} L_n^{ex} - r_n T_n + \log n + 1,$$

so that

$$M_n^{in} = \frac{\nu_n n}{r_n} + \sqrt{\frac{\nu_n n}{r_n}} A_n + \frac{\sqrt{n}\nu_n}{r_n} C_n, \tag{59}$$
$$M_n^{ex} = \nu_n n T_n - \frac{\nu_n n \log n}{r_n} - \frac{\nu_n n}{r_n} + \sqrt{\nu_n n T_n} B_n + \frac{\nu_n n}{r_n} D_n. \tag{60}$$

Corollaries 2 and 3 will follow from the next proposition, which says that $(A_n, B_n, C_n, D_n)$ are asymptotically independent.

**Proposition 11.** *Suppose the following two conditions hold:*

$$\lim_{n\to\infty} \frac{\log n}{r_n T_n} = 0, \qquad \lim_{n\to\infty} \frac{\nu_n n}{r_n} = \infty. \tag{61}$$

*Then as $n \to \infty$,*

$$(A_n, B_n, C_n, D_n) \Rightarrow (Z_1, Z_2, Z, \log W), \tag{62}$$

*where $Z$, $Z_1$, and $Z_2$ have a standard normal distribution, $W$ has an exponential distribution with mean one, and $Z$, $Z_1$, $Z_2$, and $W$ are independent. Furthermore, if the first condition in (61) is replaced by the weaker condition (13), then we still have the convergence*

$$(A_n, C_n) \Rightarrow (Z_1, Z). \tag{63}$$

To prove Proposition 11, we will need the following lemma.

**Lemma 12.** *Let* $(a_n)_{n=1}^{\infty}$, $(\delta_n)_{n=1}^{\infty}$, *and* $(\epsilon_n)_{n=1}^{\infty}$ *be three sequences such that*

$$\lim_{n\to\infty} a_n = \infty, \qquad \lim_{n\to\infty} \frac{\delta_n}{a_n} = 0, \qquad \lim_{n\to\infty} \frac{\epsilon_n}{a_n} = 0.$$

*Then for any* $M > 0$ *and* $t \in \mathbb{R}$, *we have as* $n \to \infty$,

$$\sup_{-M\le c\le M} \left| \exp\left\{ (a_n + \delta_n + c\epsilon_n) \left( \exp\left(\frac{it}{\sqrt{a_n}}\right) - \frac{it}{\sqrt{a_n}} - 1 \right) \right\} - \exp\left(-\frac{t^2}{2}\right) \right| \to 0.$$

*Proof.* Using the inequality (see Lemma 3.3.19 of[73])

$$\left| e^{ix} - \sum_{m=0}^{n} \frac{(ix)^m}{m!} \right| \le \frac{|x|^{n+1}}{(n+1)!}, \tag{64}$$

we have

$$\begin{aligned} &\sup_{-M\le c\le M} \left| (a_n + \delta_n + c\epsilon_n) \left( \exp\left(\frac{it}{\sqrt{a_n}}\right) - \frac{it}{\sqrt{a_n}} - 1 \right) + \frac{t^2}{2} \right| \\ &\quad\le \sup_{-M\le c\le M} \left( \left| (a_n + \delta_n + c\epsilon_n)\left(-\frac{t^2}{2a_n}\right) + \frac{t^2}{2} \right| + |a_n + \delta_n + c\epsilon_n| \cdot \frac{1}{6} \left| \frac{it}{\sqrt{a_n}} \right|^3 \right) \\ &\quad\le \frac{t^2(|\delta_n| + M|\epsilon_n|)}{2a_n} + \frac{t^3}{6} \left( \frac{|a_n| + |\delta_n| + M|\epsilon_n|}{a_n^{3/2}} \right), \end{aligned}$$

which goes to 0 as $n \to \infty$. The result follows because the exponential function is continuous. □

*Proof of Proposition 11.* We first prove (62) under the condition (61). We apply Theorem 3 of[74] where the marginal distributions are the distributions of $(C_n, D_n)$ and the conditional distributions are the conditional distributions of $(A_n, B_n)$ given $(C_n, D_n)$. We know from Theorem 1 that $(C_n, D_n) \Rightarrow (Z, \log W)$. For $t = (t_1, t_2) \in \mathbb{R}^2$ and $(c, d) \in \mathbb{R}^2$, let

$$\phi_n(t, (c,d)) = \mathbb{E}\left[ e^{it_1 A_n + it_2 B_n} | C_n = c, D_n = d \right], \quad \phi_0(t, (c,d)) = \mathbb{E}\left[ e^{it_1 Z_1 + it_2 Z_2} \right].$$

Note that $\phi_0(t, (c,d))$ does not depend on $(c, d)$. We need to check the following:

1. For all compact sets $I \subset \mathbb{R}^2$, we have

$$\sup_{(c,d)\in I} |\phi_n(t, (c,d)) - \phi_0(t, (c,d))| \to 0 \text{ as } n \to \infty.$$

2. The function $\phi_0(t, (c,d))$ is equicontinuous in $(c, d)$ at $t = 0$.

3. For each $t$, the function $\phi_0(t, (c,d))$ is a continuous function of $(c, d)$.

The last two conditions are trivially satisfied because $\phi_0(t, (c,d))$ is constant as a function of $(c, d)$.

Now we check the first condition. Conditional on $C_n = c$ and $D_n = d$, which means

$$L_n^{in} = \frac{n}{r_n} + \frac{c\sqrt{n}}{r_n}, \qquad L_n^{ex} = nT_n - \frac{n\log n}{r_n} - \frac{n}{r_n} + \frac{dn}{r_n},$$

the random variables $(M_n^{in}, M_n^{ex})$ are independent Poisson with means $\nu_n L_n^{in}$ and $\nu_n L_n^{ex}$ respectively. Therefore, we have

$$\begin{aligned}\phi_n(t,(c,d)) =& \exp\left\{\left(\frac{\nu_n n}{r_n} + \frac{c\nu_n\sqrt{n}}{r_n}\right)\left(\exp\left(it_1\sqrt{\frac{r_n}{\nu_n n}}\right) - it_1\sqrt{\frac{r_n}{\nu_n n}} - 1\right)\right\} \\ &\times \exp\left\{\left(\nu_n nT_n - \frac{\nu_n n\log n}{r_n} - \frac{\nu_n n}{r_n} + \frac{d\nu_n n}{r_n}\right)\left(\exp\left(\frac{it_2}{\sqrt{\nu_n nT_n}}\right) - \frac{it_2}{\sqrt{\nu_n nT_n}} - 1\right)\right\}\end{aligned}$$

and

$$\phi_0(t,(c,d)) = \exp\left(-\frac{t_1^2+t_2^2}{2}\right).$$

Note that if $|z_i| \le 1$ and $|w_i| \le 1$ for $i = 1, 2$, then $|z_1 z_2 - w_1 w_2| \le |z_1 - w_1| + |z_2 - w_2|$. It suffices to check that for all $M > 0$, as $n \to \infty$, we have

$$\sup_{-M\le c\le M}\left|\exp\left\{\left(\frac{\nu_n n}{r_n} + \frac{c\nu_n\sqrt{n}}{r_n}\right)\left(\exp\left(it_1\sqrt{\frac{r_n}{\nu_n n}}\right) - it_1\sqrt{\frac{r_n}{\nu_n n}} - 1\right)\right\} - \exp\left(-\frac{t_1^2}{2}\right)\right| \to 0. \tag{65}$$

and

$$\begin{aligned}\sup_{-M\le d\le M}\Bigg|\exp\Bigg\{&\left(\nu_n nT_n - \frac{\nu_n n\log n}{r_n} - \frac{\nu_n n}{r_n} + \frac{d\nu_n n}{r_n}\right) \\ &\times\left(\exp\left(\frac{it_2}{\sqrt{\nu_n nT_n}}\right) - \frac{it_2}{\sqrt{\nu_n nT_n}} - 1\right)\Bigg\} - \exp\left(-\frac{t_2^2}{2}\right)\Bigg| \to 0.\end{aligned} \tag{66}$$

Equations (65) and (66) follow from Lemma 12 and the assumptions in (61), which implies that (62) holds under the condition (61).

To prove (63) when the first condition in (61) is replaced by the weaker condition (13), we repeat the above argument using only $A_n$ and $C_n$. It is then necessary to check only (65), which does not require the first condition in (61). □

*Proof of Corollary 2.* Note that, when we define $\sigma_n^2 = n(\nu_n/r_n + \nu_n^2/r_n^2)$ as in Corollary 2, we have

$$\frac{1}{\sigma_n}\left(M_n^{in} - \frac{n\nu_n}{r_n}\right) = \frac{1}{\sigma_n}\left(\sqrt{\frac{\nu_n n}{r_n}}A_n + \frac{\sqrt{n}\nu_n}{r_n}C_n\right).$$

Proposition 11 implies that the distribution of the right-hand side converges to the standard normal distribution as $n \to \infty$, which is the conclusion of Corollary 2. □

*Proof of Corollary 3.* From (59) and (60), we get

$$\hat{T}_n = \frac{M_n^{ex}}{r_n M_n^{in}} + \frac{\log n + 1 + \gamma}{r_n}$$
$$= \frac{n\nu_n T_n - n\nu_n(\log n)/r_n - n\nu_n/r_n + \sqrt{n\nu_n T_n}B_n + n\nu_n D_n/r_n}{n\nu_n + \sqrt{n\nu_n r_n}A_n + \sqrt{n}\nu_n C_n} + \frac{\log n + 1 + \gamma}{r_n}$$
$$= \frac{T_n - (\log n)/r_n - 1/r_n + \sqrt{T_n/(n\nu_n)}B_n + D_n/r_n}{1 + \sqrt{r_n/(n\nu_n)}A_n + C_n/\sqrt{n}} + \frac{\log n + 1 + \gamma}{r_n}. \tag{67}$$

For a sequence of random variables $(X_n)_{n=1}^{\infty}$ and a sequence of real numbers $(a_n)_{n=1}^{\infty}$, we write $X_n = o_p(a_n)$ if $X_n/a_n$ converges to 0 in probability as $n \to \infty$. Since we are assuming (20), it follows that

$$\sqrt{r_n/(n\nu_n)}A_n = o_p(1), \qquad C_n/\sqrt{n} = o_p(1), \qquad (\log n + 1)/r_n = o_p(T_n)$$

$$\sqrt{T_n/(n\nu_n)}B_n = T_n\sqrt{1/(r_nT_n)}\sqrt{r_n/n\nu_n}B_n = o_p(T_n), \qquad D_n/r_n = o_p(T_n).$$

Therefore, writing $E_n$ for an error term which is $o_p\big(\sqrt{r_n/(n\nu_n)}A_n + C_n/\sqrt{n}\big)$, we can rewrite (67) as

$$\hat{T}_n = \Big(T_n - (\log n)/r_n - 1/r_n + \sqrt{T_n/(n\nu_n)}B_n + D_n/r_n\Big)$$
$$\times \Big(1 - \sqrt{r_n/(n\nu_n)}A_n - C_n/\sqrt{n} + E_n\Big) + \frac{\log n + 1 + \gamma}{r_n}$$
$$= T_n - \sqrt{r_n/(n\nu_n)}T_nA_n + \sqrt{T_n/(n\nu_n)}B_n - T_nC_n/\sqrt{n} + \frac{D_n + \gamma}{r_n}$$
$$+ o_p\Big(T_n\big(\sqrt{r_n/(n\nu_n)}A_n + C_n/\sqrt{n}\big)\Big). \tag{68}$$

Now we do a variance computation by replacing $A_n$, $B_n$, and $C_n$ with $Z_1$, $Z_2$, and $Z$ respectively. We have

$$\mathrm{Var}\big(-\sqrt{r_n/(n\nu_n)}T_nZ_1 + \sqrt{T_n/(n\nu_n)}Z_2 - T_nZ/\sqrt{n}\big)$$
$$= \frac{r_nT_n^2}{n\nu_n} + \frac{T_n}{n\nu_n} + \frac{T_n^2}{n} = \frac{T_n}{n\nu_n}(r_nT_n + \nu_nT_n + 1) \sim \frac{(r_n + \nu_n)T_n^2}{n\nu_n}, \tag{69}$$

where $\sim$ means that the ratio of the two sides tends to 1 as $n \to \infty$. Equation (69) and Proposition 11 imply that the distribution of

$$\frac{1}{T_n}\sqrt{\frac{n\nu_n}{r_n + \nu_n}}\left(-\sqrt{r_n/(n\nu_n)}T_nA_n + \sqrt{T_n/(n\nu_n)}B_n - T_nC_n/\sqrt{n}\right)$$

converges to the standard normal distribution. It follows from (20) and Proposition 11 that

$$\frac{1}{T_n}\sqrt{\frac{n\nu_n}{r_n + \nu_n}} \cdot \frac{D_n + \gamma}{r_n} \Rightarrow 0.$$

Thus, the conclusion of Corollary 3 follows from (68). □

### 2 Supplementary Information – Derivation of Confidence Intervals

We explain in this section how our confidence intervals were derived.

#### 2.1 Confidence interval based on internal branch lengths

Recall that we estimate the growth rate $r_n$ from the internal branch lengths by using

$$\hat{r}_n = \frac{n}{L_n^{in}}.$$

Let $0 < \alpha < 1$, and recall that $z_{\alpha/2}$ is the number such that if $Z$ has a standard normal distribution, then $P(Z > z_{\alpha/2}) = \alpha/2$. By Theorem 1, we have

$$\begin{aligned} 1 - \alpha &= \lim_{n\to\infty} \mathbb{P}\bigg( - z_{\alpha/2} \leq \frac{r_n}{\sqrt{n}}\Big(L_n^{in} - \frac{n}{r_n}\Big) \leq z_{\alpha/2}\bigg) \\ &= \lim_{n\to\infty} \mathbb{P}\bigg(\frac{n}{r_n}\Big(1 - \frac{z_{\alpha/2}}{\sqrt{n}}\Big) \leq L_n^{in} \leq \frac{n}{r_n}\Big(1 + \frac{z_{\alpha/2}}{\sqrt{n}}\Big)\bigg). \end{aligned}$$

Multiplying the inequality by $r_n/L_n^{in}$, we get

$$\begin{aligned} 1 - \alpha &= \lim_{n\to\infty} \mathbb{P}\bigg(\frac{n}{L_n^{in}}\Big(1 - \frac{z_{\alpha/2}}{\sqrt{n}}\Big) \leq r_n \leq \frac{n}{L_n^{in}}\Big(1 + \frac{z_{\alpha/2}}{\sqrt{n}}\Big)\bigg) \\ &= \lim_{n\to\infty} \mathbb{P}\bigg(\hat{r}_n\Big(1 - \frac{z_{\alpha/2}}{\sqrt{n}}\Big) \leq r_n \leq \hat{r}_n\Big(1 + \frac{z_{\alpha/2}}{\sqrt{n}}\Big)\bigg). \end{aligned}$$

Therefore, an asymptotically valid $100(1-\alpha)\%$ confidence interval for $r_n$ is given by

$$\bigg[\hat{r}_n\Big(1 - \frac{z_{\alpha/2}}{\sqrt{n}}\Big), \hat{r}_n\Big(1 + \frac{z_{\alpha/2}}{\sqrt{n}}\Big)\bigg].$$

#### 2.2 Confidence interval based on shared mutations

Recall that, if the mutation rate $\nu_n$ and the site frequency spectrum are known, but we do not have a reconstruction of the entire tree, we can estimate the growth rate from the number of shared mutations using

$$\hat{r}_n = \frac{n\nu_n}{M_n^{in}}. \tag{70}$$

Recall that in the statement of Corollary 2, we defined

$$\sigma_n^2 = \frac{n\nu_n^2}{r_n^2}\Big(1 + \frac{r_n}{\nu_n}\Big).$$

From Corollary 2, we get

$$
\begin{aligned}
1-\alpha &= \lim_{n\to\infty} \mathbb{P}\left( -z_{\alpha/2} \le \frac{1}{\sigma_n}\left(M_n^{in} - \frac{n\nu_n}{r_n}\right) \le z_{\alpha/2}\right) \\
&= \lim_{n\to\infty} \mathbb{P}\left( \frac{n\nu_n}{r_n} - z_{\alpha/2}\sigma_n \le M_n^{in} \le \frac{n\nu_n}{r_n} + z_{\alpha/2}\sigma_n\right) \\
&= \lim_{n\to\infty} \mathbb{P}\left( \frac{n\nu_n}{r_n}\left(1 - \frac{z_{\alpha/2}}{\sqrt{n}}\sqrt{1+\frac{r_n}{\nu_n}}\right) \le M_n^{in} \le \frac{n\nu_n}{r_n}\left(1 + \frac{z_{\alpha/2}}{\sqrt{n}}\sqrt{1+\frac{r_n}{\nu_n}}\right)\right).
\end{aligned}
$$

We now multiply the inequality by $r_n/M_n$ to get

$$
1-\alpha = \lim_{n\to\infty} \mathbb{P}\left( \hat{r}_n\left(1 - \frac{z_{\alpha/2}}{\sqrt{n}}\sqrt{1+\frac{r_n}{\nu_n}}\right) \le r_n \le \hat{r}_n\left(1 + \frac{z_{\alpha/2}}{\sqrt{n}}\sqrt{1+\frac{r_n}{\nu_n}}\right)\right).
$$

Note that we can not use these upper and lower bounds as a confidence interval because they involve $r_n$, which is unknown. However, we can approximate $r_n/\nu_n$ under the square root by $\hat{r}_n/\nu_n$, which equals $n/M_n^{in}$. Because $\hat{r}_n/r_n$ converges in probability to 1 when (18) holds, an asymptotically valid $100(1-\alpha)\%$ confidence interval for $r_n$ is given by

$$
\left[\hat{r}_n\left(1 - \frac{z_{\alpha/2}}{\sqrt{n}}\sqrt{1+\frac{n}{M_n^{in}}}\right),\ \hat{r}_n\left(1 + \frac{z_{\alpha/2}}{\sqrt{n}}\sqrt{1+\frac{n}{M_n^{in}}}\right)\right].
$$

#### 2.3 Confidence interval for maximum likelihood

We assume here that we have random variables $H_i = a_n + b_n U_i$, where $U_1, \ldots, U_{n-1}$ are i.i.d. and have the standard logistic distribution. A package in R can then be used to obtain a maximum likelihood estimate $\hat{b}_n$ for $b_n$. It is known (see[40]) that when $n$ is large, the estimate $\hat{b}_n$ is asymptotically normal, and the variance of $\hat{b}_n$ can be estimated using the Cramer-Rao bound, which gives

$$
\mathrm{Var}(\hat{b}_n) \sim \frac{9}{3+\pi^2} \cdot \frac{b_n^2}{n}.
$$

Therefore, letting $c = 3/\sqrt{3+\pi^2}$, we have

$$
\begin{aligned}
1-\alpha &= \lim_{n\to\infty} \mathbb{P}\left( b_n - z_{\alpha/2}\cdot\frac{cb_n}{\sqrt{n}} \le \hat{b}_n \le b_n + z_{\alpha/2}\cdot\frac{cb_n}{\sqrt{n}}\right) \\
&= \lim_{n\to\infty} \mathbb{P}\left( b_n\left(1 - z_{\alpha/2}\cdot\frac{c}{\sqrt{n}}\right) \le \hat{b}_n \le b_n\left(1 + z_{\alpha/2}\cdot\frac{c}{\sqrt{n}}\right)\right).
\end{aligned}
$$

We now divide the inequality by $b_n\hat{b}_n$. Recalling that $r_n = 1/b_n$ and that the estimator for $r_n$ can be expressed as $\hat{r}_n = 1/\hat{b}_n$, we arrive at

$$
1-\alpha = \lim_{n\to\infty} \mathbb{P}\left( \hat{r}_n\left(1 - z_{\alpha/2}\cdot\frac{c}{\sqrt{n}}\right) \le r_n \le \hat{r}_n\left(1 + z_{\alpha/2}\cdot\frac{c}{\sqrt{n}}\right)\right).
$$

This leads to our confidence interval for $r_n$, which is

$$\left[\hat{r}_n\left(1-\frac{cz_{\alpha/2}}{\sqrt{n}}\right),\ \hat{r}_n\left(1+\frac{cz_{\alpha/2}}{\sqrt{n}}\right)\right].$$

When we calculate this confidence interval from data, the random variables $H_i$ that are used to obtain the estimate $\hat{r}_n$ only approximately have a logistic distribution. We have not proved the asymptotic validity of the confidence interval using this approximation, but the confidence interval performs well in simulations.

#### 2.4 Confidence interval for tumor age

Recall that when $r_n$ is known, we can estimate the tumor age $T_n$ by $\hat{T}_n$. By Corollary 3, when (20) holds, we have

$$\frac{1}{T_n}\sqrt{\frac{n\nu_n}{r_n+\nu_n}}\left(\hat{T}_n-T_n\right)\Rightarrow Z,$$

where $Z$ has a standard normal distribution. Therefore,

$$\begin{aligned}1-\alpha &= \lim_{n\to\infty}\mathbb{P}\left(-z_{\alpha/2}\le\frac{1}{T_n}\sqrt{\frac{n\nu_n}{r_n+\nu_n}}\left(\hat{T}_n-T_n\right)\le z_{\alpha/2}\right)\\ &= \lim_{n\to\infty}\mathbb{P}\left(T_n\left(1-z_{\alpha/2}\sqrt{\frac{r_n+\nu_n}{n\nu_n}}\right)\le\hat{T}_n\le T_n\left(1+z_{\alpha/2}\sqrt{\frac{r_n+\nu_n}{n\nu_n}}\right)\right)\\ &= \lim_{n\to\infty}\mathbb{P}\left(\frac{\hat{T}_n}{1+z_{\alpha/2}\sqrt{\frac{r_n+\nu_n}{n\nu_n}}}\le T_n\le\frac{\hat{T}_n}{1-z_{\alpha/2}\sqrt{\frac{r_n+\nu_n}{n\nu_n}}}\right).\end{aligned}$$

The mutation rate $\nu_n$ is not assumed here to be known, but by rearranging (70) and using the result of Corollary 2, we see that $\nu_n$ can be estimated by

$$\hat{\nu}_n=\frac{r_nM_n^{in}}{n},$$

and then $\hat{\nu}_n/\nu_n$ converges in probability to 1. When we replace $\nu_n$ by $\hat{\nu}_n$, the square root in the expression above becomes

$$\sqrt{\frac{r_n+\hat{\nu}_n}{n\hat{\nu}_n}}=\frac{1}{\sqrt{n}}\sqrt{1+\frac{n}{M_n^{in}}}.$$

It follows that for $0<\alpha<1$, an asymptotically valid $100(1-\alpha)\%$ confidence interval for $T_n$ can be obtained by

$$\left[\frac{\hat{T}_n}{1+\frac{z_{\alpha/2}}{\sqrt{n}}\sqrt{1+\frac{n}{M_n^{in}}}},\ \frac{\hat{T}_n}{1-\frac{z_{\alpha/2}}{\sqrt{n}}\sqrt{1+\frac{n}{M_n^{in}}}}\right].$$

### 3 Supplementary Information - Data application

#### 3.1 Agreement across methods for growth rate estimates

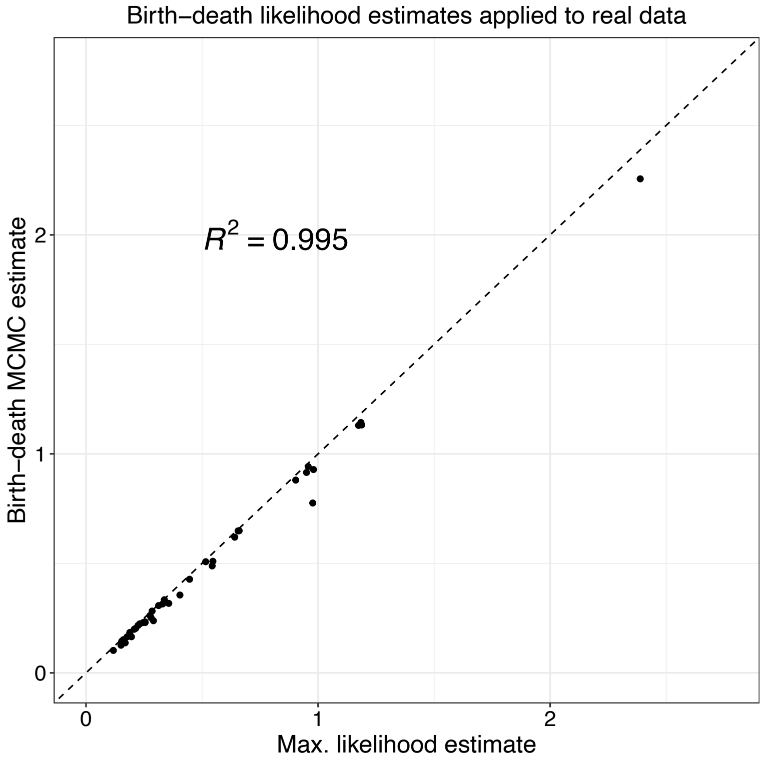


Supplementary Figure 1: **Maximum likelihood and birth-death MCMC agree on blood data:** There are a few cases where the maximum likelihood estimate is slightly higher than the birth-death MCMC, but the differences are generally very small. The agreement indicates that our assumptions of $n << N, T \to \infty$, and $n \to \infty$ do not significantly affect the estimates when applied to the data.

We see excellent agreement between the birth-death MCMC and our maximum likelihood estimates when applied to real data (Supp. Fig. 1). This motivated our decision to include only the maximum likelihood estimates in Application to human blood datasets for ease of illustration. The birth-death MCMC agreement with our maximum likelihood estimates provides validation that the approximations required to use the maximum likelihood estimate did not affect estimates in any significant way, at least in the context of this dataset.

While agreement between the maximum likelihood and internal lengths estimates is generally good when applied to the real data (Supp. Fig. 2), the internal lengths estimate tends to be higher than the maximum likelihood estimate in many cases. We hypothesize that this may be due to slight differences in the fitness of cells within the clone. While, on average, clones fit the neutral expectation (see Fig. 5A), there may be cases where the random merging of lineages is violated to some degree. Slight fitness differences within a clone, leading to non-random lineage merging, could reduce the sum of internal lengths, leading to higher estimates of $r$, based on Eq. 4. In order to test this, we re-generated trees by randomly merging the lineages using the coalescence times from the real data. In these re-generated trees (see Supp. Fig. 2B), the internal lengths estimates agreed more closely with the maximum likelihood estimates, suggesting that slight differences in fitness within a clone may be responsible for the discrepancies that we see in the real data sets.

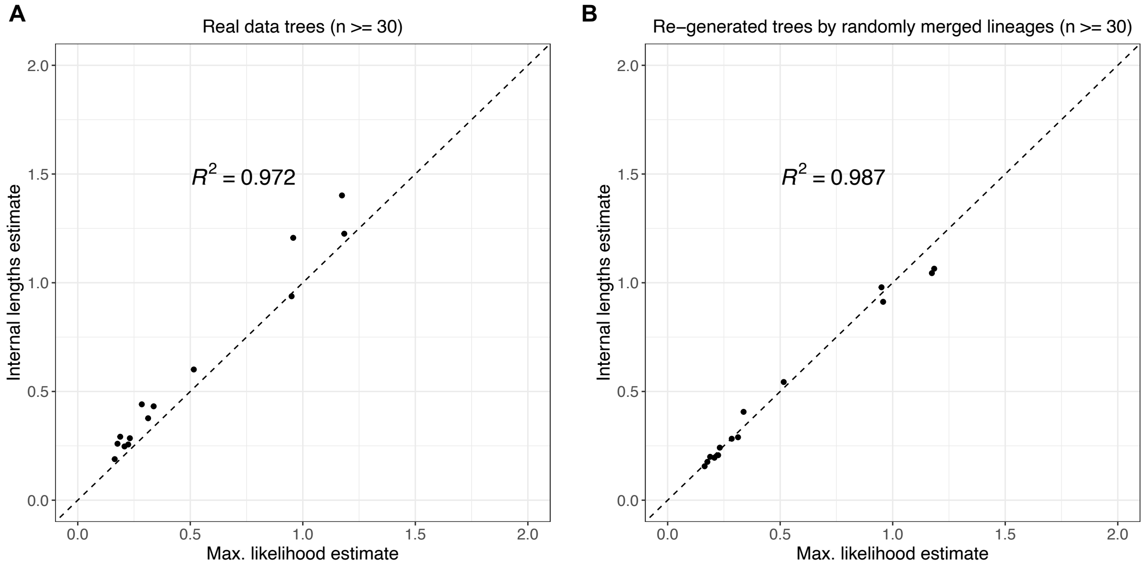


Supplementary Figure 2: **Maximum likelihood and internal lengths estimate. A:** Internal lengths estimates for some clones are higher than maximum likelihood estimates. **B:** When we re-generate trees by taking the coalescence times from the data and randomly merging the lineages, we see better agreement between internal lengths and maximum likelihood estimates. Note: we use only those real data trees with $n \geq 30$ sampled cells in this analysis, to ensure that the effect is less likely to be caused by chance.

#### 3.2 Aberrant Cell Fraction

Williams et al. define a quantity for aberrant cell fraction (ACF) as the number of cells within a specific expanding clone divided by the total number of cells sequenced[22]. ACF is an approximation of the clonal fraction at the time of sampling estimated from single cell-derived colonies. ACF can be incorporated into the inference methods introduced by Williams et al., which include a Markov chain Monte Carlo (MCMC)-based method called Phylofit and a method that employs Approximate Bayesian computation (ABC). In simulations where the growth trajectory is logistic with a carrying capacity equal to a clonal fraction of 1, Phylofit with ACF outperforms Phylofit without ACF as well as our methods (see Supp. Fig. 3).

However, data[19,22,24] show that these Phylofit with ACF estimates are often discordant with early growth rate estimates. Williams et al. show that Phylofit produces significantly different estimates for growth rate depending on whether ACF is incorporated into the likelihood equation, suggesting that some clones show "smaller than expected final clonal fractions" (see Ext. Data Fig. 7c from Williams et al.[22]). Fabre et al.[19], who also include data from Mitchell et al.[24], compare their phylodynamic estimates using *phylodyn*[29] to the expected clone size later in life. Similar to Williams et al., they conclude that "at least for some clones and genes, the dynamics observed in later life are not representative of those that prevail earlier"[19]. Combining data from all three sources[19,22,24], we show in Supp. Fig. 4A that most clones do not reach the expected fraction that would be predicted by a

**A**

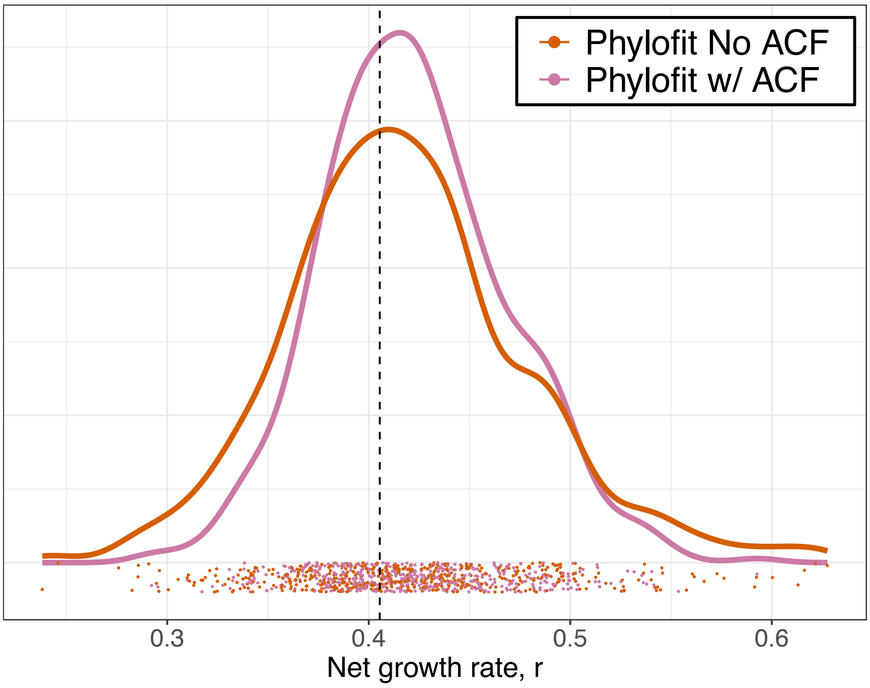


**B**

| Method | RMSE | Mean Estimate | SD Estimate | True Growth Rate |
|---|---|---|---|---|
| Phylofit w/ ACF | 0.049 | 0.423 | 0.046 | 0.405 |
| Max. likelihood | 0.060 | 0.423 | 0.057 | 0.405 |
| Phylofit No ACF | 0.062 | 0.418 | 0.060 | 0.405 |
| Internal lengths | 0.065 | 0.404 | 0.065 | 0.405 |

Supplementary Figure 3: **Incorporating aberrant cell fraction (ACF) leads to improved estimates on simulated data.** **A:** Estimated growth rates from Phylofit with (pink) and without (orange) ACF metric shows that ACF does give a slight advantage in the simulated case. Function *run_selection_sim()* from the *rsimpop* package[22] was used to simulate a logistic birth-death clonal expansion with growth rate $r = \log(1.5) = 0.405$ and a carrying capacity equal to a total HSC population size of $N = 100,000$. That is, we allowed the clone's carrying capacity to be equal to a clonal fraction (or ACF) of 1. The simulated clone was allowed to expand for 33 years following driver initiation, so that the expected population size was roughly 85% of the carrying capacity. $n = 100$ cells were then sampled to reconstruct the tree. The process was repeated 500 times. **B:** Table shows performance across methods for same simulations from (A), including our methods and ordered by root mean square error (RMSE).

logistic growth trajectory reaching a clonal fraction of 1. A similar analysis is shown in Figure 4d of Fabre et al.[19].

Our methods for estimating the net growth rate are derived from coalescent theory (specifically coalescent point processes), and estimate the growth rate during the early expansion phase of the clone. Specifically, because we estimate the growth rate from early coalescence times, our estimates reflect the growth rate during the time period when the number of cells in the clone is on the order of the sample size, $n$, which is typically between 10 and 100 in the data we have analyzed. Aberrant cell

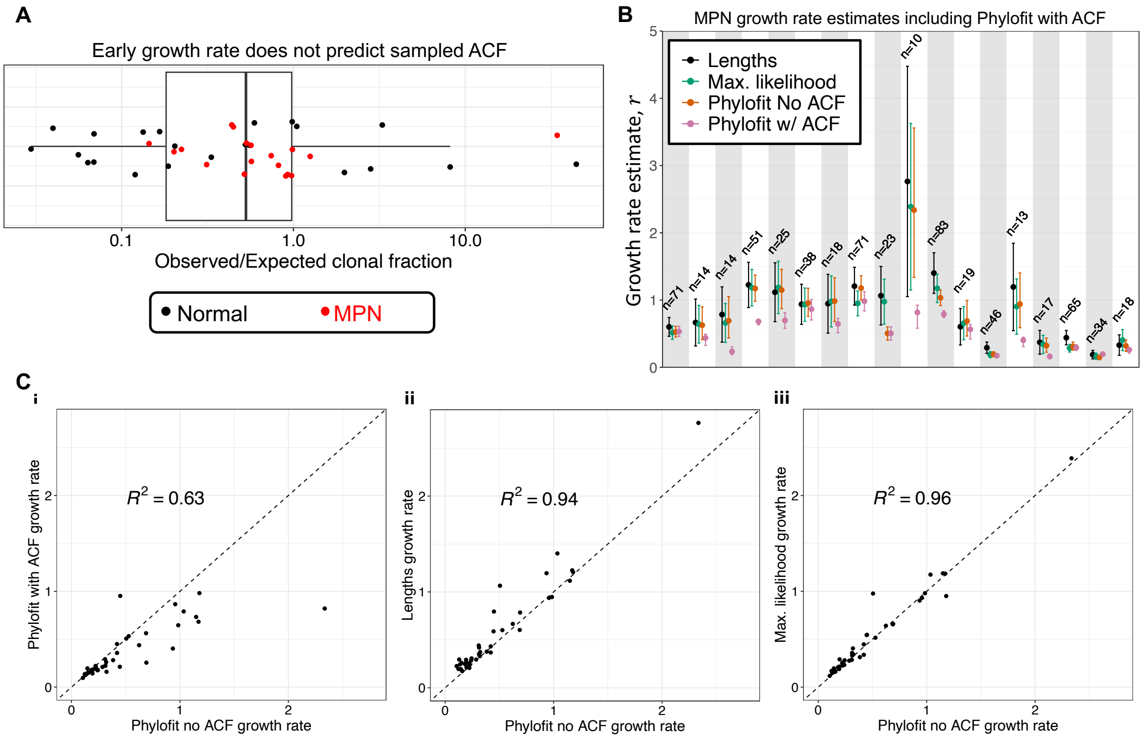


Supplementary Figure 4: **Early growth rate is not predicted by aberrant cell fraction (ACF). A:** Ratio of observed to expected clonal fraction assuming a logistic growth trajectory. There is a clear bias shown for clones which expand rapidly at first (high expected ACF) but decelerate (Observed ACF $<$ Expected ACF). Parameters used to calculate expected ACF are Phylofit (no ACF) growth rate and HSC carrying capacity $N = 100,000$. **B:** Growth rate estimates of clones from individuals with MPN shows that Phylofit with ACF is a low outlier in many cases. **C:** Correlation between Phylofit without ACF and Phylofit with ACF (i), internal lengths method (ii), and maximum likelihood method (iii) from estimates applied to hematopoietic clones. Phylofit with ACF is not capturing the same expansion rates estimated by other methods. Specifically, the correlation coefficient between Phylofit with ACF and Phylofit without ACF (i) is lowest of all the methods, with a clear bias towards lower growth rates estimated by the Phylofit with ACF method. Note that all hematopoietic clones included except the two clones from the Van Egeren dataset[23] are excluded from these Phylofit with ACF analyses because the cells were not sampled randomly.

fraction (ACF) estimates, on the other hand, will be affected by the growth rate from clone initiation all the way until the sampling time. Note that ACF estimates are, by definition, only relevant for detectable population sizes above a certain VAF threshold, when calculating the fraction of cells within a given clone is possible using bulk sequencing. Therefore, ACF estimates are possible when the number of cells in the clone is on the order of $N$, the total number of hematopoietic stem cells, which is estimated to be between 25,000 and 300,000[20,35,36]. Both Fabre et al.[19] and Williams et al.[22] observed slower growth rates and reduced ACF at later timepoints than would be expected based on the early expansion. Using our early growth rate estimates, we confirm these conclusions, demonstrating that incorporating ACF leads to reduced growth rate estimates that do not agree with those produced by our methods or those from Phylofit without ACF. In order to compare our estimates to the early growth rate estimates from other methods, such as Phylofit[22], it is necessary to exclude ACF from the likelihood calculation. Fortunately, Phylofit has been designed to run with or without ACF[22]. Once we remove ACF, we find similar estimates across methods (see Supp. Fig. 4B-C).

Possible explanations for the discordance between growth rates from early expansion and those using ACF are treatment effects, frequency dependent selection, and nested or independent clonal expansions which may outcompete the original clone. Clinical evidence suggests that some patients have relatively stable clonal fractions below 1 for extended periods of time[75]. Further, estimates of clonal fraction may be affected by sampled cell type (Peripheral blood vs. bone marrow and granulocytes vs. mononuclear cells vs. whole blood), as shown in Supp. Fig. 5. As noted previously, Van Egeren et al.[23] observe this in the context of *JAK2* mutant cells, which appear at higher frequency in erythroid progenitors. Therefore, the use of clonal fraction (or ACF) in the estimation of early growth rates using coalescence times is prone to errors due to measuring difficulties and poorly understood growth trajectories of clones at high fraction. While measures of clonal fraction are valuable clinically[75], there is no single growth rate metric that can currently be used to explain early coalescent and late clonal fraction data in most cases.

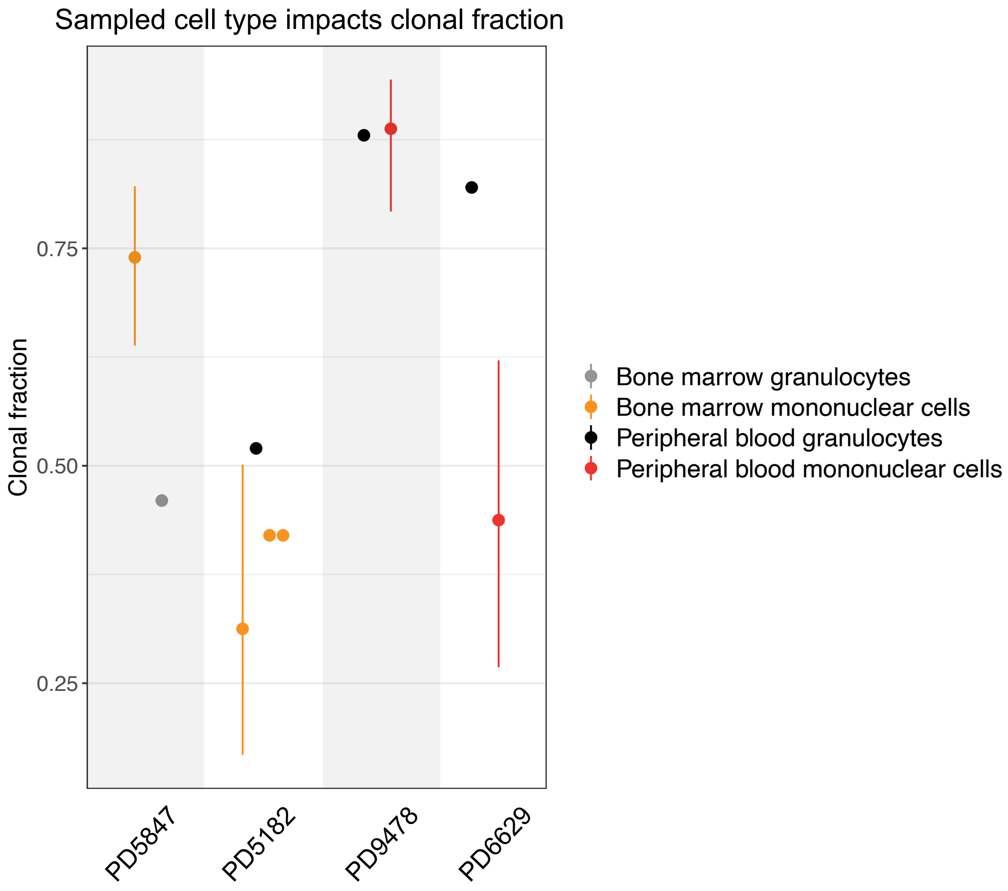


Supplementary Figure 5: **Clonal fraction estimates depend on sampled cell types.** In three out of four individuals with samples from different cell types taken within a month of each other, dominant clonal fraction is significantly different depending on the sampled cell type (all except PD9478). Estimates with error bars represent those taken from single cell colony data, with 95% confidence intervals based on binomial sampling. Points without error bars represent bulk recapture samples at mean depth $> 300\times$. All data from Williams et al.[22].

##### 3.3 Annotating Clones

As noted in the main text, "early mergers" are instances which will affect our methods significantly. For an example of an early merger occurring in a simulated tree, see Supp. Fig. 6A. Such an early merge in the ancestry of two sampled cells leads to a sum of internal lengths much greater than would otherwise be expected. Also, the distribution of coalescence times would now have an outlier. Thus, both the internal lengths method and the method of maximum likelihood would be affected, and the resulting growth rate estimate from each method would be lower than the true value.

In the context of hematopoiesis, there are several possible explanations for why an early merger might occur. First, there may be an expansion within a clone due to a cell within that clone acquiring an additional fitness advantage. We call this a "nested expansion". In Supp. Fig. 6B, we see that the *DNMT3A* clone has two such expansions, one resulting from a *CBL* mutation, and the other resulting from a *JAK2* mutation. While these expansions are a result of known driver mutations and have more sampled cells, it is possible for a recent or slow growing expansion to only be captured by two sampled cells. Further, it is possible for the expansion to be caused by an unknown driver mutation or epigenetic change, complicating the annotation of a such a nested expansion.

Second, the early merger can be a result of chance. While unlikely, the sampling may capture two cells which have a recent common ancestor. If stem cell turnover independent of further clonal expansions is high or the clone population size is low, sampled cells are more likely to have recent common ancestors. This probability will be influenced by the number of total hematopoietic stem cells (HSCs) as well as the normal turnover rate. Fewer stem cells and higher turnover increase the probability of early mergers, consistent with coalescent theory in critical branching processes[61].

In both of these cases, estimating the main or parent clone will be more accurate if the early merger is removed from the phylogenetic tree. In the dataset we analyze, there were no clear instances of early mergers without annotated drivers. That is, the case of Supp. Fig. 6A did not occur. However, many cases similar to Supp. Fig. 6B occur, where a larger nested clone or one with a known driver is present. In those cases, we remove the nested expansion(s) and estimate the growth rate using the remaining samples. If the nested expansion has 10 or more samples, we also estimate the growth rate of the nested clone independently.

Supp. Fig. 6B shows an example of manual annotation of clones. Most of this work was performed in the papers that generated the data[19,22–24]. However, there are cases where our approach differs slightly from the annotation in those works. For example, as shown in Supp. Fig. 6B, we include an expanded clone with an unknown driver which is not included in the work of Williams et al.[22]. Other expanded clones without known drivers are annotated by the authors in the work of Fabre et al.[19] and Mitchell et al.[24] and are not present in the data from Van Egeren et al.[23], so they do not lead to differences in clonal annotation between our work and these three papers. Further, in cases where nested expansions occur, we remove the expansions from the tree that is ultimately used for our estimates, leaving only one resulting tip (sampled cell) from each nested expansion, so as to preserve the original coalescence event preceding the acquisition of an added fitness advantage. It is unclear whether this step is performed in the previous analyses[19,22,24]. Nested expansions do not appear in the data from Van Egeren et al.[23]. Because we use the same tree and clone annotations when running Phylofit and our methods, these differences do not affect the consistency across methods. Many sources are available for tree reconstruction and further work will be required to automate the process of annotating clones and identifying nested clones. Such work will likely require metrics of tree balance in addition to coalescent theory. For the time being, manual annotation of clones is the best option, and we attempted to use the most accurate trees as input.

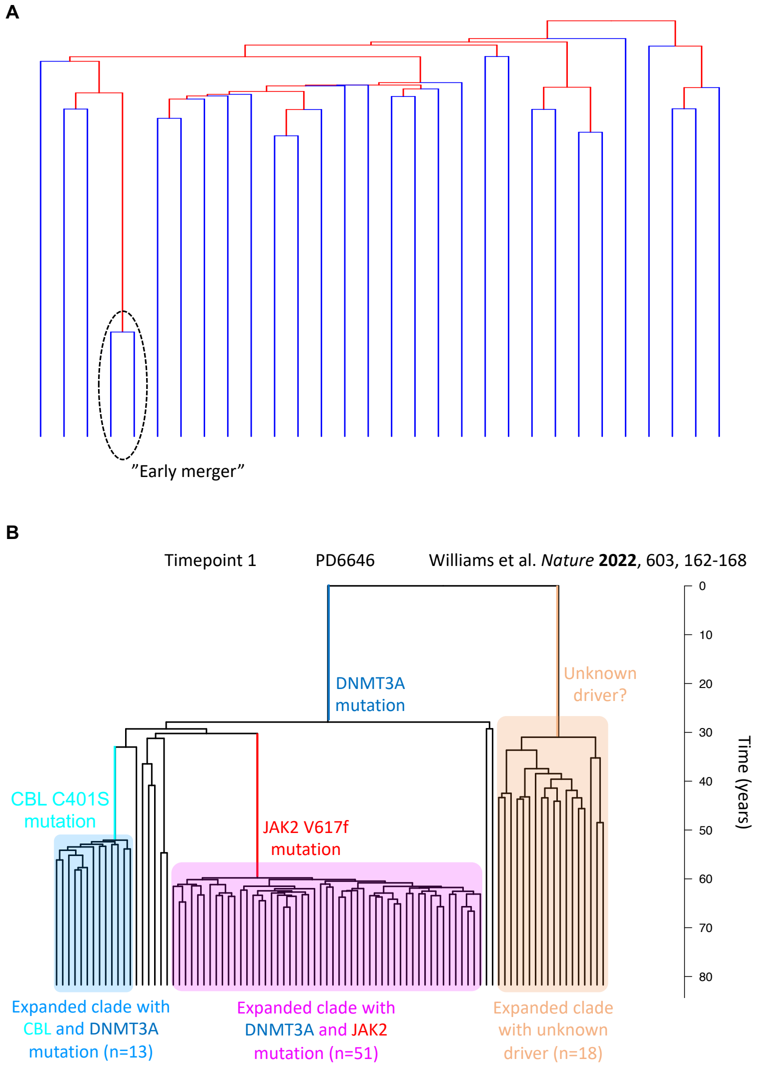


Supplementary Figure 6: **Example trees. A:** An example of an early merger which would make our methods inaccurate. Excess internal lengths which is a result of a possible nested expansion and/or normal stem cell turnover would lead to a smaller growth rate than would otherwise be estimated without such an early merger. Similarly, maximum likelihood estimate using the distribution of coalescence times would lead to a smaller growth rate. If the merger is caused by a known driver or thought to be a nested expansion it can be removed. However, it may be difficult to decide whether to remove the early merger in cases where there is no clear driver. **B:** A real data example of a reconstructed phylogenetic tree from Williams et al.[22] with clades with greater than 10 sampled cells annotated. Each tip of the tree represents a sampled cell. Here, we see a *DNMT3A* mutation leading to nested clones, one with a *CBL* mutation and one with a *JAK2* mutation. This tree also shows a clonal expansion with an unknown driver, and we estimated a growth rate for this clone.

#### 3.4 Supplementary Tables for blood data applications

| Patient + Timepoint | Patient age (yr) | Age at diagnosis (yr) | Number of cells in clone, n | Number of total sampled cells | Driver | Ext. to Int. lengths ratio | Max. likelihood estimate | Internal lengths estimate | Birth-death MCMC estimate | Phylofit (no ACF) estimate | Phylofit (with ACF) estimate | Clone age (Max. likelihood estimate, yr) |
|---|---|---|---|---|---|---|---|---|---|---|---|---|
| PD4781 timepoint 1 | 84.5 | 73 | 38 | 39 | JAK2:p.V617F and 9pUPD and TET2:p.Q1632* and 7p- and 7q+ | 17.3 | 0.95 (0.7 - 1.2) | 0.94 (0.64 - 1.24) | 0.92 (0.68 - 1.18) | 0.96 (0.76 - 1.18) | 0.87 (0.7 - 1.01) | 23.8 |
| PD4781 timepoint 2 | 86.8 | 73 | 18 | 22 | JAK2:p.V617F and 9pUPD and TET2:p.Q1632* and 7p- and 7q+ | 20.2 | 0.98 (0.6 - 1.36) | 0.95 (0.51 - 1.38) | 0.93 (0.6 - 1.35) | 0.99 (0.68 - 1.33) | 0.65 (0.51 - 0.73) | 26.0 |
| PD5117 timepoint 1 | 82.7 | 64 | 46 | 88 | JAK2:p.V617F | 16.0 | 0.19 (0.14 - 0.23) | 0.29 (0.21 - 0.38) | 0.18 (0.14 - 0.23) | 0.19 (0.16 - 0.23) | 0.17 (0.14 - 0.2) | 77.8 |
| PD5147 timepoint 1 | 91.7 | 81 | 65 | 66 | PPM1D:p.T483fs*3 and TET2:p.S657fs*42 | 21.6 | 0.28 (0.23 - 0.34) | 0.44 (0.33 - 0.55) | 0.28 (0.23 - 0.34) | 0.31 (0.26 - 0.38) | 0.29 (0.25 - 0.34) | 62.8 |
| PD5179 timepoint 1 | 50.4 | 34 | 83 | 92 | JAK2:p.V617F and 9+ and 9q- and 1q+ | 25.8 | 1.17 (0.96 - 1.39) | 1.4 (1.1 - 1.7) | 1.13 (0.94 - 1.35) | 1.03 (0.92 - 1.15) | 0.79 (0.74 - 0.84) | 24.3 |
| PD5182 timepoint 1 | 33.5 | 32 | 10 | 32 | JAK2:p.V617F and 9pUPD | 42.1 | 2.39 (1.15 - 3.63) | 2.76 (1.05 - 4.48) | 2.26 (1.25 - 3.63) | 2.34 (1.34 - 3.56) | 0.82 (0.58 - 0.92) | 16.8 |
| PD5182 timepoint 3 | 54.1 | 32 | 23 | 93 | JAK2:p.V617F and 9pUPD and 1q+ | 15.5 | 0.98 (0.64 - 1.31) | 1.06 (0.63 - 1.5) | 0.78 (0.52 - 1.11) | 0.51 (0.4 - 0.6) | 0.5 (0.41 - 0.6) | 24.2 |
| PD5847 timepoint 1 | 45.6 | 44 | 71 | 96 | JAK2:p.V617F and 9pUPD and TET2:p.N281fs*1 | 12.8 | 0.96 (0.77 - 1.14) | 1.21 (0.93 - 1.49) | 0.94 (0.76 - 1.14) | 1.18 (1 - 1.36) | 0.98 (0.84 - 1.12) | 15.5 |
| PD6629 timepoint 1 | 61.0 | 54 | 17 | 32 | DNMT3A:p.R882H | 16.4 | 0.34 (0.21 - 0.48) | 0.37 (0.2 - 0.55) | 0.32 (0.21 - 0.47) | 0.32 (0.23 - 0.43) | 0.16 (0.13 - 0.2) | 56.9 |
| PD6629 timepoint 1 | 61.0 | 54 | 14 | 32 | DNMT3A:p.R882H and JAK2:p.V617F | 17.6 | 0.64 (0.36 - 0.92) | 0.67 (0.32 - 1.02) | 0.62 (0.39 - 0.92) | 0.63 (0.41 - 0.9) | 0.44 (0.33 - 0.49) | 32.5 |
| PD6629 timepoint 2 | 63.0 | 54 | 14 | 27 | DNMT3A:p.R882H and JAK2:p.V617F | 22.9 | 0.66 (0.37 - 0.95) | 0.79 (0.37 - 1.2) | 0.65 (0.4 - 0.99) | 0.69 (0.44 - 1.05) | 0.23 (0.19 - 0.3) | 34.7 |
| PD6634 timepoint 1 | 60.3 | 26 | 34 | 36 | PPM1D:p.Q462* | 4.7 | 0.16 (0.12 - 0.21) | 0.19 (0.13 - 0.25) | 0.15 (0.1 - 0.2) | 0.15 (0.12 - 0.18) | 0.2 (0.17 - 0.22) | 60.2 |
| PD6646 timepoint 1 | 81.7 | 76 | 13 | 90 | DNMT3A:p.? and CBL:p.C401S | 31.9 | 0.9 (0.49 - 1.31) | 1.2 (0.55 - 1.84) | 0.88 (0.53 - 1.36) | 0.94 (0.59 - 1.4) | 0.4 (0.31 - 0.45) | 30.8 |
| PD6646 timepoint 1 | 81.7 | 76 | 51 | 90 | DNMT3A:p.? and JAK2:p.V617F | 22.3 | 1.18 (0.91 - 1.46) | 1.23 (0.89 - 1.56) | 1.14 (0.9 - 1.42) | 1.17 (0.98 - 1.37) | 0.68 (0.63 - 0.72) | 22.8 |
| PD6646 timepoint 1 | 81.7 | 76 | 18 | 90 | unknown | 12.6 | 0.4 (0.25 - 0.56) | 0.33 (0.18 - 0.48) | 0.36 (0.23 - 0.51) | 0.32 (0.24 - 0.41) | 0.26 (0.21 - 0.29) | 53.2 |
| PD6646 timepoint 2 | 85.4 | 76 | 25 | 27 | DNMT3A:p.? and JAK2:p.V617F | 25.0 | 1.19 (0.8 - 1.58) | 1.12 (0.68 - 1.55) | 1.13 (0.8 - 1.53) | 1.15 (0.87 - 1.46) | 0.7 (0.58 - 0.81) | 26.5 |
| PD7271 timepoint 1 | 24.0 | 20 | 19 | 88 | JAK2:p.V617F | 10.5 | 0.66 (0.41 - 0.9) | 0.6 (0.33 - 0.87) | 0.65 (0.43 - 0.93) | 0.69 (0.47 - 0.99) | 0.56 (0.42 - 0.64) | 23.3 |
| PD9478 timepoint 1 | 68.8 | 53 | 71 | 80 | JAK2:p.F537_K539delinsL and DNMT3A:p.Y908* | 9.8 | 0.52 (0.42 - 0.62) | 0.6 (0.46 - 0.74) | 0.51 (0.41 - 0.61) | 0.53 (0.45 - 0.61) | 0.53 (0.46 - 0.61) | 25.4 |

**Supplementary Table 1: Model results for data from Williams et al.**[22]

| Patient ID | Patient age (yr) | Age at diagnosis (yr) | Number of cells in clone, n | Driver | Ext. to Int. lengths ratio | Max. Likelihood estimate | Internal lengths estimate | Birth-death MCMC estimate | Phylofit (no ACF) estimate | Published estimate | Clone age (max. likelihood estimate, yr) |
|---|---|---|---|---|---|---|---|---|---|---|---|
| vanEgerenET1 | 34 | 34 | 22 | JAK2:p.V617F | 14.8 | 0.55 (0.36 - 0.74) | 0.8 (0.46 - 1.13) | 0.51 (0.35 - 0.71) | 0.45 (0.34 - 0.57) | 0.49 (0.39 - 0.58) | 26.8 |
| vanEgerenET2 | 63 | 63 | 13 | JAK2:p.V617F | 12.1 | 0.45 (0.24 - 0.65) | 0.37 (0.17 - 0.57) | 0.43 (0.26 - 0.65) | 0.42 (0.27 - 0.61) | 0.36 (0.27 - 0.45) | 42.2 |

**Supplementary Table 2: Model results for data from Van Egeren et al.**[23] Note: Phylofit with ACF cannot be calculated for these clones because cells were not randomly sampled.

| Patient | Patient age (yr) | Number of cells in clone, n | Number of total sampled cells | Driver | Ext. to Int. lengths ratio | Max. likelihood estimate | Internal lengths estimate | Birth-death MCMC estimate | Phylofit (no ACF) estimate | Phylofit (with ACF) estimate | Clone age (Max. likelihood estimate, yr) |
|---|---|---|---|---|---|---|---|---|---|---|---|
| PD34493 | 83 | 15 | 92 | SF3B1:k666n and DelY | 4.7 | 0.17 (0.1 - 0.24) | 0.2 (0.1 - 0.3) | 0.14 (0.07 - 0.21) | 0.12 (0.08 - 0.16) | 0.14 (0.11 - 0.17) | 54.4 |
| PD34493 | 83 | 48 | 92 | unknown | 11.8 | 0.23 (0.18 - 0.29) | 0.28 (0.2 - 0.37) | 0.22 (0.18 - 0.28) | 0.24 (0.2 - 0.28) | 0.18 (0.14 - 0.21) | 64.1 |
| PD34493 | 83 | 10 | 92 | unknown | 7.9 | 0.2 (0.09 - 0.3) | 0.19 (0.07 - 0.31) | 0.16 (0.09 - 0.27) | 0.14 (0.09 - 0.21) | 0.14 (0.09 - 0.2) | 65.4 |
| PD41276 | 79 | 78 | 84 | SF3B1:k666n | 9.9 | 0.34 (0.27 - 0.4) | 0.43 (0.34 - 0.53) | 0.33 (0.27 - 0.4) | 0.42 (0.33 - 0.54) | 0.36 (0.31 - 0.41) | 35.4 |
| PD41305 | 73 | 12 | 91 | unknown | 17.8 | 0.33 (0.17 - 0.49) | 0.42 (0.18 - 0.66) | 0.31 (0.19 - 0.48) | 0.31 (0.2 - 0.46) | 0.23 (0.16 - 0.26) | 54.0 |
| PD41305 | 73 | 15 | 91 | unknown | 13.1 | 0.28 (0.16 - 0.39) | 0.29 (0.15 - 0.44) | 0.26 (0.16 - 0.39) | 0.29 (0.19 - 0.41) | 0.21 (0.15 - 0.24) | 60.0 |
| PD41305 | 73 | 17 | 91 | unknown | 29.2 | 0.54 (0.33 - 0.76) | 0.59 (0.31 - 0.87) | 0.49 (0.31 - 0.71) | 0.45 (0.33 - 0.58) | 0.21 (0.18 - 0.23) | 59.3 |

**Supplementary Table 3: Model results for data from Fabre et al.**[19]

| Patient | Patient age (yr) | Number of cells in clone, n | Number of total sampled cells | Driver | Ext. to Int. lengths ratio | Max. likelihood estimate | Internal lengths estimate | Birth-death MCMC estimate | Phylofit (no ACF) estimate | Phylofit (with ACF) estimate | Clone age (Max. likelihood estimate, yr) |
|---|---|---|---|---|---|---|---|---|---|---|---|
| KX003 | 81 | 109 | 328 | unknown | 12.4 | 0.31 (0.26 - 0.36) | 0.38 (0.31 - 0.45) | 0.31 (0.26 - 0.36) | 0.38 (0.33 - 0.44) | 0.28 (0.26 - 0.29) | 51.7 |
| KX003 | 81 | 10 | 328 | unknown | 6.5 | 0.16 (0.08 - 0.24) | 0.17 (0.07 - 0.28) | 0.15 (0.08 - 0.25) | 0.16 (0.1 - 0.24) | 0.15 (0.09 - 0.21) | 61.6 |
| KX004 | 78 | 76 | 451 | DNMT3A | 10.6 | 0.21 (0.17 - 0.25) | 0.25 (0.19 - 0.3) | 0.2 (0.16 - 0.24) | 0.18 (0.16 - 0.2) | 0.18 (0.16 - 0.19) | 76.5 |
| KX004 | 78 | 27 | 451 | DNMT3A | 10.0 | 0.25 (0.17 - 0.32) | 0.24 (0.15 - 0.33) | 0.23 (0.16 - 0.31) | 0.22 (0.17 - 0.27) | 0.19 (0.16 - 0.21) | 64.3 |
| KX004 | 78 | 16 | 451 | unknown | 8.3 | 0.25 (0.15 - 0.36) | 0.24 (0.12 - 0.36) | 0.23 (0.15 - 0.34) | 0.23 (0.16 - 0.3) | 0.21 (0.15 - 0.24) | 55.3 |
| KX004 | 78 | 11 | 451 | DNMT3A | 12.1 | 0.21 (0.11 - 0.32) | 0.27 (0.11 - 0.42) | 0.2 (0.12 - 0.31) | 0.19 (0.12 - 0.29) | 0.16 (0.11 - 0.19) | 62.9 |
| KX004 | 78 | 13 | 451 | unknown | 12.8 | 0.36 (0.19 - 0.52) | 0.35 (0.16 - 0.54) | 0.32 (0.19 - 0.49) | 0.31 (0.21 - 0.42) | 0.22 (0.17 - 0.25) | 51.1 |
| KX004 | 78 | 23 | 451 | DNMT3A | 10.1 | 0.15 (0.1 - 0.21) | 0.24 (0.14 - 0.34) | 0.14 (0.1 - 0.2) | 0.14 (0.1 - 0.18) | 0.14 (0.1 - 0.17) | 69.4 |
| KX004 | 78 | 46 | 451 | unknown | 7.3 | 0.22 (0.17 - 0.28) | 0.26 (0.18 - 0.33) | 0.22 (0.17 - 0.27) | 0.22 (0.18 - 0.27) | 0.22 (0.18 - 0.26) | 49.4 |
| KX004 | 78 | 12 | 451 | unknown | 8.6 | 0.12 (0.06 - 0.17) | 0.23 (0.1 - 0.35) | 0.1 (0.05 - 0.16) | 0.11 (0.07 - 0.15) | 0.1 (0.06 - 0.14) | 71.8 |
| KX008 | 76 | 53 | 367 | unknown | 7.4 | 0.18 (0.14 - 0.22) | 0.26 (0.19 - 0.33) | 0.16 (0.13 - 0.21) | 0.15 (0.13 - 0.17) | 0.15 (0.13 - 0.18) | 62.5 |
| KX008 | 76 | 29 | 367 | unknown | 6.8 | 0.15 (0.1 - 0.2) | 0.29 (0.19 - 0.4) | 0.13 (0.08 - 0.17) | 0.13 (0.1 - 0.16) | 0.13 (0.1 - 0.16) | 60.2 |
| KX008 | 76 | 14 | 367 | unknown | 14.2 | 0.28 (0.16 - 0.4) | 0.31 (0.15 - 0.47) | 0.25 (0.16 - 0.38) | 0.24 (0.17 - 0.33) | 0.18 (0.14 - 0.2) | 63.9 |
| KX008 | 76 | 15 | 367 | unknown | 6.8 | 0.25 (0.15 - 0.36) | 0.21 (0.1 - 0.31) | 0.23 (0.15 - 0.34) | 0.22 (0.15 - 0.3) | 0.21 (0.15 - 0.25) | 52.7 |
| KX008 | 76 | 15 | 367 | unknown | 8.7 | 0.29 (0.17 - 0.41) | 0.21 (0.11 - 0.32) | 0.24 (0.15 - 0.36) | 0.2 (0.14 - 0.25) | 0.18 (0.14 - 0.21) | 64.9 |

**Supplementary Table 4: Model results for data from Mitchell et al.**[24]

#### 3.5 Supplementary Table of site frequency information

| ID | Driver | n | k = 2 | k = 3 | k = 4 | k = 5 | k = 6 | k = 7 | k = 8 | k = 9 | k > 9 |
|---|---|---|---|---|---|---|---|---|---|---|---|
| Neutral Expectation 1/(k(k-1)) | NA | NA | 0.5000 | 0.1667 | 0.0833 | 0.0500 | 0.0333 | 0.0238 | 0.0179 | 0.0139 | 0.1101 |
| Mean Upper C.I. | NA | NA | 0.5204 | 0.2029 | 0.0998 | 0.0845 | 0.0664 | 0.0539 | 0.0386 | 0.0215 | 0.1264 |
| Aggregated Mean | NA | NA | 0.4723 | 0.1727 | 0.0814 | 0.0633 | 0.0460 | 0.0344 | 0.0209 | 0.0148 | 0.1015 |
| Mean Lower C.I. | NA | NA | 0.4242 | 0.1425 | 0.0629 | 0.0420 | 0.0256 | 0.0149 | 0.0032 | 0.0081 | 0.0767 |
| PD34493 | SF3B1 | 15 | 0.2642 | 0.0951 | 0.0951 | 0.0000 | 0.2059 | 0.3202 | 0.0194 | 0.0000 | 0.0000 |
| PD34493 | unknown | 48 | 0.5134 | 0.1393 | 0.0505 | 0.0421 | 0.0062 | 0.0362 | 0.0133 | 0.0078 | 0.1912 |
| PD34493 | unknown | 10 | 0.2939 | 0.2810 | 0.0493 | 0.0665 | 0.3094 | 0.0000 | 0.0000 | 0.0000 | NA |
| PD41305 | unknown | 12 | 0.4075 | 0.2819 | 0.0930 | 0.1300 | 0.0000 | 0.0000 | 0.0000 | 0.0346 | 0.0530 |
| PD41305 | unknown | 15 | 0.5187 | 0.2679 | 0.0630 | 0.0000 | 0.0000 | 0.0642 | 0.0237 | 0.0000 | 0.0626 |
| PD41305 | unknown | 17 | 0.4237 | 0.1239 | 0.0723 | 0.1232 | 0.0596 | 0.0000 | 0.0000 | 0.0000 | 0.1973 |
| PD41276 | SF3B1 | 78 | 0.5051 | 0.2109 | 0.1366 | 0.0270 | 0.0320 | 0.0000 | 0.0078 | 0.0079 | 0.0728 |
| KX003 | unknown | 109 | 0.5727 | 0.1373 | 0.0803 | 0.0304 | 0.0292 | 0.0186 | 0.0122 | 0.0046 | 0.1149 |
| KX003 | unknown | 10 | 0.2999 | 0.5485 | 0.0127 | 0.0554 | 0.0000 | 0.0836 | 0.0000 | 0.0000 | NA |
| KX004 | DNMT3A | 76 | 0.4825 | 0.1538 | 0.0622 | 0.0242 | 0.0458 | 0.0214 | 0.0355 | 0.0185 | 0.1561 |
| KX004 | DNMT3A | 27 | 0.5155 | 0.1465 | 0.0491 | 0.0327 | 0.0612 | 0.0843 | 0.0262 | 0.0000 | 0.0846 |
| KX004 | unknown | 16 | 0.3904 | 0.1856 | 0.0713 | 0.2513 | 0.0142 | 0.0000 | 0.0000 | 0.0000 | 0.0871 |
| KX004 | DNMT3A | 11 | 0.4735 | 0.2684 | 0.0000 | 0.0000 | 0.1972 | 0.0420 | 0.0000 | 0.0000 | 0.0190 |
| KX004 | unknown | 13 | 0.5700 | 0.0975 | 0.1218 | 0.0464 | 0.0000 | 0.0060 | 0.0000 | 0.0000 | 0.1583 |
| KX004 | DNMT3A | 23 | 0.5347 | 0.1198 | 0.0056 | 0.0439 | 0.0616 | 0.0000 | 0.0000 | 0.0894 | 0.1451 |
| KX004 | unknown | 46 | 0.5612 | 0.1235 | 0.0824 | 0.0549 | 0.0000 | 0.0099 | 0.0348 | 0.0223 | 0.1110 |
| KX004 | unknown | 12 | 0.1871 | 0.2032 | 0.1123 | 0.0000 | 0.0000 | 0.0357 | 0.3390 | 0.0384 | 0.0843 |
| KX008 | unknown | 53 | 0.4165 | 0.1522 | 0.0287 | 0.0828 | 0.0550 | 0.0133 | 0.0072 | 0.0638 | 0.1805 |
| KX008 | unknown | 29 | 0.3572 | 0.1629 | 0.0492 | 0.0594 | 0.0000 | 0.0145 | 0.0000 | 0.0308 | 0.3261 |
| KX008 | unknown | 14 | 0.2418 | 0.1355 | 0.3125 | 0.0154 | 0.0360 | 0.2587 | 0.0000 | 0.0000 | 0.0000 |
| KX008 | unknown | 15 | 0.6576 | 0.1035 | 0.1012 | 0.0174 | 0.0114 | 0.0000 | 0.0606 | 0.0000 | 0.0482 |
| KX008 | unknown | 15 | 0.3832 | 0.0462 | 0.1407 | 0.2805 | 0.0000 | 0.0000 | 0.0000 | 0.0000 | 0.1494 |
| PD7271_1 | JAK2 | 19 | 0.8203 | 0.0597 | 0.0750 | 0.0000 | 0.0046 | 0.0076 | 0.0000 | 0.0000 | 0.0329 |
| PD5182_1 | JAK2 and 9pUPD and 9+ | 10 | 0.5782 | 0.1709 | 0.0000 | 0.1095 | 0.0858 | 0.0557 | 0.0000 | 0.0000 | NA |
| PD5182_3 | JAK2 and 9pUPD and 1q+ and 9+ | 23 | 0.1196 | 0.2148 | 0.0655 | 0.2299 | 0.0000 | 0.0240 | 0.0000 | 0.0000 | 0.3462 |
| PD5847_1 | JAK2 and TET2 and 9pUPD and 9+ | 71 | 0.5843 | 0.1930 | 0.1139 | 0.0131 | 0.0105 | 0.0201 | 0.0029 | 0.0041 | 0.0581 |
| PD5179_1 | JAK2 and 1q+ and 9+ and 9q- | 83 | 0.3761 | 0.1718 | 0.0774 | 0.0934 | 0.0336 | 0.0443 | 0.0173 | 0.0270 | 0.1592 |
| PD6634_1 | PPM1D | 34 | 0.5100 | 0.1311 | 0.0579 | 0.0963 | 0.0000 | 0.0000 | 0.0000 | 0.0487 | 0.1561 |
| PD9478_1 | JAK2 and DNMT3A and 9+ | 71 | 0.5191 | 0.2072 | 0.0710 | 0.0180 | 0.0551 | 0.0013 | 0.0231 | 0.0304 | 0.0748 |
| PD6629_1 | DNMT3A | 17 | 0.4360 | 0.0588 | 0.2022 | 0.1061 | 0.1285 | 0.0380 | 0.0000 | 0.0000 | 0.0304 |
| PD6629_1 | JAK2 and DNMT3A | 14 | 0.7292 | 0.0350 | 0.0163 | 0.1438 | 0.0000 | 0.0319 | 0.0328 | 0.0000 | 0.0110 |
| PD6629_2 | JAK2 and DNMT3A | 14 | 0.5986 | 0.1298 | 0.0986 | 0.0708 | 0.0376 | 0.0387 | 0.0000 | 0.0000 | 0.0258 |
| PD6646_1 | DNMT3A and CBL | 13 | 0.4840 | 0.2313 | 0.0943 | 0.0000 | 0.0627 | 0.0502 | 0.0000 | 0.0469 | 0.0305 |
| PD6646_1 | JAK2 and DNMT3A | 51 | 0.4680 | 0.1884 | 0.0757 | 0.0725 | 0.0750 | 0.0088 | 0.0000 | 0.0000 | 0.1115 |
| PD6646_1 | unknown | 18 | 0.3929 | 0.3738 | 0.0758 | 0.0000 | 0.0000 | 0.0000 | 0.0207 | 0.0187 | 0.1180 |
| PD6646_2 | JAK2 and DNMT3A | 25 | 0.4651 | 0.1615 | 0.1956 | 0.0843 | 0.0190 | 0.0000 | 0.0158 | 0.0131 | 0.0455 |
| PD5117_1 | JAK2 | 46 | 0.5177 | 0.1160 | 0.0149 | 0.1325 | 0.0156 | 0.0351 | 0.0088 | 0.0217 | 0.1376 |
| PD4781_1 | JAK2 and TET2 and 9pUPD and 9+ and 7p- and 7q+ | 38 | 0.5451 | 0.2498 | 0.0303 | 0.0067 | 0.0064 | 0.0442 | 0.0066 | 0.0112 | 0.0997 |
| PD4781_2 | JAK2 and TET2 and 9pUPD and 9+ and 7p- and 7q+ | 18 | 0.7003 | 0.0969 | 0.0699 | 0.0104 | 0.0000 | 0.0067 | 0.0000 | 0.0610 | 0.0548 |
| PD5147_1 | TET2 and PPM1D | 65 | 0.3894 | 0.1936 | 0.1626 | 0.0085 | 0.1276 | 0.0282 | 0.0125 | 0.0220 | 0.0556 |
| vanEgerenET1 | JAK2 | 22 | 0.2118 | 0.2858 | 0.0791 | 0.0545 | 0.0419 | 0.0000 | 0.1566 | 0.0000 | 0.1702 |
| vanEgerenET2 | JAK2 | 13 | 0.8208 | 0.0000 | 0.0527 | 0.0233 | 0.1028 | 0.0000 | 0.0004 | 0.0000 | 0.0000 |

**Supplementary Table 5: Neutral expectation is consistent with site frequency data from blood.** First row shows the neutral expectation found by Durrett[42]. Highlighted rows 2-4 show aggregated data across all clones. The confidence intervals for the aggregated data capture the neutral expectation. Remaining rows show frequency spectrum for each individual clone. Clones with $n = 10$ have NA listed for $k > 9$ because it is not plausible to determine which mutations occurred after the driver was acquired if the mutations are present in all sampled cells of the clone. CSV file available, see data availability.

##### 3.6 Supplementary Tables of simulation statistics

| Method | Number of samples, n | Net growth rate, r | Clone age, T | Number of simulated trees | Mean estimate | SD estimate | Median estimate | Normalized RMSE | 95% Coverage probability | Runtime (s) | Number of cores | Number of chains |
|---|---|---|---|---|---|---|---|---|---|---|---|---|
| Lengths | 10 | 0.5 | 40 | 500 | 0.638 | 0.260 | 0.578 | 0.588 | 0.960 | 0.002 +/- 0.003 | 1 | NA |
| Max. likelihood | 10 | 0.5 | 40 | 500 | 0.688 | 0.247 | 0.635 | 0.621 | 0.908 | 0.018 +/- 0.004 | 1 | NA |
| Phylofit | 10 | 0.5 | 40 | 500 | 0.616 | 0.232 | 0.558 | 0.518 | 0.824 | 22.525 +/- 6.456 | 3 | 3 |
| Birth-death MCMC | 10 | 0.5 | 40 | 500 | 0.620 | 0.216 | 0.572 | 0.493 | 0.890 | 31.347 +/- 10.193 | 1 | 1 |
| Lengths | 30 | 0.5 | 40 | 500 | 0.538 | 0.102 | 0.532 | 0.218 | 0.950 | 0.003 +/- 0.003 | 1 | NA |
| Max. likelihood | 30 | 0.5 | 40 | 500 | 0.561 | 0.088 | 0.556 | 0.214 | 0.952 | 0.009 +/- 0.009 | 1 | NA |
| Phylofit | 30 | 0.5 | 40 | 500 | 0.538 | 0.090 | 0.530 | 0.195 | 0.820 | 32.321 +/- 2.472 | 3 | 3 |
| Birth-death MCMC | 30 | 0.5 | 40 | 500 | 0.530 | 0.083 | 0.524 | 0.176 | 0.960 | 101.948 +/- 30.499 | 1 | 1 |
| Lengths | 50 | 0.5 | 40 | 500 | 0.521 | 0.073 | 0.510 | 0.151 | 0.974 | 0.005 +/- 0.003 | 1 | NA |
| Max. likelihood | 50 | 0.5 | 40 | 500 | 0.536 | 0.064 | 0.528 | 0.146 | 0.956 | 0.01 +/- 0.016 | 1 | NA |
| Phylofit | 50 | 0.5 | 40 | 500 | 0.522 | 0.069 | 0.514 | 0.145 | 0.804 | 58.766 +/- 4.849 | 3 | 3 |
| Birth-death MCMC | 50 | 0.5 | 40 | 500 | 0.514 | 0.061 | 0.506 | 0.125 | 0.954 | 195.15 +/- 72.69 | 1 | 1 |
| Lengths | 100 | 0.5 | 40 | 500 | 0.508 | 0.050 | 0.505 | 0.102 | 0.948 | 0.009 +/- 0.015 | 1 | NA |
| Max. likelihood | 100 | 0.5 | 40 | 500 | 0.521 | 0.042 | 0.520 | 0.093 | 0.948 | 0.008 +/- 0.003 | 1 | NA |
| Phylofit | 100 | 0.5 | 40 | 500 | 0.514 | 0.048 | 0.515 | 0.101 | 0.798 | 194.063 +/- 117.937 | 3 | 3 |
| Birth-death MCMC | 100 | 0.5 | 40 | 500 | 0.508 | 0.041 | 0.508 | 0.083 | 0.948 | 848.529 +/- 178.573 | 1 | 1 |
| Lengths | 200 | 0.5 | 40 | 500 | 0.504 | 0.037 | 0.503 | 0.075 | 0.948 | 0.019 +/- 0.005 | 1 | NA |
| Max. likelihood | 200 | 0.5 | 40 | 500 | 0.511 | 0.031 | 0.512 | 0.066 | 0.948 | 0.012 +/- 0.003 | 1 | NA |
| Phylofit | 200 | 0.5 | 40 | 500 | 0.506 | 0.037 | 0.510 | 0.075 | 0.722 | 394.321 +/- 35.829 | 3 | 3 |
| Birth-death MCMC | 200 | 0.5 | 40 | 500 | 0.503 | 0.031 | 0.504 | 0.062 | 0.948 | 1156.507 +/- 291.044 | 1 | 1 |
| Lengths | 500 | 0.5 | 40 | 500 | 0.504 | 0.023 | 0.502 | 0.047 | 0.944 | 0.057 +/- 0.03 | 1 | NA |
| Max. likelihood | 500 | 0.5 | 40 | 500 | 0.507 | 0.019 | 0.506 | 0.041 | 0.928 | 0.018 +/- 0.011 | 1 | NA |
| Phylofit | 500 | 0.5 | 40 | 500 | 0.506 | 0.023 | 0.505 | 0.047 | 0.728 | 1894.601 +/- 485.482 | 3 | 3 |
| Birth-death MCMC | 500 | 0.5 | 40 | 500 | 0.503 | 0.019 | 0.502 | 0.039 | 0.930 | 3603.017 +/- 651.681 | 1 | 1 |

**Supplementary Table 6: Simulation Results at various sample sizes, $n$.** Summary statistics for each method applied to 500 simulated trees with clone age $T$, growth rate $r$, and number of samples $n$ specified. Birth rate $\lambda$ was sampled from a uniform distribution on $[r, 1+r]$ and death rate $\mu = \lambda - r$. Normalized Root Mean Square Error (RMSE) is shown to account for differences in the actual growth rate. Normalized Root Mean Square Error sums the square of fractional error values instead of actual error values. CSV file available, see data availability.

| Method | Number of samples, n | Net growth rate, r | Clone age, T | Number of simulated trees | Mean estimate | SD estimate | Median estimate | Normalized RMSE | 95% Coverage probability | Runtime (s) | Number of cores | Number of chains |
|---|---|---|---|---|---|---|---|---|---|---|---|---|
| Lengths | 100 | 0.15 | 40 | 500 | 0.243 | 0.063 | 0.227 | 0.748 | 0.104 | 0.007 +/- 0.013 | 1 | NA |
| Max. likelihood | 100 | 0.15 | 40 | 500 | 0.251 | 0.073 | 0.230 | 0.831 | 0.034 | 0.006 +/- 0.003 | 1 | NA |
| Phylofit | 100 | 0.15 | 40 | 500 | 0.158 | 0.024 | 0.157 | 0.170 | 0.720 | 175.973 +/- 80.499 | 3 | 3 |
| Birth-death MCMC | 100 | 0.15 | 40 | 500 | 0.154 | 0.024 | 0.154 | 0.160 | 0.952 | 89.558 +/- 150.259 | 1 | 1 |
| Lengths | 100 | 0.20 | 40 | 500 | 0.243 | 0.048 | 0.233 | 0.320 | 0.698 | 0.008 +/- 0.004 | 1 | NA |
| Max. likelihood | 100 | 0.20 | 40 | 500 | 0.248 | 0.052 | 0.236 | 0.351 | 0.568 | 0.007 +/- 0.003 | 1 | NA |
| Phylofit | 100 | 0.20 | 40 | 500 | 0.208 | 0.026 | 0.207 | 0.134 | 0.734 | 197.479 +/- 88.093 | 3 | 3 |
| Birth-death MCMC | 100 | 0.20 | 40 | 500 | 0.203 | 0.022 | 0.202 | 0.110 | 0.968 | 121.125 +/- 147.098 | 1 | 1 |
| Lengths | 100 | 0.25 | 40 | 500 | 0.269 | 0.030 | 0.267 | 0.140 | 0.916 | 0.009 +/- 0.015 | 1 | NA |
| Max. likelihood | 100 | 0.25 | 40 | 500 | 0.275 | 0.029 | 0.270 | 0.152 | 0.844 | 0.007 +/- 0.002 | 1 | NA |
| Phylofit | 100 | 0.25 | 40 | 500 | 0.259 | 0.031 | 0.260 | 0.128 | 0.712 | 194.654 +/- 95.431 | 3 | 3 |
| Birth-death MCMC | 100 | 0.25 | 40 | 500 | 0.254 | 0.026 | 0.253 | 0.105 | 0.942 | 161.998 +/- 131.174 | 1 | 1 |
| Lengths | 100 | 0.50 | 40 | 500 | 0.508 | 0.050 | 0.505 | 0.102 | 0.948 | 0.009 +/- 0.015 | 1 | NA |
| Max. likelihood | 100 | 0.50 | 40 | 500 | 0.521 | 0.042 | 0.520 | 0.093 | 0.948 | 0.008 +/- 0.003 | 1 | NA |
| Phylofit | 100 | 0.50 | 40 | 500 | 0.514 | 0.048 | 0.515 | 0.101 | 0.798 | 194.063 +/- 117.937 | 3 | 3 |
| Birth-death MCMC | 100 | 0.50 | 40 | 500 | 0.508 | 0.041 | 0.508 | 0.083 | 0.948 | 848.529 +/- 178.573 | 1 | 1 |
| Lengths | 100 | 0.75 | 40 | 500 | 0.769 | 0.074 | 0.765 | 0.102 | 0.964 | 0.01 +/- 0.026 | 1 | NA |
| Max. likelihood | 100 | 0.75 | 40 | 500 | 0.783 | 0.068 | 0.781 | 0.101 | 0.936 | 0.008 +/- 0.001 | 1 | NA |
| Phylofit | 100 | 0.75 | 40 | 500 | 0.771 | 0.072 | 0.767 | 0.100 | 0.786 | 188.874 +/- 115.011 | 3 | 3 |
| Birth-death MCMC | 100 | 0.75 | 40 | 500 | 0.763 | 0.066 | 0.761 | 0.090 | 0.944 | 698.912 +/- 96.786 | 1 | 1 |
| Lengths | 100 | 1.00 | 40 | 500 | 1.021 | 0.107 | 1.014 | 0.109 | 0.926 | 0.009 +/- 0.015 | 1 | NA |
| Max. likelihood | 100 | 1.00 | 40 | 500 | 1.043 | 0.091 | 1.043 | 0.100 | 0.936 | 0.008 +/- 0.001 | 1 | NA |
| Phylofit | 100 | 1.00 | 40 | 500 | 1.030 | 0.098 | 1.031 | 0.102 | 0.748 | 193.766 +/- 114.867 | 3 | 3 |
| Birth-death MCMC | 100 | 1.00 | 40 | 500 | 1.015 | 0.088 | 1.015 | 0.089 | 0.946 | 894.859 +/- 101.631 | 1 | 1 |

**Supplementary Table 7: Simulation Results at various growth rates, $r$.** Summary statistics for each method applied to 500 simulated trees with clone age $T$, growth rate $r$, and number of samples $n$ specified. Birth rate $\lambda$ was sampled from a uniform distribution on $[r, 1 + r]$ and death rate $\mu = \lambda - r$. Normalized Root Mean Square Error (RMSE) is shown to account for differences in the actual growth rate. Normalized Root Mean Square Error sums the square of fractional error values instead of actual error values. CSV file available, see data availability.
